# Programmable Edge-to-Edge Assembly of RNA Nanostructures

**DOI:** 10.64898/2026.03.29.714238

**Authors:** Cody Geary, Mai P. Tran, Erik Poppleton, Alena Taskina, Kerstin Göpfrich

**Affiliations:** Biophysical Engineering Group, Heidelberg University, Center for Molecular Biology of Heidelberg University (ZMBH), Berliner Str. 45, 69120 Heidelberg, Germany; Biomolecular Mechanics, Max Planck Institute for Polymer Research, Ackermannweg 10, 55128 Mainz, Germany; Cluster of Excellence SynthImmune, Heidelberg University, Germany

## Abstract

Constructing complex three-dimensional RNA nanostructures requires precise molecular connectors for controlled self-assembly. Existing RNA–RNA connectors, such as kissing loops, are restricted to coaxial, end-to-end joining, limiting the range of accessible geometries. Here, we introduce the alpha kissing loop (alphaKL), a compact, sequence-programmable RNA connector that enables edge-to-edge helix association. The alphaKL combines a four-nucleotide kissing loop with minor- and major-groove triplex interactions that pre-organize an *α*-shaped conformation compatible with co-transcriptional folding. Embedded into RNA origami tiles, alphaKLs drive multivalent assembly into filaments and lattices, visualized at nanoscale resolution by atomic force microscopy. All-atom molecular dynamics simulations reveal that triplex occupancy at each sequence position controls the preferred inter-helical angle. Cooperative backbone contacts progressively rigidify the multi-alphaKL interface beyond what individual motifs achieve alone. By linking helices along their edges rather than their ends, the alphaKL expands the structural design space of programmable RNA nanostructures, unlocking previously inaccessible architectures and applications.

## Introduction

Programmable RNA nanostructures leverage predictable base pairing and modular tertiary motifs to enable the assembly of complex, novel folds.^1–3^ In practice, these architectures are realized through a limited set of connectors that define how helices can be joined into higher-order structures. Across this toolkit, a single geometric constraint dominates: helices meet end-to-end through coaxial stacking. Kissing loops (KLs),^4,5^ paranemic crossover interactions,^6–8^ and related KL connector strategies^9,10^ all rely on base pairing at helix termini followed by continuous stacking across the junction (Figure 1a). This mechanism couples recognition and stability to the helix end, restricting where connections can form and confining RNA assembly to architectures accessible through coaxial alignment.

**Figure 1:**
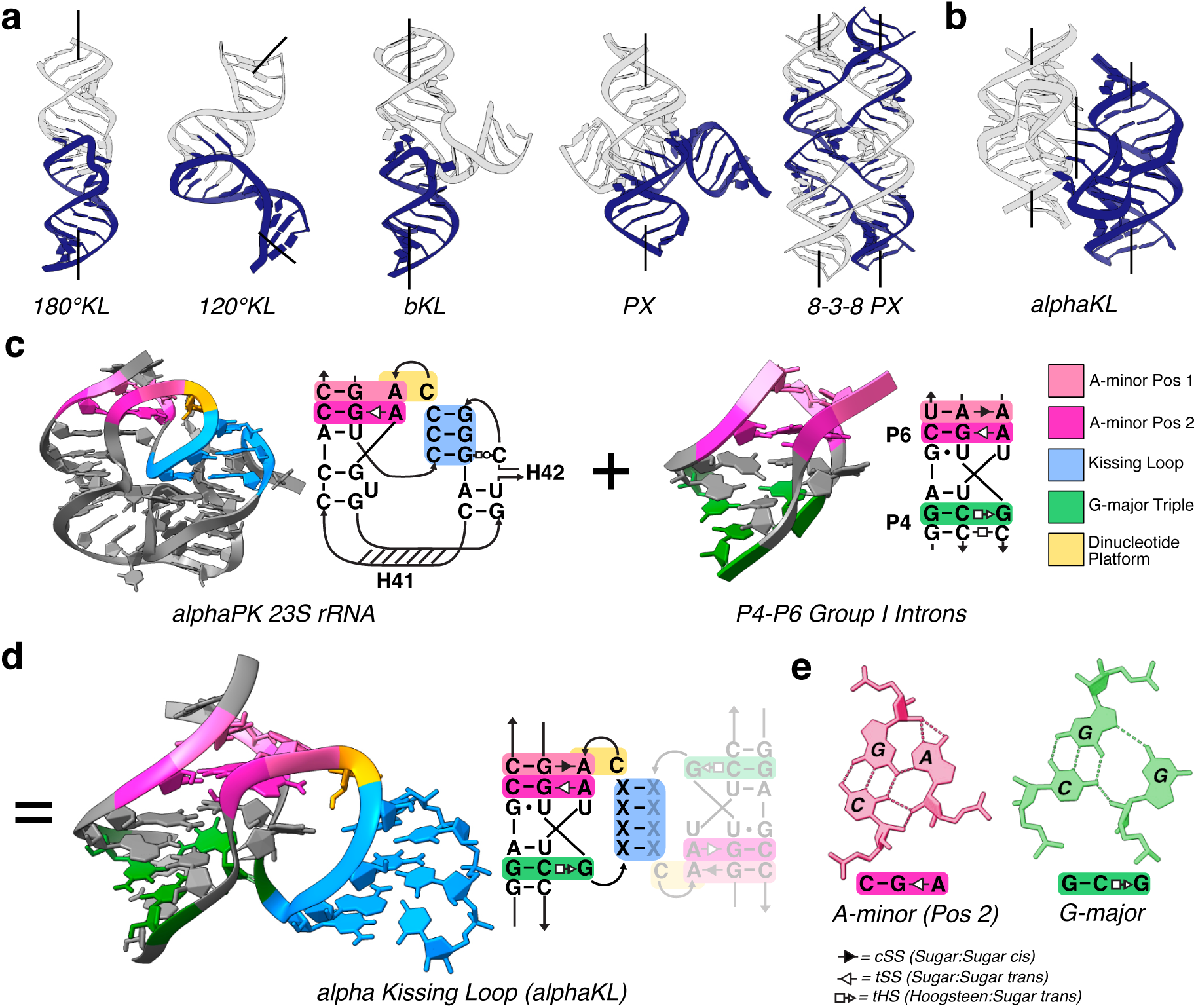
Design principle of the alphaKL and its structural features. **a)** Examples of KL architectures used for nanodesign. All examples have continuous coaxial stacking between light gray and dark blue double-stranded helix units. Left-to-right: 180° KL, 120° KL, branched KL, paranemic crossover (PX), 8-3-8 PX pattern. **b)** In contrast, the alphaKL is a new class of connector that links RNA helices edge-to-edge and forms no coaxial stacks between the white and dark blue units. **c)** Structure and diagram of alpha PK domain from 23S rRNA of *H. marismortui* (PDB ID 1S72) and P4-P6 junction from Group I Ribozyme of *Twort* (PDB ID 1Y0Q). **d)** Theoretical structure of alphaKL motif created by grafting fragments from alpha PK domain with P4-P6 junction **(c)**. **e)** Hydrogen bond patterns of A-minor triplex at position 2 (pos 2) and G-major groove triplex.

End-to-end coaxial stacking motifs dominate RNA nanostructure design, but linking helices along their edges poses a fundamentally different challenge while opening up an entirely new design space. During cotranscriptional folding, RNA structures form sequentially as they are synthesized by the polymerase,^11^ so any lateral connector must fold reliably within kinetically constrained pathways both *in vitro* and *in vivo*.^12–14^ A loop tethered along the side of a helix is flexible and can adopt multiple conformations, some of which partially stack with the parent helices. Even if these alternatives are transient, they can interfere with productive edge-to-edge association, creating kinetic bottlenecks. In the absence of coaxial stacking to stabilize the kissing interface, a successful connector must somehow pre-organize the loop to both avoid misfolding and provide sufficient rigidity to the resulting assembly. These considerations define the design requirements for a reliable edge-to-edge RNA connector.

A solution emerges from natural ribosomal RNA: the alpha-loop pseudoknot (alphaPK).^15,16^ In this motif, A-minor interactions anchor the loop to its parent helix, suggesting a mechanism by which the loop geometry could be pre-organized to favor productive long-range contacts. The geometric scaffold is highly conserved across both the 16S and 23S ribosomal subunits,^17^ whereas the sequence of the pseudoknotted loop can be reassigned.^18^ This combination of fixed geometry and sequence-programmable recognition defines the key requirements for a lateral RNA connector.

Here, we engineered these principles into a synthetic motif, the alpha kissing loop (alphaKL). By extending the natural alphaPK loop, introducing a complementary G-major triplex derived from group I ribozymes,^19^ and converting the intramolecular pseudoknot into a symmetric intermolecular interface, the alphaKL links helices edge-to-edge through coupled base pairing and groove-mediated contacts (Figure 1b). The motif preserves A-form helical geometry, is sequence-programmable, and folds cotranscriptionally within established RNA origami workflows.^11^ Integration into the ROAD^3^ and pyFuRNAce^20^ design platforms makes lateral helix association immediately accessible. By enabling helix–helix connectivity independent of coaxial stacking, the alphaKL expands the design space for RNA nanostructures beyond architectures defined by helix termini.

## Results

### Motif Design by Rational Engineering

Edge-to-edge connectivity was implemented by re-engineering the alphaPK motif^16^ as a synthetic, sequence-programmable dimer that mediates edge-to-edge helix association (Figure 1b). Using a modular recombination design strategy,^1,2^ the natural alphaPK topology was adapted to support complementary loop-loop pairing and major-groove triplex stabilization. When incorporated into cotranscriptionally folding RNA origami tiles,^11^ alphaKL connectors enable nanoscale architectures inaccessible to conventional KLs.

In the native ribosomal pseudoknot,^15^ an adenosine A-minor interaction precedes a sharp backbone turn into a three-nucleotide KL, producing a characteristic “*α*-shaped” topology^16^ (Figure 1c). Strict phylogenetic conservation of the A-minor packing geometry (Figure S1) highlights its structural importance to the ribosome, while mutational analysis shows that sequence variation of the pseudoknotted loop is tolerated.^18^ This conserved geometry, combined with sequence variability, provided a rational starting point for engineering a programmable edge-to-edge interface.

To convert this intramolecular fold into a stable intermolecular connector, the KL was extended from three to four nucleotides to increase stability and permit stable self-complementary pairing (blue, Figure 1e–f). Loop elongation required symmetrical lengthening of the 5’-strand entering the major groove. To stabilize the expanded interface, an additional guanine was introduced to form a major-groove triplex contact (Figure S2) analogous to those observed in group I ribozymes.^19^ Five-nucleotide or longer loops introduced steric clashes at the strand entry and exit positions (Figure S3) and were not pursued. The resulting alphaKL brings together several stabilizing interactions — A-minor packing, a G-major triplex, and a programmable four-nucleotide kissing loop — into a compact connector that links helices along their edges (Figure 1e–g).

Because RNA origami tiles are built on a precise helical geometry, where each base pair occupies a defined position along the helix, even single base pair variations can distort the overall tile shape.^21^ The alphaKL was therefore engineered to preserve canonical A-form helical alignment without introducing additional bend or twist. A single unpaired uridine at the 3’ loop terminus relieves local strain while maintaining backbone continuity.

Geometric compatibility was validated using fragment-based assembly from high-resolution RNA structures, a technique that was previously applied to engineer artificial RNA junctions with A-minor interactions.^1^ An alphaPK fragment from helix 18 of the *T. thermophilus* 16S rRNA was embedded within an ideal A-form helix, expanded to accommodate the four-nucleotide loop, and fused to a triplex-junction fragment from the phage *Twort* group I ribozyme. After stereochemical refinement using QRNAS, ^22^ the motif was incorporated into the RNA Origami Automated Design (ROAD) platform^3^ and pyFuRNAce,^20^ enabling sequence generalization and integration into RNA origami tiles. We extended ROAD to support 3D modeling of alphaKL motifs and included additional sequence optimization routines that improve the predicted RNA folding energetics and partition function. Additionally, the alphaKL motif has been implemented in the pyFuRNAce webserver,^20^ making the design of edge-to-edge connections in RNA origami accessible with a graphical user interface for labs around the world. The resulting models are geometrically consistent and suitable for molecular dynamics (MD) simulations and experimental implementation.

### AlphaKL Assembly Characterization by AFM

We designed a series of constructs to characterize alphaKL function in RNA origami using our enhanced version of the ROAD design pipeline.^3^ All constructs were transcribed and assembled *in vitro* at 37°C (without annealing step) and analyzed by atomic force microscopy (AFM) to evaluate folding and self-assembly behavior.

To test whether alphaKLs could mediate edge-to-edge association, we designed two versions of a nanoring-forming RNA tile: ^9^ a control lacking alphaKL connectors, and a variant with three alphaKL sites per outer edge – one self-complementary and two complementary flanking sites (sc and b-b’ connectors, Figure 2a,b). Cotranscriptional folding of tiles lacking alphaKL connectors produced monomeric nanorings composed of four to six RNA tiles (Figure 2a and c, Figure S4). Tiles bearing three alphaKLs per edge further assembled into an extended network of connected nanorings (Figure 2d), in which taller regions at the ring interfaces are consistent with the expected locations of individual alphaKL connectors (see inset and Figure S5). AFM height profiles matched the expected ∼ 33 nm outer diameter predicted by assembling ROAD models into hexamers (Figure S6). The measured average spacing between connectors was 5.77 ± 0.30 nm (N=10), in close agreement with the designed separation of two helical turns (6.18 nm) (Figure S7). Occasional double-height rings at some assembly edges are consistent with partial multilayer stacking (Figure 2d, S6).

**Figure 2:**
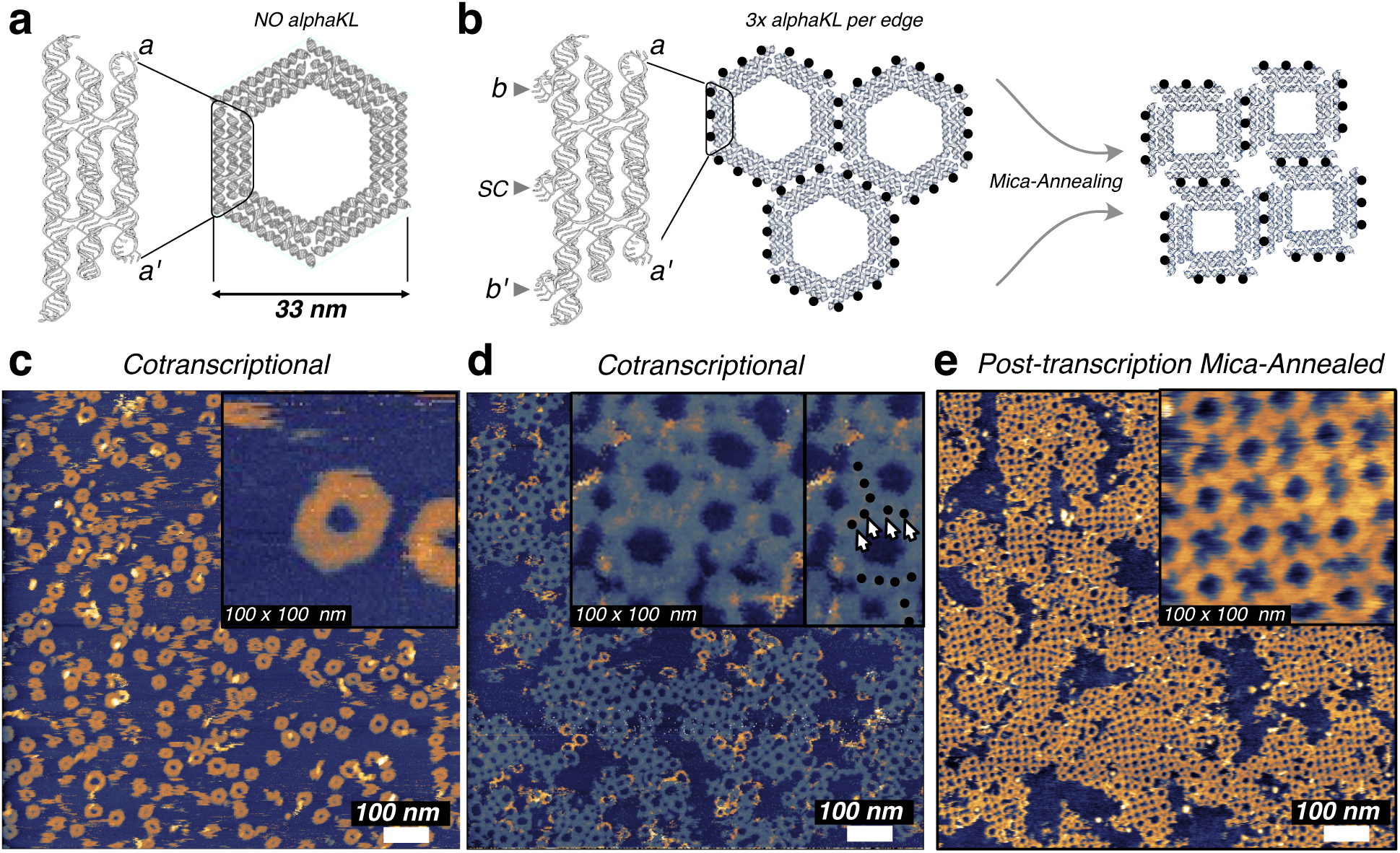
AlphaKL enables side-by-side assembly of RNA origami nanorings into surface tilings. **a)** Design schematic of RNA tiles assembling into multimeric rings via 120° KL a–a’ connectors. Without alphaKLs, these rings are unable to interlink. **b)** Assembly schematic of tiles with three alphaKLs per long edge: two complementary (b-b’) and one self-complementary (sc). **c)** AFM image of control tiles lacking alphaKLs directly after co-transcriptional folding. **d)** AFM image of tiles bearing three alphaKLs each directly after cotranscriptional folding. Inset: enlarged view highlighting individual alphaKL connectors (dots with increased height). **e)** AFM of the same design from (d) annealed to mica post-transcription (55°C – 25°C, 90 min).

In a separate experiment, samples were subjected to post-transcriptional annealing on mica (55°C–25°C). This treatment yielded more regular and densely packed assemblies (Figure 2e) compared to unannealed controls (Figure S10). Whereas cotranscriptional products displayed a mixture of hexagonal and square lattices (Figure S4), annealed samples predominantly formed square lattices (Figure S8–S9). These observations indicate that alphaKL-mediated assemblies form cotranscriptionally and remain capable of limited structural rearrangement under mild annealing conditions.

### Predictive Modeling of Sequence Variants

In the natural alphaPK motif, a dinucleotide platform caps one end of the KL interface and defines a sharp turn on the minor-groove side of the *α*-shaped loop. Dinucleotide platforms are planar stacking motifs that create defined interfaces for tertiary interactions and stabilize long-range RNA packing through *π*-stacking and backbone organization.^23,24^ Phylogenetic analysis shows that this minor-groove region varies across species (Figure S1), with CA being the most frequently observed configuration among alphaPK structures in the PDB, and AA and GG representing natural alternatives documented in the RNA 3D Atlas.^25^ This natural variation motivated us to systematically probe how the platform sequence influences tertiary stabilization (Figure S11). The two A-minor triples that form after the 4-nt KL are named ‘A-minor pos 1’ and ‘A-minor pos 2’, where pos 2 corresponds to the most strictly conserved minor groove contact in the natural motif. We embedded each variant into a minimal hairpin dimer design to isolate local structural effects, naming them according to the nucleotides flanking the 4-nt KL (5’-M—NNNN—mmm-3’): G/CAA (CA platform), G/AAA (AA platform), G/GGA (GG platform), G/UGA (synthetic variant), and U/AUU (negative triplex control) (Figure S13).

Conventional RNA structure prediction tools did not reliably reproduce the engineered dual-triplex topology, including physics-based approaches such as SimRNA^26^ and fragment-assembly methods such as RNAcomposer^27,28^ (Figure S12). To better capture the composite motif’s geometry, we turned to AlphaFold3,^29^ reasoning that its training on native RNA structures might allow partial recognition of this synthetic fold. We modeled minimal alphaKL dimers in the presence of 50 explicit Mg^2+^ ions (Figure 3a) which can help stabilize triplex interactions. All variants formed loop-loop contacts with coaxially aligned helices (Figure 3a); however, confidence scores (pLDDT) were low across all variants, and markedly lower for the unstructured negative control U/AUU (Figure S12). The low pLDDT scores are expected for a synthetic motif absent from the training database, yet AlphaFold3 consistently recovered the characteristic *α*-shaped loop topology more reliably than the coarse-grained tools, which produced this conformation only sporadically. These results suggest that, despite being synthetic, the composite motif shares sufficient geometric features with native tertiary contacts to be partially recognized by AlphaFold3.

**Figure 3:**
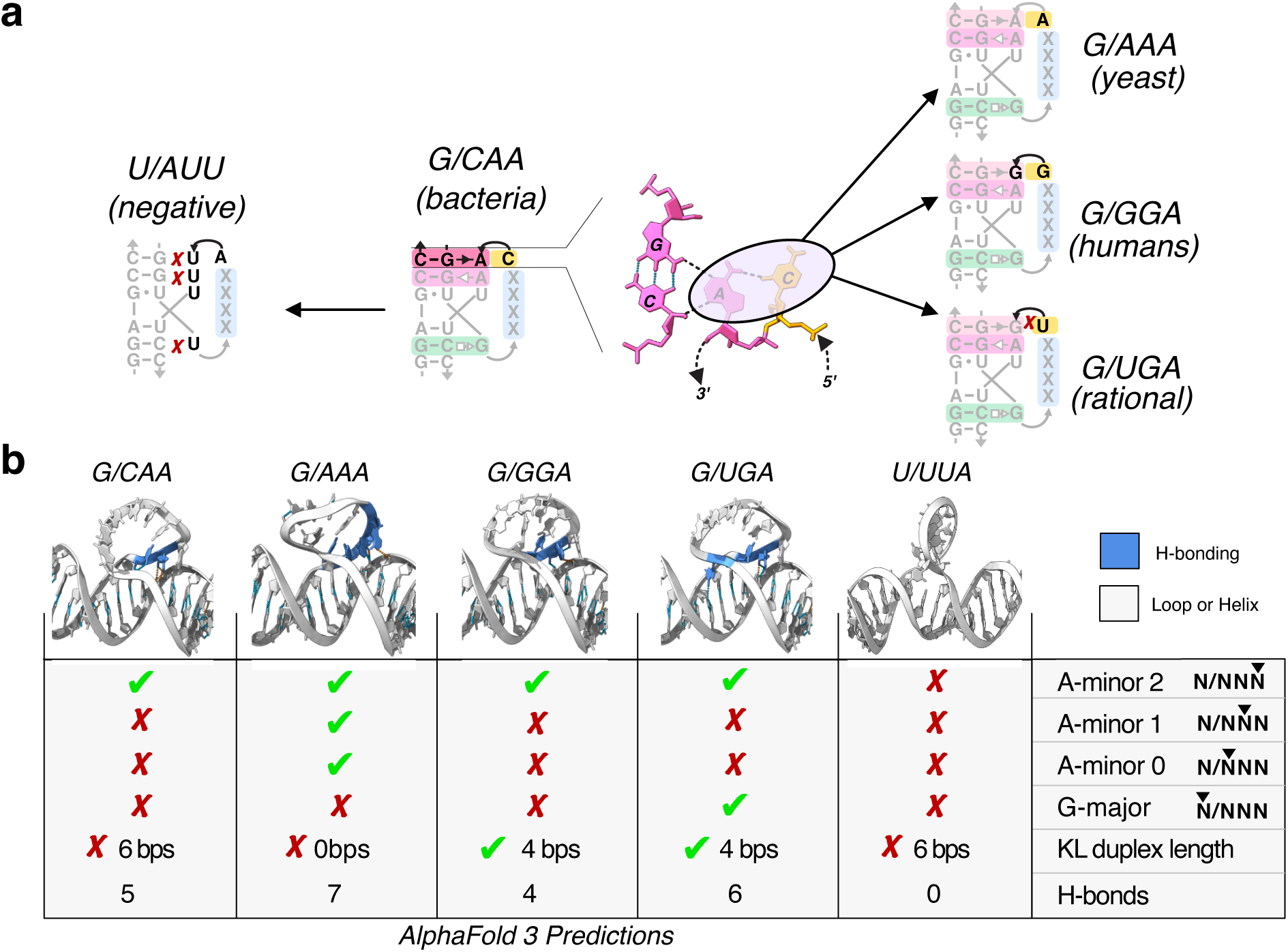
Structural hypotheses for alphaKL sequence variants from AlphaFold3 modeling. Although confidence scores are low for this synthetic motif, AlphaFold3 consistently recovers the *α*-shaped topology and generates differential predictions across platform variants that inform sequence–structure hypotheses tested experimentally. **a)** Diagram of alphaKL sequence variants tested. **b)** AlphaFold3-predicted alphaKL hairpin dimers with 50 explicit Mg^2+^ ions. Zoom-ins of the simulated alphaKL loops in dimeric context are shown here, with blue colored nucleotides indicating formation of H-bonds between the loop and helix. A tabular summary of identified interactions is further provided.

In G/CAA, the flanking C-G pairs adjacent to the KL are mutually complementary, and AlphaFold3 consistently extended the designed 4-bp KL into an unintended 6-bp KL. This alternative structure competes with the intended *α*-topology, and can be suppressed by choosing a platform sequence that lacks this complementarity.

The recurrent extension of the KL in this context points to an alternative structure that can be selectively suppressed by picking a different platform sequence. In contrast, the G/AAA variant stabilized both A-minor positions, and positioned a third adenosine to reinforce minor-groove packing, yielding the highest total number of hydrogen bonds (Figure 3b). The G/UGA variant stabilized one A-minor and the G-major triplex, whereas the G/GGA variant formed only a single A-minor interaction. As expected, the U/AUU control failed to form stable triplex contacts, but also adopted an extended 6-bp kissing geometry (Figure 3b).

These models suggest that single-nucleotide substitutions at the platform are predicted to differentially modulate triplex occupancy, hydrogen-bonding networks, and interface geometry. Replacing C with A in the G/AAA design suppresses unintended C-G extension while preserving minor-groove triplex interactions. Additionally, the G/UGA variant strengthens the major-groove triplex at the expense of minor-groove stabilization (Figure 3c). Together, these predictions suggest a sequence-structure tradeoff in which both the minor- and major-groove triplex contacts engage the same loop segment, and therefore stabilizing one interaction may come at the expense of the other.

### Comparison of alphaKL Sequence Variants by AFM

Each alphaKL variant was incorporated into a rectangular RNA origami tile designed to polymerize into linear fibrils, directly testing whether the predicted structural differences translate into measurable assembly outcomes (Figure 4a). This experimental series directly tests the structural hypotheses generated above: variants predicted to cleanly adopt the designed fold should produce not only longer but more regularly organized fibrils, while those susceptible to competing conformations should show increased heterogeneity and disorder.

**Figure 4:**
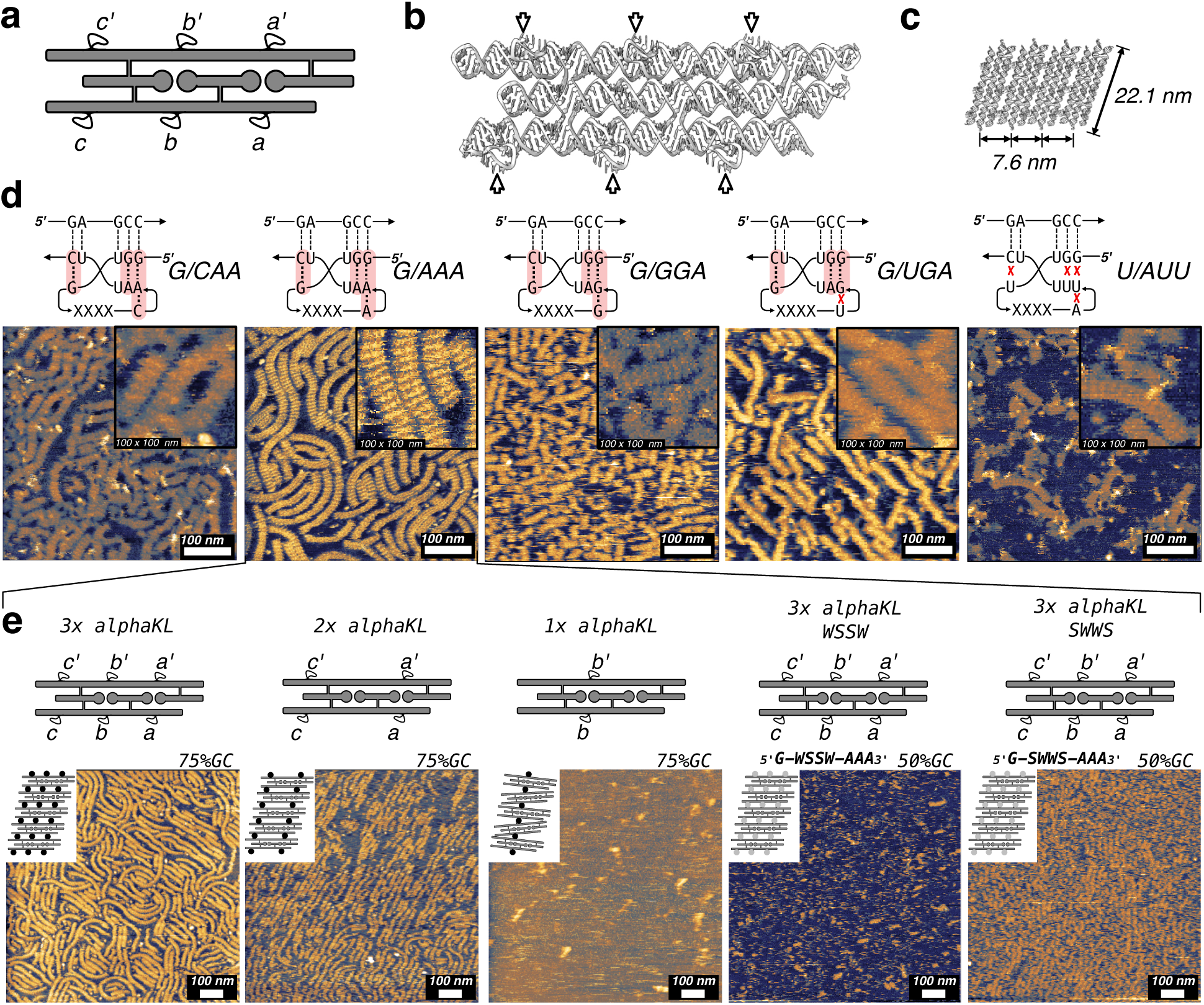
AlphaKL-platform sequence influences regularity of alphaKL-mediated fibril assembly. **a)** Strand diagram of a rectangular RNA tile designed to form linear fibrils via three alphaKL sites per edge (a–a’, b–b’, c–c’). **b)** Theoretical model generated by ROAD software. AlphaKLs are indicated by arrows. **c)** Diagram showing assembly scheme, predicted size, and spacing of units. **d)** AFM images of fibrils cotranscriptionally assembled in solution, for different motif variants. **e)** AFM images of tiles bearing three, two, or one alphaKL per edge, and with lower energy alphaKLs with either WSSW or SWWS constraint.

AFM imaging of cotranscriptionally assembled samples (Figure 4d, S14) revealed pronounced differences in fibril length and organizational regularity across variants. Pairwise comparisons with Bonferroni correction reveal a three-tier hierarchy of assembly performance (Figure S15). G/AAA and G/UGA formed the longest and most continuous fibrils (mean 148 ± 99 nm and 130 ± 97 nm respectively) and were statistically indistinguishable from each other (p_adj_=1), establishing them as the highest-performing variants of the alphaKL. G/CAA and G/GGA formed a second tier of intermediate assembly (mean 111 ± 80 nm and 73 ± 28*nm* nm, respectively), again not significantly different from each other (p_adj_=1), though G/GGA was significantly shorter than G/AAA (p_adj_=0.00337). The U/AUU negative control was significantly shorter than all other variants (p_adj_≤0.0001) producing predominantly short fragments and aggregates (mean 53 ± 34 nm).

ROAD modeling estimates a periodic spacing of 7.6 nm between fibril units (Figure 4b,c) consistent with the measured segmentation lengths of 6.8 ± 0.9 nm (N=17) (Figure S16). Fibril length distributions also correlate with migration patterns observed by native PAGE (Figure S17). AFM likely underestimates true contour length due to surface-induced fragmentation of twisted filaments, as reported for DNA origami ribbons.^30^ Thus, shorter fibril lengths observed for some alphaKL variants may partially reflect surface-induced breakage of strained assemblies rather than intrinsic limits on polymerization in solution.

We next examined the dependence of assembly stability on the strength and number of alphaKL connectors using the G/AAA platform. Tiles bearing three, two, or one G/AAA alphaKL per interface were compared by AFM (Figure 4e, left). Reducing the number of connectors progressively shortened fibril length and increased heterogeneity. We further modulated connector strength by reducing GC content of the KL interactions (75%-GC versus 50%-GC variants; Figure 4e, right). Assemblies with lower GC content were observed to have reduced fibril length and stability. Extended, well-defined fibrils required three alphaKL connections per interface with 75% GC content, whereas fewer or weaker connections produced shorter and more dynamic assemblies.

### All-Atom Molecular Dynamics of AlphaKL Variants

A central question for alphaKL design is whether local triplex contacts translate into measurable rigidity at the helix-helix interface. Five independent all-atom MD simulations of minimal monomer and dimer systems for each variant quantified triplex persistence and inter-helical orientation for G/CAA, G/AAA, G/GGA, G/UGA, and U/AUU (Figures S18–S23). To examine whether this behavior scales, we also simulated a dimer of two full three-helix tiles connected by three G/AAA alphaKLs (Figure 5c, S24) in a system comprising nearly 30,000 RNA atoms and 1.9 million solvent atoms. Together, these two datasets reveal a two-level stabilization hierarchy: triplex contacts determine the preferred interhelical angle at the motif level, while accumulated backbone-backbone hydrogen bonds progressively rigidify the interface at the tile scale.

**Figure 5:**
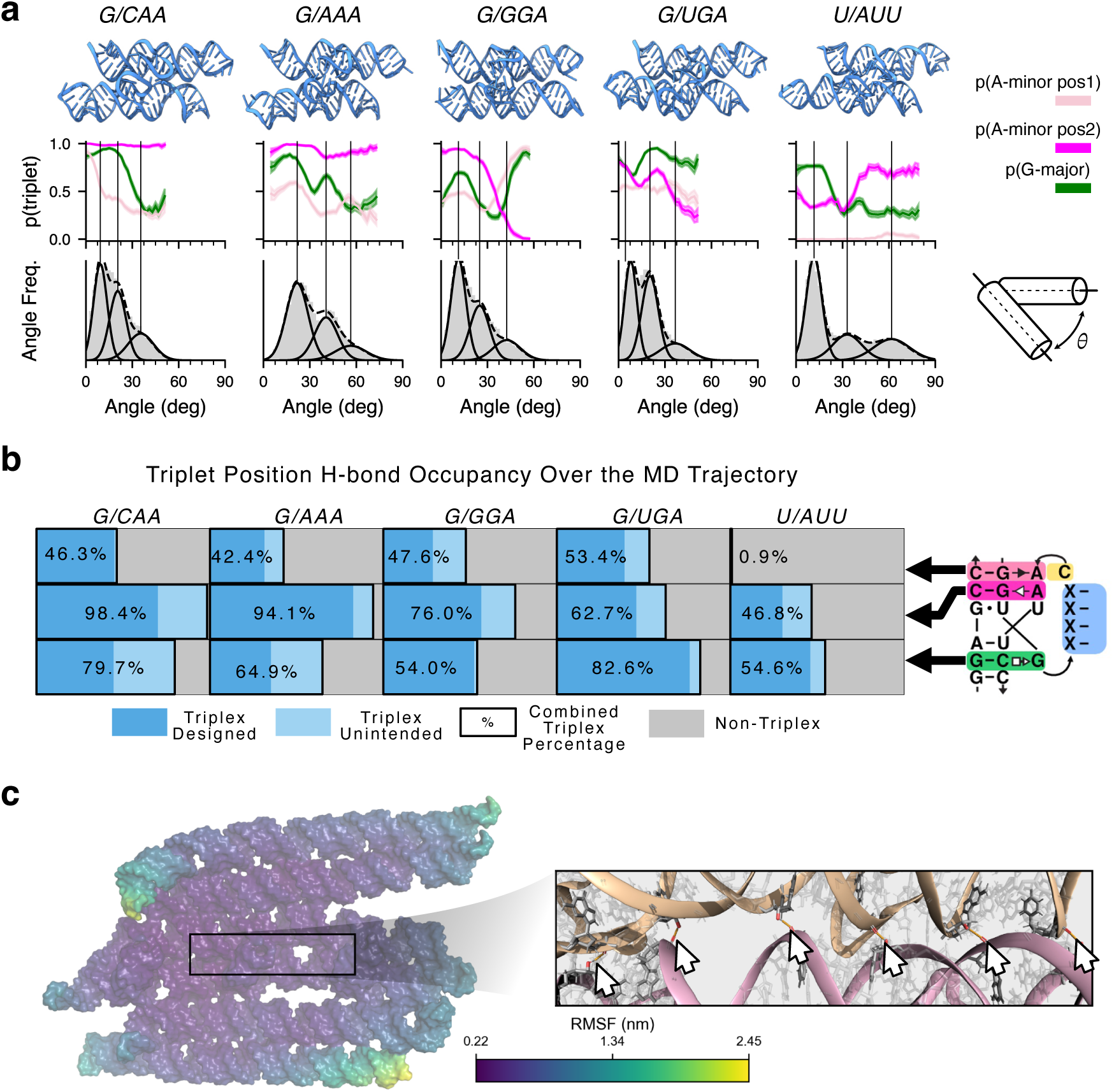
Atomistic modeling and molecular dynamics simulations. **a)** (top) Final configuration from one of the five 2 µs MD simulations of alphaPK motif variants dimerizing two short hairpins. (bottom) Angle distribution of the duplex (gray) plotted against the probability that each nucleotide is involved in a triplex in each angle bin (colors). The angle *θ* is measured from the helical axis of the stems where the alphaKLs are embedded. The angle distributions were fit with a three-term Gaussian mixture model (black lines). **b)** Hydrogen bond occupancy of the three positions that form triplex interactions. **c)** (left) Atomistic MD simulation of a full-sized RNA origami tile-tile interaction containing three aligned alphaKLs. The average configuration from a 500 ns simulation of the the dimer is shown with a surface rendering corresponding to the RMSF of each atom. (right) A close-up view of the tile-tile interface in a final simulated configuration with extensive backbone-backbone hydrogen bonds, indicated by white arrows.

At the motif level, the interhelical angles (*θ* angles) depend on which triplex contacts are active. Variants with persistent contacts, such as G/AAA and G/UGA, stay near a preferred *θ* angle, while G/GGA and U/AUU sample a much broader range (Figure 5a, S36–S37). The hydrogen-bond occupancy analysis clarifies why: peaks in the angle distribution correspond to peaks in the hydrogen-bond occupancy, demonstrating the importance of tertiary contact formation for the overall geometry (Figure 5a, b). A-minor pos2 is the dominant contact, with occupancy above 94% in both G/CAA and G/AAA, while A-minor pos1 contributes more modestly across all variants. G-major occupancy is where the variants diverge most: G/UGA leads at 82.6%, while G/GGA and G/AAA show intermediate persistence. Notably, G/AAA matches G/CAA on A-minor stabilization despite producing more uniform assemblies, supporting the conclusion that the performance difference comes from suppressing the competing six-base-pair duplex rather than from stronger triplex contacts. The U/AUU negative control lacks sustained occupancy at all positions, confirming that the triplex sequences are essential for interface stability. These motif-level results establish what each variant contributes at a single junction, but leave open whether the behavior scales when multiple alphaKLs act in concert.

Our simulation of tile-tile interactions addresses the scaling question directly (Figure 5c, S24). Over 500 ns, the alphaKLs remained intact and showed the lowest RMSF values in the structure, highlighting the interface as its most rigid region. Beyond triplex stabilization, lateral helix packing gave rise to collective backbone-backbone hydrogen bonds in the form of ribose zippers — ribose-ribose (2’OH-2’OH) and ribose-phosphate (2’OH-O_P_). These types of contacts that were rare in the minimal dimer simulations but accumulated readily when three alphaKLs aligned (Figure 5c (right), S35). Each additional backbone hydrogen bond was associated with a mean interhelical angle decrease of 2.49*^◦^* (Figure S32), progressively narrowing the interface as contacts built up. Multiple aligned alphaKLs therefore rigidify the interface through a mechanism that no single motif can produce alone, motivating the tile-level design optimizations in the following section.

### Optimization of AlphaKL Tile Geometry

With the input from the MD simulations, we returned to design optimization. Fibril assembly quality scales with the number and strength of alphaKL connections per interface. Attempts to extend this into 2D by adding 180° KL contacts along fibril edges produced entangled aggregates rather than ordered lattices (Figure S42), indicating that strong irreversible contacts prevent the error correction needed for cooperative nucleation. Two new constructs were therefore designed using alphaKL contacts exclusively, targeting either 1D fibril elongation or 2D lattice growth.

For two-dimensional assembly, we designed a minimalist wireframe square with four 90°-bend motifs,^31^ two alphaKLs per edge, and a single 180° KL closing each corner (Figure 6a,b). After cotranscriptional folding these constructs initially appeared disordered under AFM, consistent with weakly associating tiles below the nucleation threshold. Time-resolved high-speed AFM imaging revealed gradual emergence of ordered domains, expanding over successive scans into essentially defect-free square lattice domains (Figure 6c, S43), with individual tile cavities clearly resolved in the inset. This nucleation-and-growth behavior suggests that two alphaKLs per edge places tile-tile contacts in a sufficiently reversible range for cooperative lattice formation.

**Figure 6:**
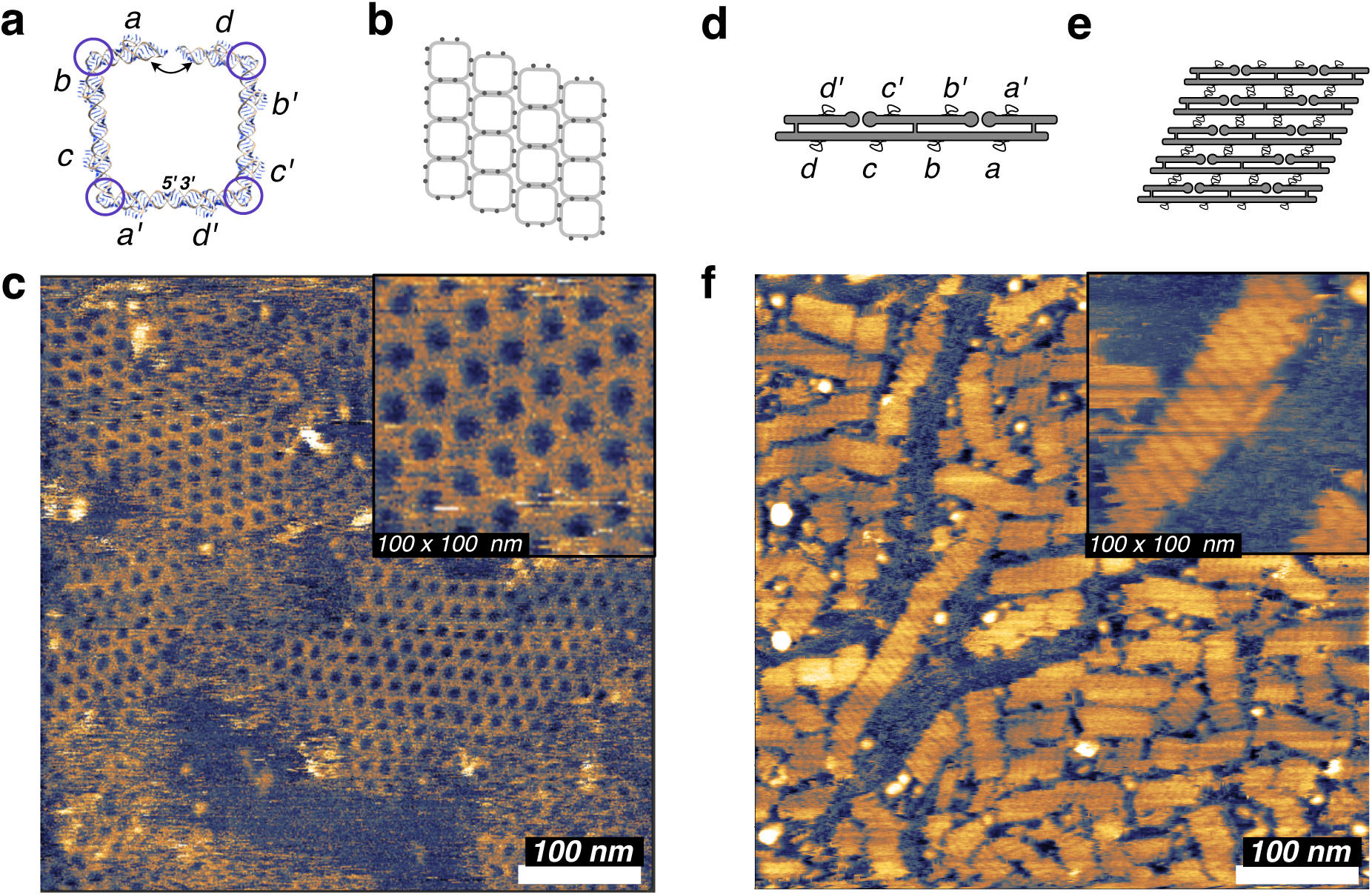
Optimization of alphaKL tile geometry for 1D and 2D assembly. **a)** Design schematic of a wireframe square tile assembled from four 90°-bend motifs, with two alphaKLs per edge (a–d, a’–d’) and 180° KL corner closures. **b)** Expected square tiling pattern upon 2D assembly. **c)** AFM image of cotranscriptionally folded square tiles after 1 h on mica at room temperature, showing nucleated square lattice domains with low defect density. Inset: high-magnification view with individual tile cavities resolved. **d)** Schematic of two-helix ribbon tile designs with four evenly spaced alphaKLs per edge. **e)** Expected ribbon assembly geometry for the evenly spaced design. **f)** AFM image of cotranscriptionally assembled ribbon fibrils, showing straighter and wider assemblies than the three-helix design. Inset: high-magnification view showing internal ribbon periodicity. Scale bars, 100 nm.

For one-dimensional assembly, we redesigned the three-helix fibril tile by removing the central helix and introducing crossovers, generating a symmetric two-helix ribbon architecture with four evenly spaced alphaKLs per edge (Figure 6d,e,f). A closely spaced control design produced no detectable assembly (Figure 6d, top), establishing a minimum geometric separation for independent motif folding. With correct spacing, the four-connector interface produced straighter fibrils, with curvature reduced approximately twofold relative to the three-helix G/AAA design (∼ 0.005nm*^−^*^1^ versus ∼ 0.010nm*^−^*^1^, Figure S44). Together, these results indicate that alphaKL interaction strength governs assembly dimensionality: weaker contacts enable cooperative 2D lattice formation, while stronger contacts promote 1D fibril elongation.

## Discussion

The alphaKL defines a previously inaccessible class of RNA connector. Existing programmable RNA connectors — including 180° KLs,^5^ 120° KLs,^9^ branched KLs,^10^ and paranemic crossover interactions^6–8^ — link helices end-to-end through coaxial stacking (Figure 1a), coupling both recognition and stability to helix termini. While lateral contacts occur in natural RNA, such as loop–receptor interactions in TectoRNA systems,^32,33^ these rely on structurally specific motifs drawn from a limited family of sequences^23^ and are not readily interchangeable within modular design frameworks. DNA nanotechnology has shown that lateral connectivity is geometrically feasible through shape complementarity^34^ or triplex interactions,^35–37^ but an RNA equivalent compatible with A-form geometry and cotranscriptional folding has been lacking. The alphaKL fills this role for RNA, providing a sequence-programmable connector that links helices along their edges rather than at termini (Figure 1b). Among all 256 possible 4-nt sequences, 32 variants with three G–C pairs are predicted to form stable interfaces (Figures S45–S46). This subset, when combined with the requirement for multiple matched KLs per interface, suggests that even a modest sequence space is sufficient to support complex, addressable RNA assemblies.

AlphaKL behavior is governed by a sequence–structure hierarchy across two scales. At the motif level, single-nucleotide substitutions at the dinucleotide platform redistribute stabilization between minor- and major-groove triplex contacts. G/CAA and G/AAA exhibit comparable triplex stability when initialized in the designed conformation; the observed performance gap reflects suppression of a competing six-base-pair duplex rather than differences in intrinsic triplex strength. G/UGA achieves comparable assembly through a complementary mechanism, trading minor-groove stabilization for increased G-major triplex occupancy, consistent across both AlphaFold3 predictions and molecular dynamics simulations. Despite reduced confidence scores associated with the novelty of the motif, AlphaFold3 reliably distinguishes variants that adopt the designed fold, while molecular dynamics quantifies their stability once formed. Together, these approaches provide a coherent mechanistic framework that explains experimental outcomes in the absence of atomic-resolution structures. The differences in assembly observed by AFM emerge directly from the balance between competing tertiary interactions rather than the strength of any single contact.

At the tile scale, simulations reveal an emergent stabilization mechanism absent from the isolated motif. Arrays of aligned alphaKLs accumulate backbone–backbone hydrogen bonds (ribose zippers) across the helix–helix interface, with each additional contact associated with a mean interhelical angle decrease of 2.49*^◦^*. Individual motifs define local geometry, but collective accumulation drives global straightening and rigidification beyond what any single junction can achieve. This scaling behavior makes copy number a direct design variable: two alphaKLs per edge maintain reversible interactions compatible with cooperative two-dimensional lattice nucleation, whereas three or four promote robust one-dimensional fibril elongation. The resulting thresholded avidity parallels multivalent DNA origami assembly^38,39^ and introduces a geometry-based assembly gate to the RNA origami toolkit alongside existing regulatory strategies. ^40^

The underlying design principle extends beyond the helix–helix interface demonstrated here. Expanding the loop to accommodate longer sequences allows the same triplex-stabilized architecture to present programmable single-stranded elements, enabling nucleic acid capture (Figure S47, top). Structural models further suggest that alphaKL-like motifs can recognize terminal RNA loops, forming branched loop-to-helix complexes with defined geometry (Figure S47, bottom). Although these configurations remain to be experimentally validated, they point to a general strategy in which A-minor-mediated pre-organization enables programmable tertiary recognition across diverse RNA contexts.

The alphaKL is fully compatible with cotranscriptional folding and integrates directly into one-pot transcription–assembly workflows, including the ROAD^3^ and pyFuRNAce^20^ design platforms. Programmable RNA connectors have already enabled translational applications such as multivalent therapeutic nanoparticles,^41^ and fibers.^42^ By introducing lateral interface geometry, the alphaKL expands the range of achievable architectures and may enable new modes of multivalency and targeting inaccessible to end-to-end assembly. Recent demonstrations of RNA origami assembly in synthetic cells^43^ and in human cells^44^ suggest that access to lateral connectivity will be essential for constructing increasingly complex RNA-based molecular machines and cellular scaffolds.

## Methods

### Alpha Kissing Loop Motif and Variant Design

The alphaKL motif was designed by combining two structural fragments from high-resolution RNA structures. An alphaPK fragment from helix 18 of *T. thermophilus* 16S rRNA (PDB ID: 4V51) was selected from 15 recurrent occurrences catalogued in the RNA 3D Motif Atlas,^25^ providing the minor-groove packing geometry and backbone topology of the *α*-shaped loop. A triplex-forming element from the P4-P6 domain of the *Twort*group I ribozyme (PDB ID: 1Y0Q) contributed the G-major groove contact.^19^ Sequence variants were designed by substituting the dinucleotide platform and flanking loop positions to modulate triplex occupancy, yielding G/AAA (AA platform), G/CAA (CA platform), G/GGA (GG platform), G/UGA (synthetic, enhanced G-major), and U/AUU (negative control)

### 3D Modeling and Structural Refinement

Fragment alignment and backbone grafting were performed in UCSF ChimeraX,^45^ with breakpoints chosen to preserve base stacking and avoid torsional discontinuities. The merged model was stereochemically refined using QRNAS^22^ to optimize bond lengths, torsion angles, and base stacking under secondary-structure restraints. Sequence variants were generated from the refined coordinates using Rosetta *rna-thread*.^46,47^ Complementary alphaKL dimers were modeled by aligning recognition loops along an idealized A-form helix and refining the interface in QRNAS to ensure correct base pairing and backbone register.

### Design of RNA Constructs and Software Development

RNA origami constructs were designed using an extended version of the ROAD platform,^3^ modified to support alphaKL placement along user-specified helical edges while maintaining backbone continuity and A-form registry. Updated motif libraries were incorporated, including a twist-corrected^21^ 180° KL (PDB ID: 7PTQ) and the 90°-bend motif^31^ (PDB ID: 3P59). Sequences were optimized by iteratively minimizing ensemble defect using a gradient-descent script (dragon.pl) while constraining conserved triplex and platform residues, then verified using ViennaRNA 2.7^48^ and NUPACK.^49,50^ All sequences are available in the Supplementary Information (SI Figures S48-S62). The alphaKL module is also available through the pyFuRNAce web interface.^20^

### *In Vitro* Transcription and Cotranscriptional Folding

RNA constructs were synthesized by run-off transcription using T7 RNA polymerase (Thermo Scientific). DNA templates (purchased from IDT) were amplified by PCR (Phusion, NEB) from gBlocks carrying a T7 promoter: 98°C 30s, then 35 cycles of (98°C 10s, 62°C 15s, 72°C 15s), 72°C 10min. Transcription reactions (50 µL, 37 *^◦^*C, 45 min) contained 50 mM Tris-HCl (pH 7.6–7.9), 6 mM Mg(OAc)_2_, 40 mM NaOAc, 40 mM KCl, 1 mM DTT, 1 mM each NTP, 1 U µL*^−^*^1^ T7 RNA polymerase, and 200 ng DNA template. RNA folded cotranscriptionally and was used directly without further annealing or purification.

### Molecular Dynamics Simulations

MD simulations were performed with GROMACS 2025.1 (minimal dimers) or 2023.3 (full tiles)^51^ using the AMBERff99bsc0*χ*_OL3_ force field.^52^ Starting models were built using ROAD and then refined with QRNAS.^22^ The RNA structures were converted to GROMACS format and solvated in TIP3P water^53^ using pdb2gmx. Models were ionized to match experimental conditions: neutralizing Na^+^ plus 40 mM NaCl, 40 mM KCl, and 6 mM MgCl_2_. Five independent replicas were prepared for each dimer system and three for each monomer and tile system. After energy minimization (steepest descent), systems were equilibrated in NVT (100 ps, 310 K, stochastic velocity rescaling^54^) then NPT (100 ps, 1 bar, Parrinello-Rahman barostat^55^), with heavy-atom position restraints throughout (force constant 1000 kJ mol*^−^*^1^ nm*^−^*^1^). Production runs used a stochastic cell-rescaling barostat^56^ for 2 µs (minimal dimers) or 500 ns (full tiles), with GPU-accelerated non-bonded and PME calculations. Simulation frames were saved every 200 ps (minimal dimers) or 500 ps (full tiles) for analysis. The first 100 ns of each trajectory was excluded from analysis. RMSD/RMSF were calculated with GROMACS tools; hydrogen bonds and helix axes were assigned using DSSR^57^ (v2.4.6-2024nov15). Data analysis used custom Python scripts^58–60^ (see code availability) and structures were rendered in PyMOL v3.2.0a^61^ and UCSF ChimeraX.^45^

### Atomic Force Microscopy and Assembly Characterization

Unless otherwise noted, all AFM images were acquired directly from unpurified transcription reactions without post-transcriptional annealing. Immediately after transcription, 3 µL of the reaction mix was deposited onto freshly cleaved mica pre-treated with 50 µL AFM imaging buffer (10 mM Tris-acetate, 1 mM EDTA (TAE), 20 mM Mg^2+^). AFM was performed in fluid on a Bruker JPK NanoWizard 2 in tapping mode (QI or FastScan) with SNL10 and FastScanD cantilevers. Where indicated, samples were annealed on mica by temperature ramping from 55 *^◦^*C to 25 *^◦^*C over 90 min on a hot plate. Images were processed and analyzed in Gwyddion.^62^

## Data Visualization and Analysis

Plot graphing and statistical tests were performed using Matplotlib 3.10.7 and Numpy 2.3.4 for MD simulation data.

## Code Availability

All custom written code was deposited on GitHub at []. https://github.com

## Data Availability

Supporting data is available in the Supplementary Information. All raw data and analysis files used in the study are deposited on HeiData (doi: available upon acceptance).

## Acknowledgements

This work was funded by the European Union (ERC, ENSYNC, 101076997); the Deutsche Forschungsgemeinschaft (DFG, German Research Foundation) under Germany’s Excellence Strategy – EXC-3018/1 – 533587280 (Excellence Cluster SynthImmune) and under TRR 392: Molecular evolution in prebiotic environments (Project number 521256690), as well as the Federal Ministry of Education and Research (BMBF) and the Ministry of Science Baden-Württemberg within the framework of the Excellence Strategy of the Federal and State Governments of Germany. MD simulations were performed on the HPC system Raven at the Max Planck Computing and Data Facility.

