## Supplementary Information for "Programmable Edge-to-Edge Assembly of RNA Nanostructures"

### Figure S1. Phylogenetic variation of alphaPK motif

| Sequence | Counts |
|---|---|
| AGGGCAAG*CU | 2 |
| UGCCGGAA*UA | 2 |
| CGGCCAAC*GG | 1 |
| CGGCUAAC*GG | 1 |
| UCCCCAAG*CA | 1 |
| CCCCCAAG*CG | 1 |
| UCCCAAAA*UA | 1 |
| UCCCAAAG*CA | 1 |
| UCCCUAAA*UA | 1 |

| Sequence | Counts |
|---|---|
| GGGCAA* | 2 |
| GCCGGA* | 2 |
| CCCCAA* | 2 |
| CCCAAA* | 2 |
| GGCCAA* | 1 |
| GGCUAA* | 1 |
| CCCUAA* | 1 |

| Sequence | Counts |
|---|---|
| UGCCGGAA*UA | 3 |

| Sequence | Counts |
|---|---|
| GCCGGA* | 3 |

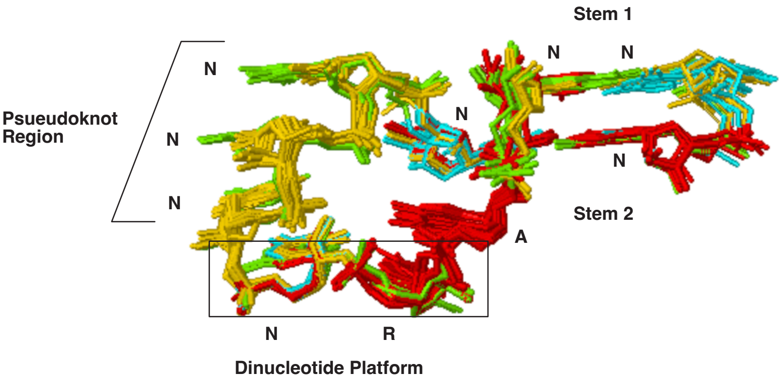


Figure S1: Variants of the alphaPK motif from different ribosomal structures provide us with different options for structural design. In the RNA 3D Atlas this motif is catagorized as ”Motif IL_11411.1” or SSU/LSU pseudoknot, and ”Motif IL_07396.1” for the eukaryotic ribosomal variant with G-G cSH.

**Figure S2. Structural detail of minor and major groove triplexes**

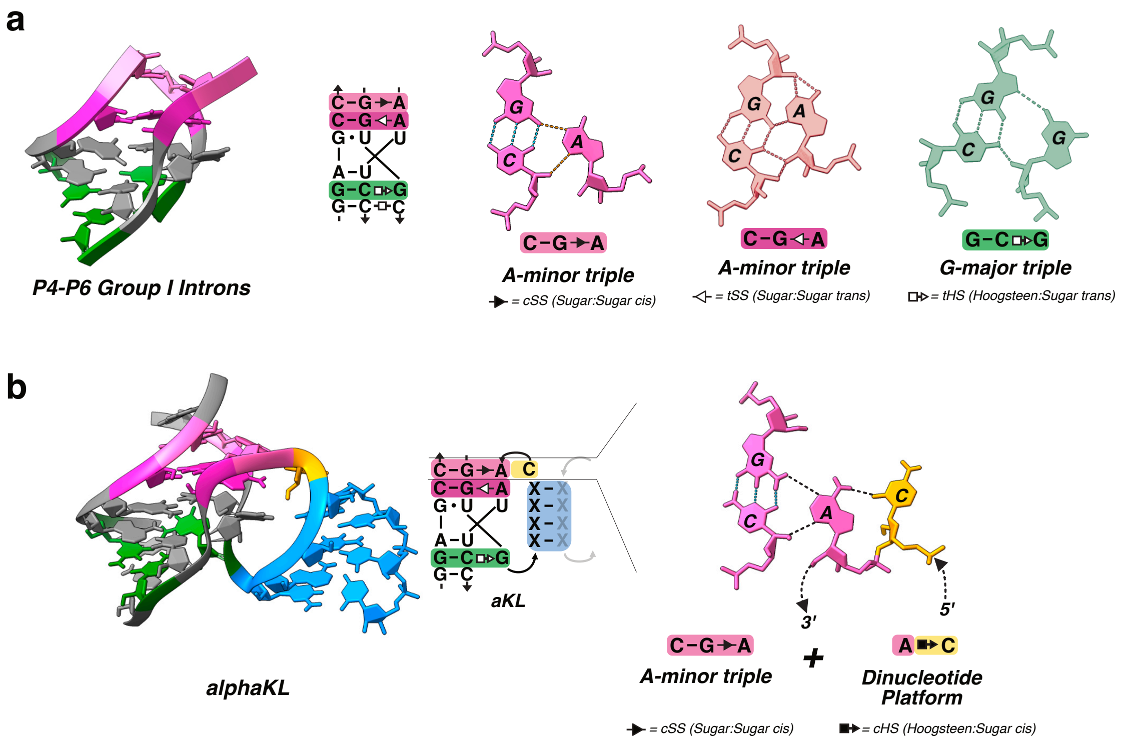


Figure S2: Detailed view of A-minor and G-major triplexes. The dinucleotide platform serves as a means to stabilize a tight turn in the backbone, stacking on the 4 nt KL.

**Figure S3. Clashing arrangement of 5nt alphaKL**

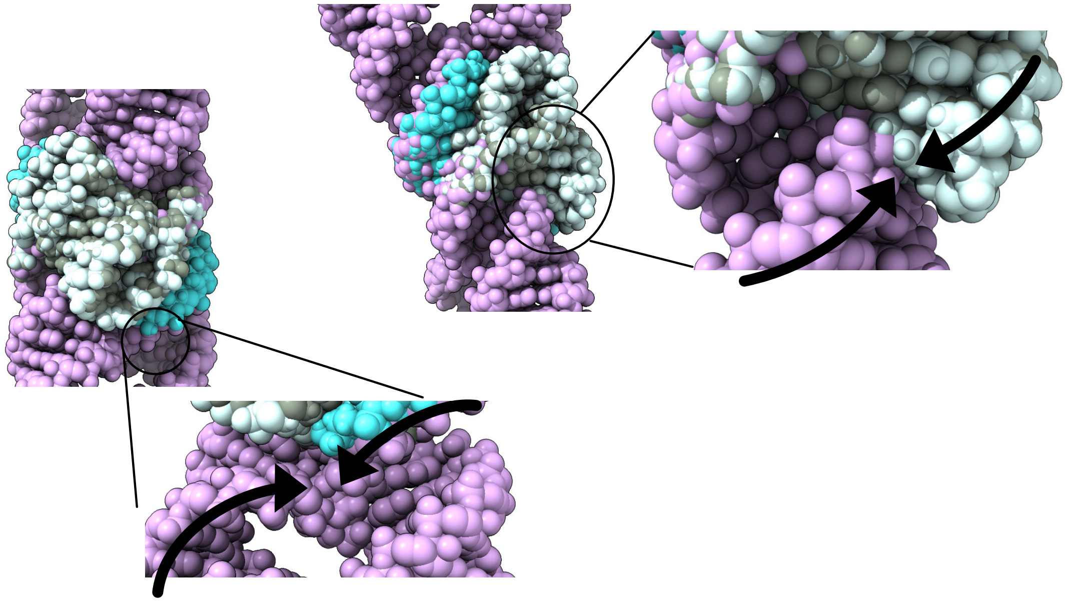

Figure S3: Molecular model of 5 nt alphaKL variant, showing clashing between the motif and partner and a non-parallel angle of interface.

#### Figure S4. Additional AFM images

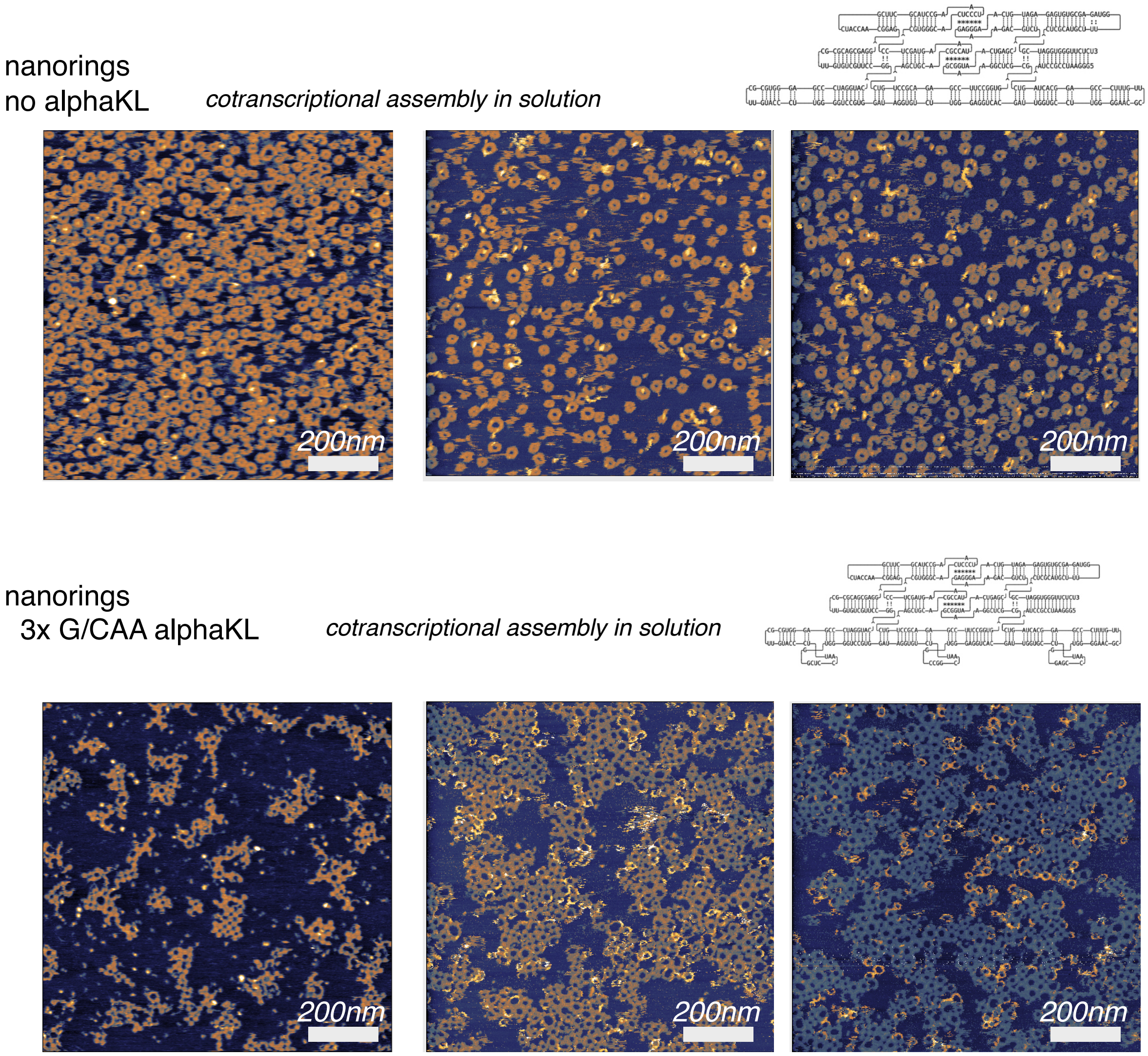


Figure S4: Additional AFM images of nanoring assemblies after cotranscriptional assembly in solution.

#### Figure S5. Additional AFM images

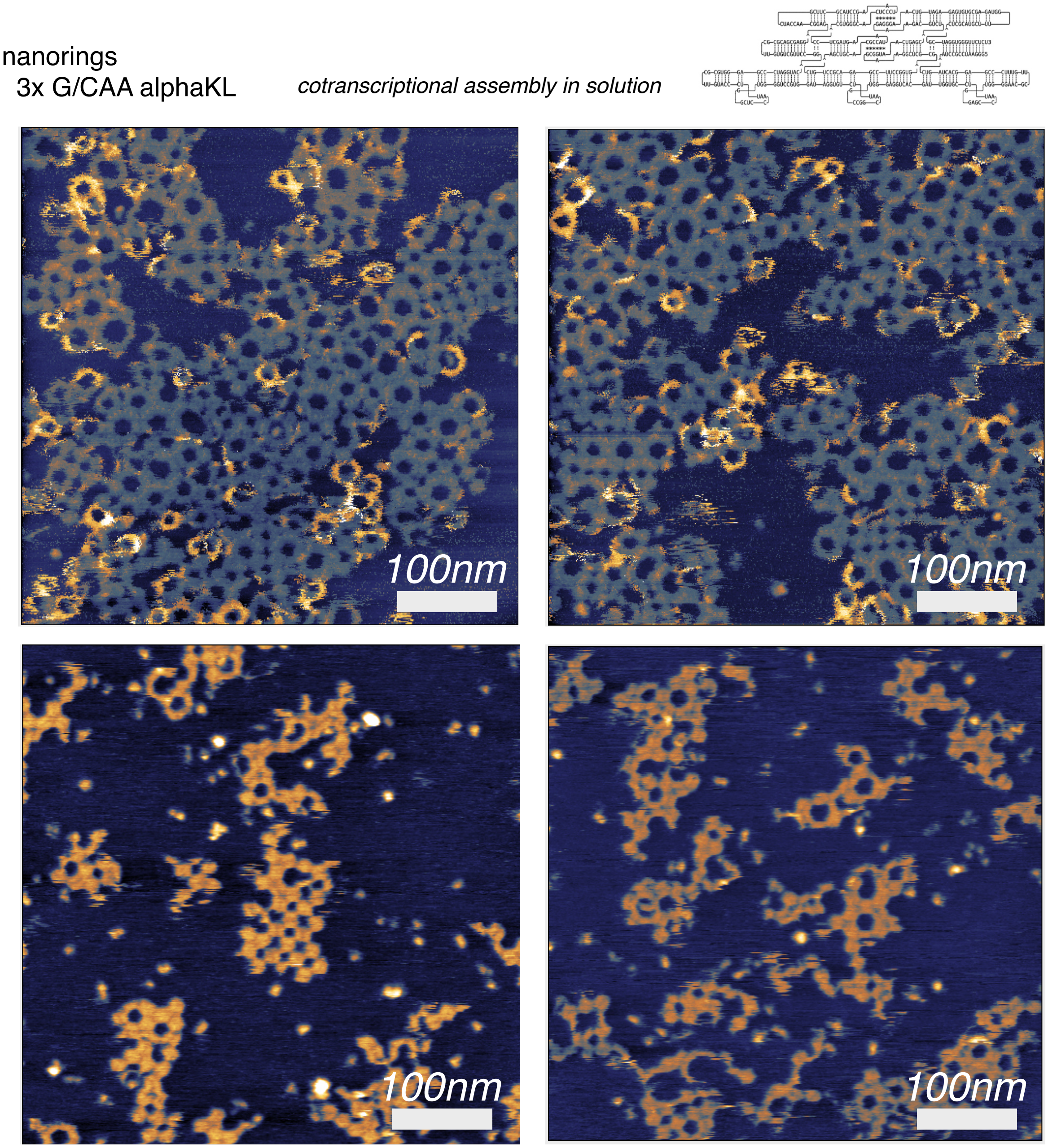


Figure S5: High resolution AFM images of nanoring assemblies after cotranscriptional assembly in solution, showing connections between alphaKL units.

#### Figure S6. AFM height profiles

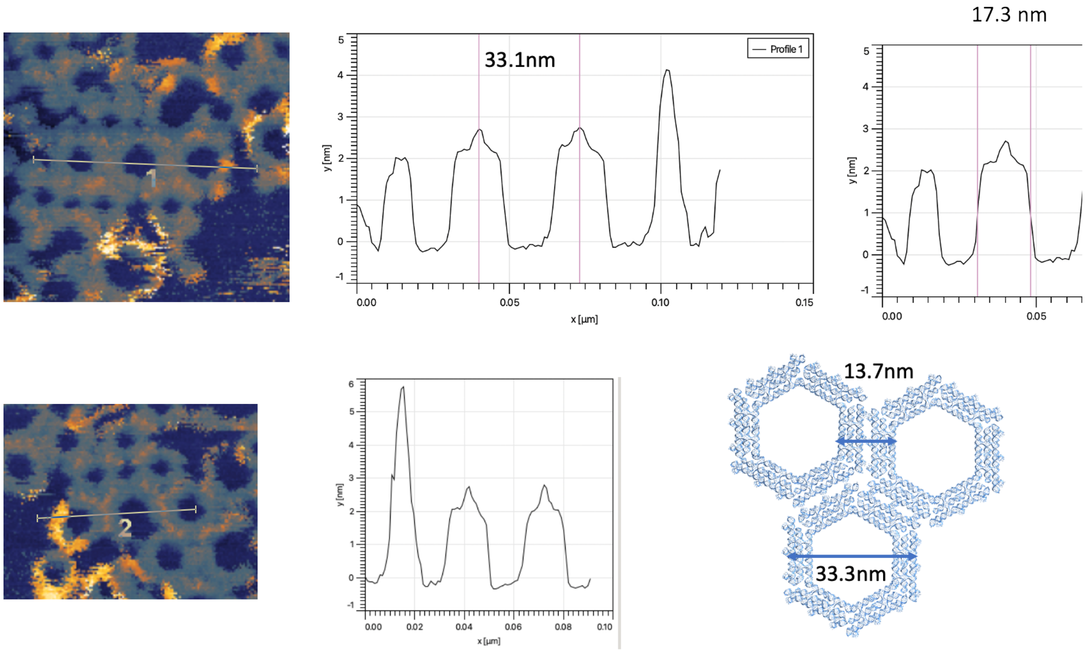


Figure S6: AFM height profile of nanoring assemblies.

**Figure S7. AFM height profile of individual alphaKLs.**

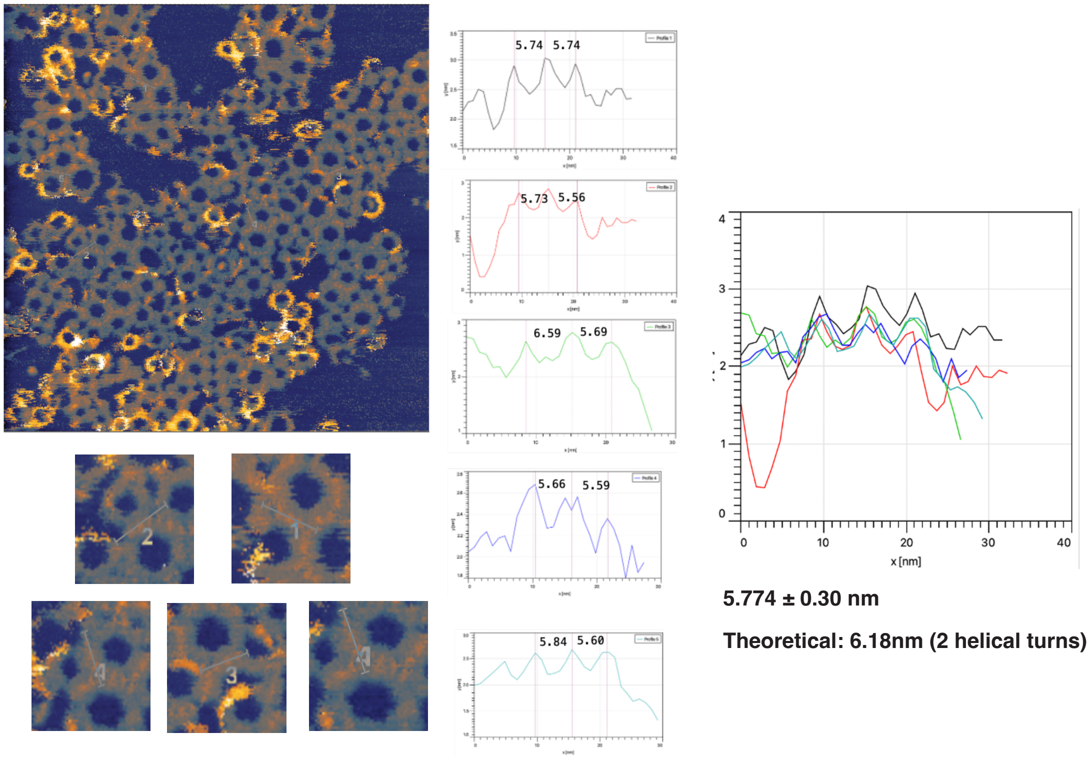


Figure S7: AFM height profile of individual alphaKLs at the connection interface.

**Figure S8. Mica-annealed nanoring lattice with G/CAA alphaKL**

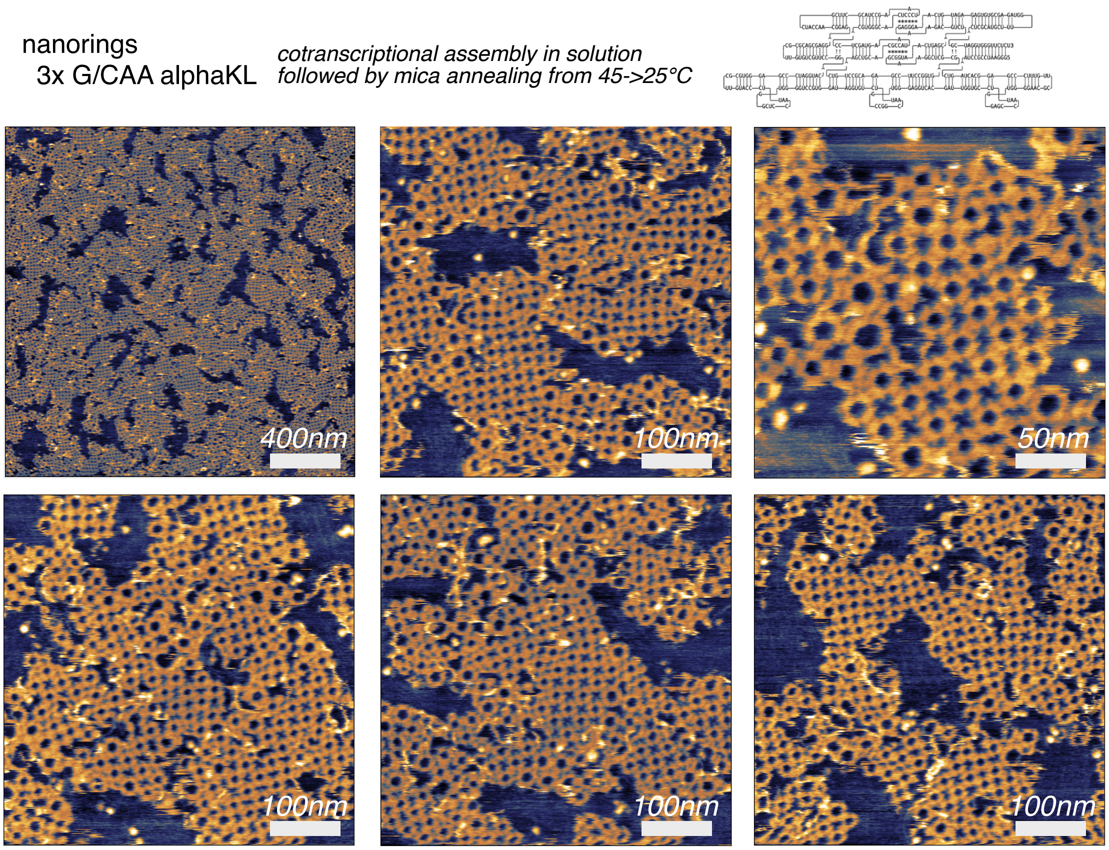


Figure S8: Additional AFM images of nanoring assemblies. After cotranscriptional assembly in solution RNA products were deposited on 45°C mica, and slowly cooled to 25°C over 90 minutes in an insulated and sealed chamber.

#### Figure S9. Mica-annealed nanoring lattice with G/CAA alphaKL

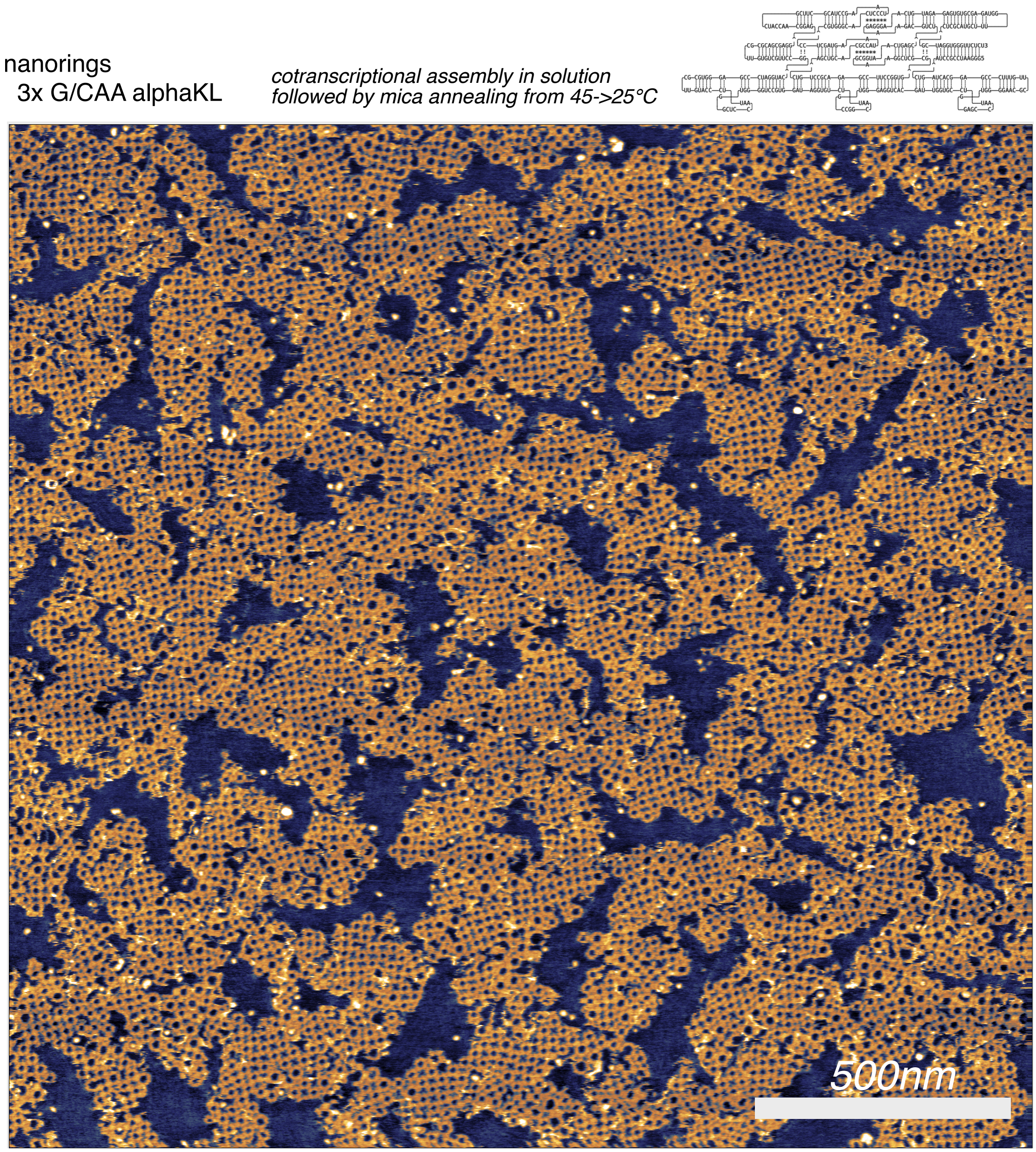


Figure S9: Additional AFM images of nanoring assemblies. After cotranscriptional assembly in solution RNA products were deposited on 45°C mica, and slowly cooled to 25°C over 90 minutes in an insulated and sealed chamber.

**Figure S10. Nanoring-3x-alphaKL domain size measurement**

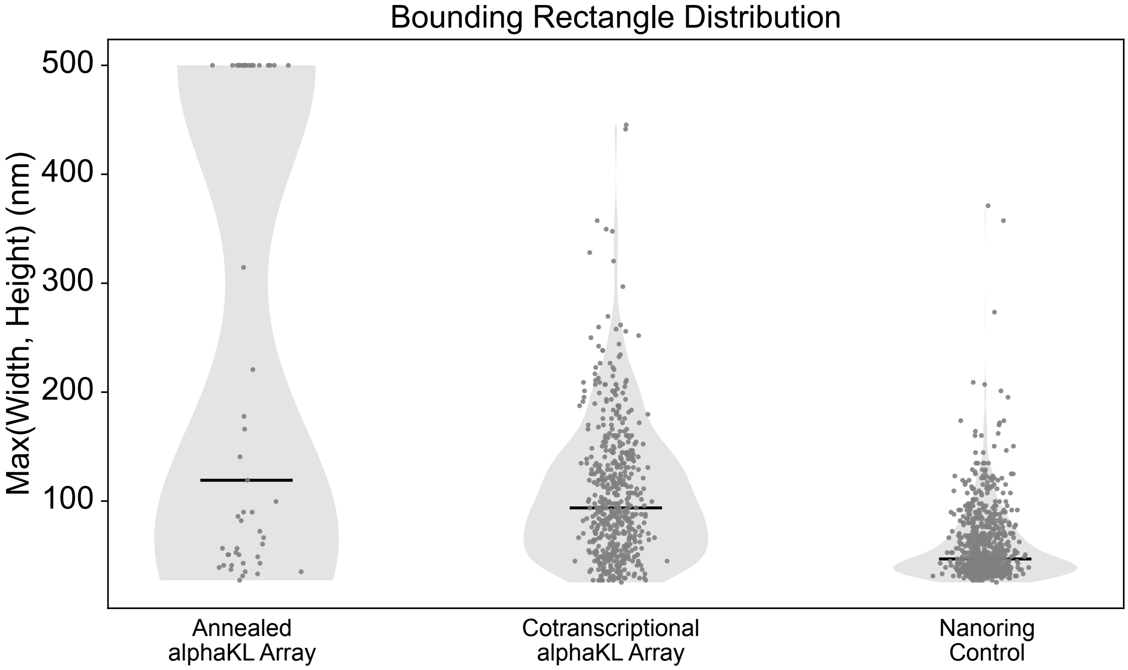


Figure S10: Violin plots showing the maximum dimension of bounding rectangle of single domain assemblies of Nanoring-3x-alphaKL nanorings joined edge-to-edge by alphaKL linkers. RNA samples were prepared by cotranscriptional folding at 37°C for 40 minutes. The annealed sample was deposited on mica and annealed in imaging buffer from 45°C to 25°C over 90 minutes before imaging. The cotranscriptional sample was deposited on mica in imaging buffer and imaged immediately after deposition. The negative control is a nanoring-forming sample that lacks alphaKL connectors.

#### Figure S11. Dinucleotide platforms

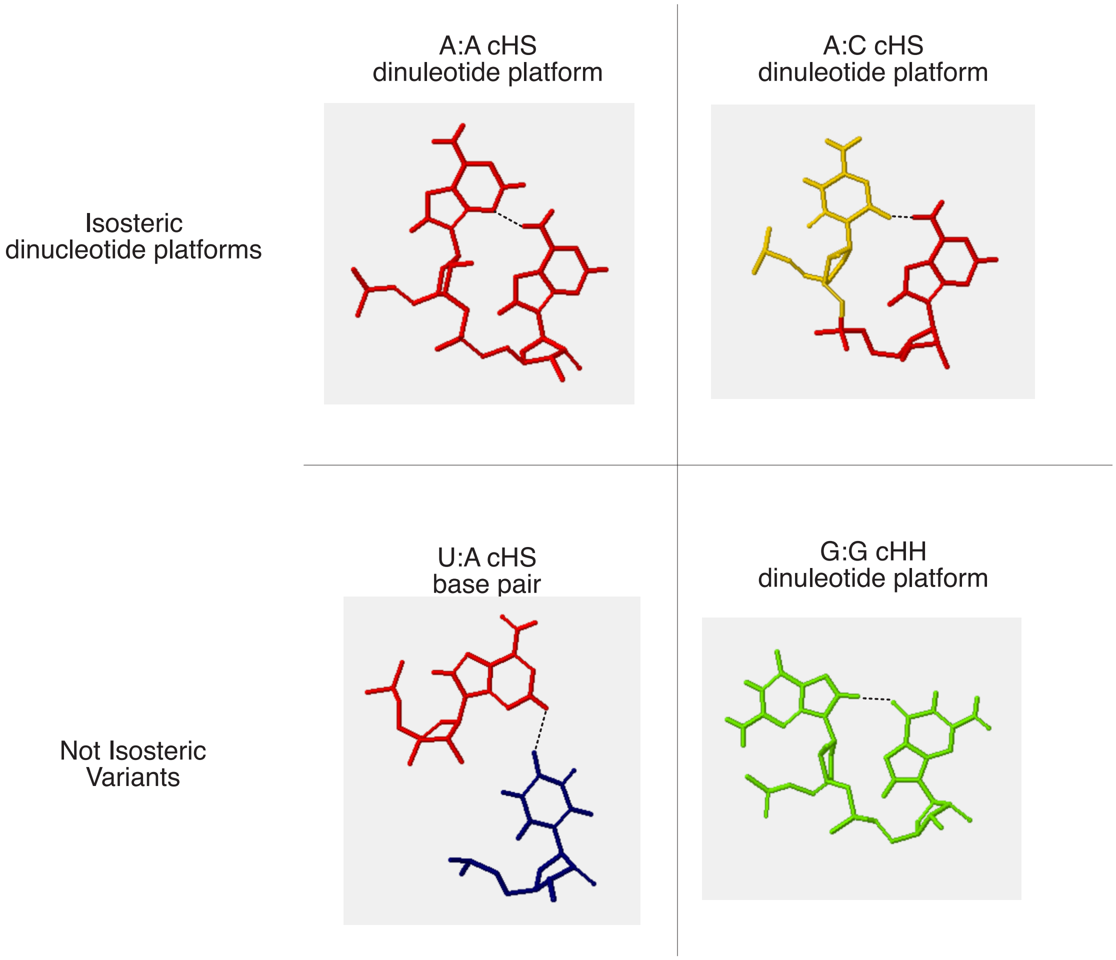


Figure S11: Isosteric structures of dinucleotide platform pairs and tested sequence variants.

#### Figure S12. Summary of Ab Initio Predictions from Different Models

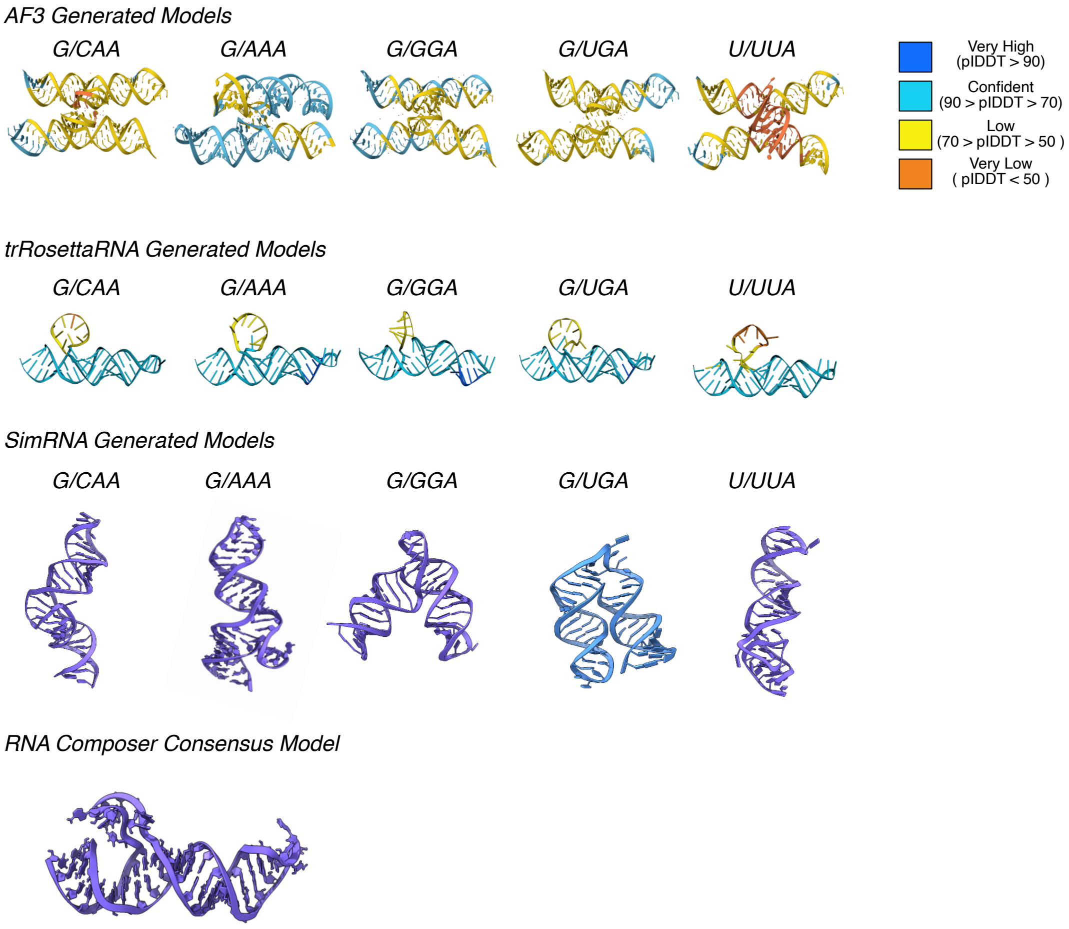


Figure S12: Ab Initio Folding Results from Different Models.

**Figure S13. Sequence motif for alphaKL**

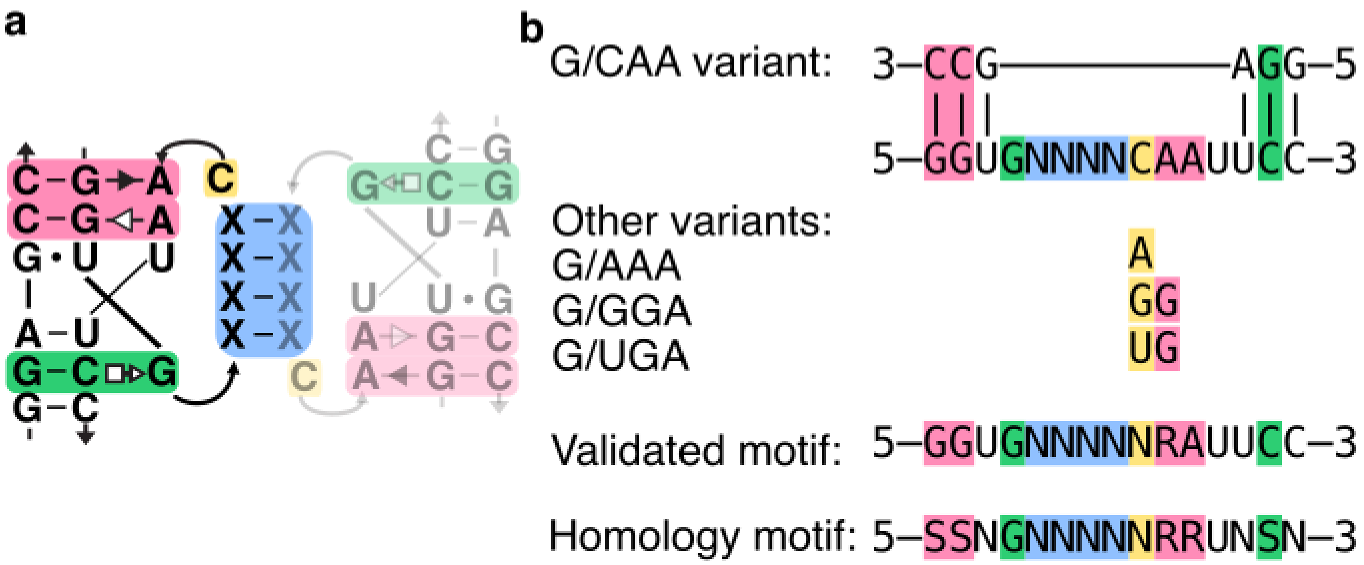


Figure S13: Summary of sequences tested in this report and the overall motif of sequences capable of forming the alphaKL structure. **a** 2D diagram showing the strand routing and intermolecular interactions in the alphaKL. **b** Flattened 2D diagram showing the sequences tested in this report colored based on **a**. The 'validated motif' is the consensus motif of structures experimentally validated here. The 'homology motif' includes additional regions of sequence flexibility suggested by naturally-occurring ribosomal alphaPK sequences.

Validated motif:

`GGUGNNNNNRAUUCC`

Homology motif:

`SSNGNNNNNRRUNSN`

#### Figure S14. Additional AFM images of fibrils

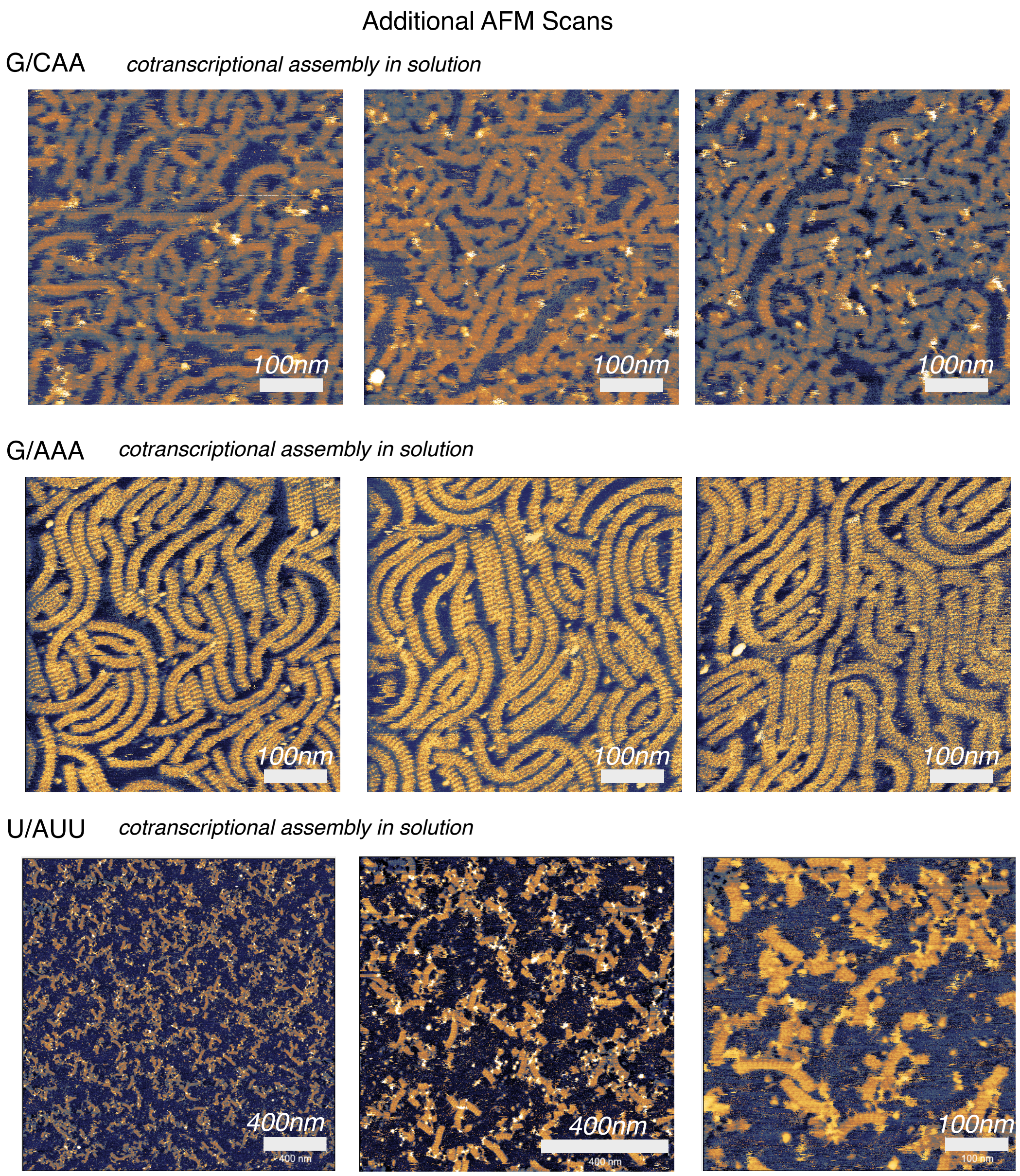


Figure S14: Additional 500 nm scans of G/AAA G/CAA and U/AUU fibrils after cotranscriptional assembly in solution.

**Figure S15. RNA fiber length measurement**

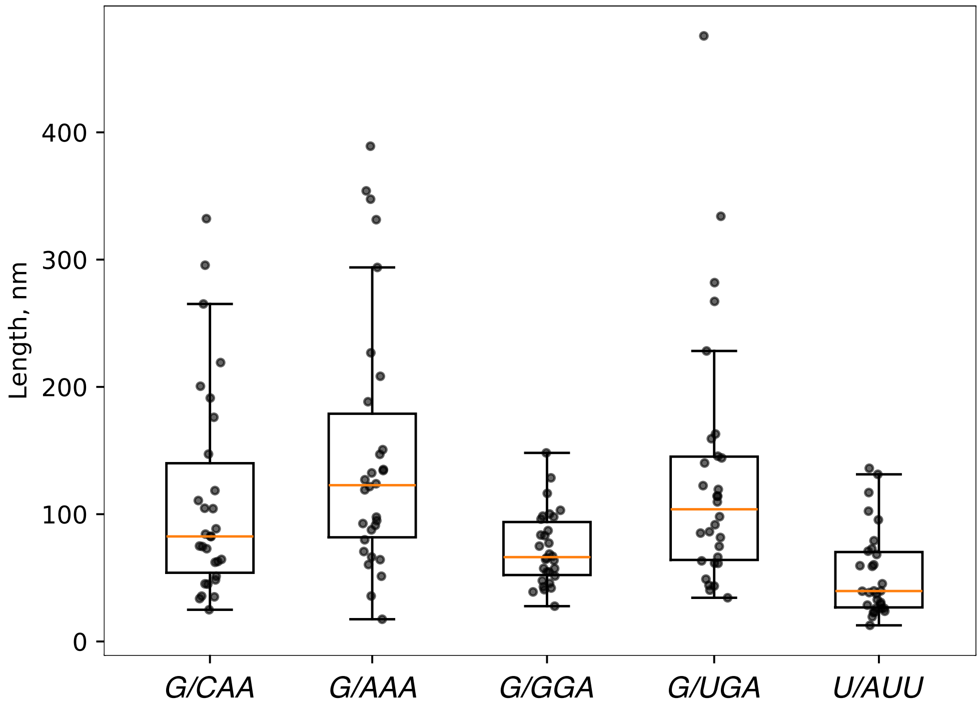


Figure S15: Measurements of G/AAA, G/CAA, U/AUU, G/UGA, and G/GGA fiber lengths in nm after cotranscriptional assembly in solution.

**G/AAA:** $N = 30$, mean = 148, median = 123, std. = 99.4
**G/CAA:** $N = 30$, mean = 111, median = 82.5, std. = 80.3
**U/AUU:** $N = 30$, mean = 53.1, median = 39.6, std. = 33.6
**G/UGA:** $N = 30$, mean = 130, median = 104, std. = 97.4
**G/GGA:** $N = 30$, mean = 72.8, median = 66.3, std. = 28.2

Pairwise comparisons (Mann-Whitney U, Bonferroni-adjusted):

| | | |
|---|---|---|
| G/AAA vs. G/CAA | $p_{adj} = 0.594$ | ns |
| G/AAA vs. U/AUU | $p_{adj} = 3.09 \times 10^{-5}$ | *** |
| G/AAA vs. G/UGA | $p_{adj} = 1$ | ns |
| G/AAA vs. G/GGA | $p_{adj} = 3.37 \times 10^{-3}$ | ** |
| G/CAA vs. U/AUU | $p_{adj} = 3.01 \times 10^{-3}$ | ** |
| G/CAA vs. G/UGA | $p_{adj} = 1$ | ns |
| G/CAA vs. G/GGA | $p_{adj} = 1$ | ns |
| U/AUU vs. G/UGA | $p_{adj} = 8.29 \times 10^{-5}$ | *** |
| U/AUU vs. G/GGA | $p_{adj} = 4.23 \times 10^{-2}$ | * |
| G/UGA vs. G/GGA | $p_{adj} = 8.31 \times 10^{-2}$ | ns |

#### Figure S16. Measurement of alphaKL fiber periodicity

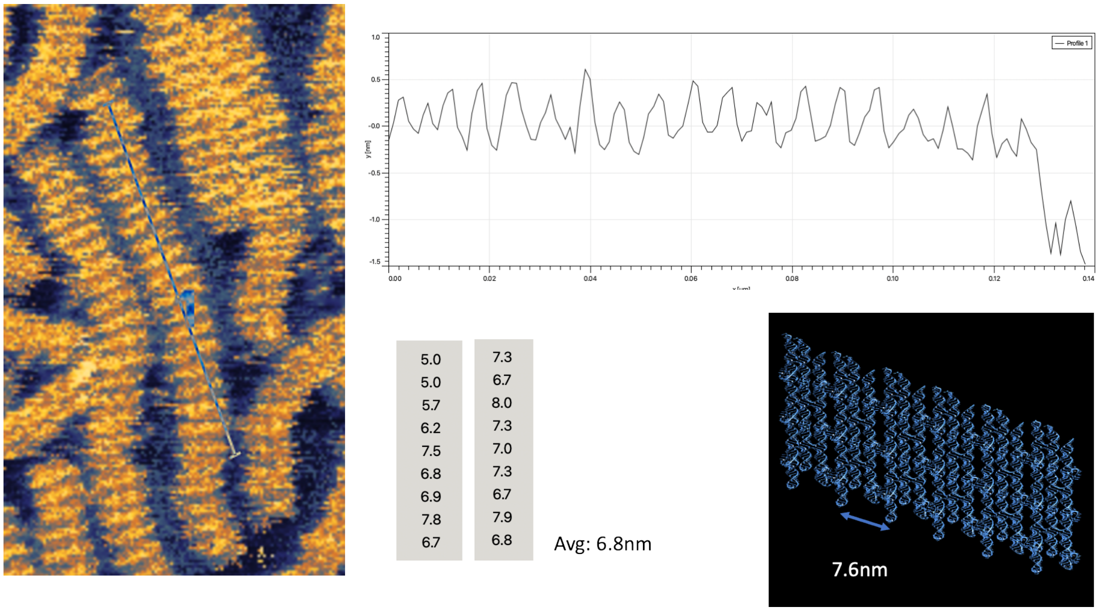


Figure S16: AFM height profile of G/AAA alphaKL fibrils revealing an average periodocity of 6.8 nm.

**Figure S17. Native gel electrophoresis**

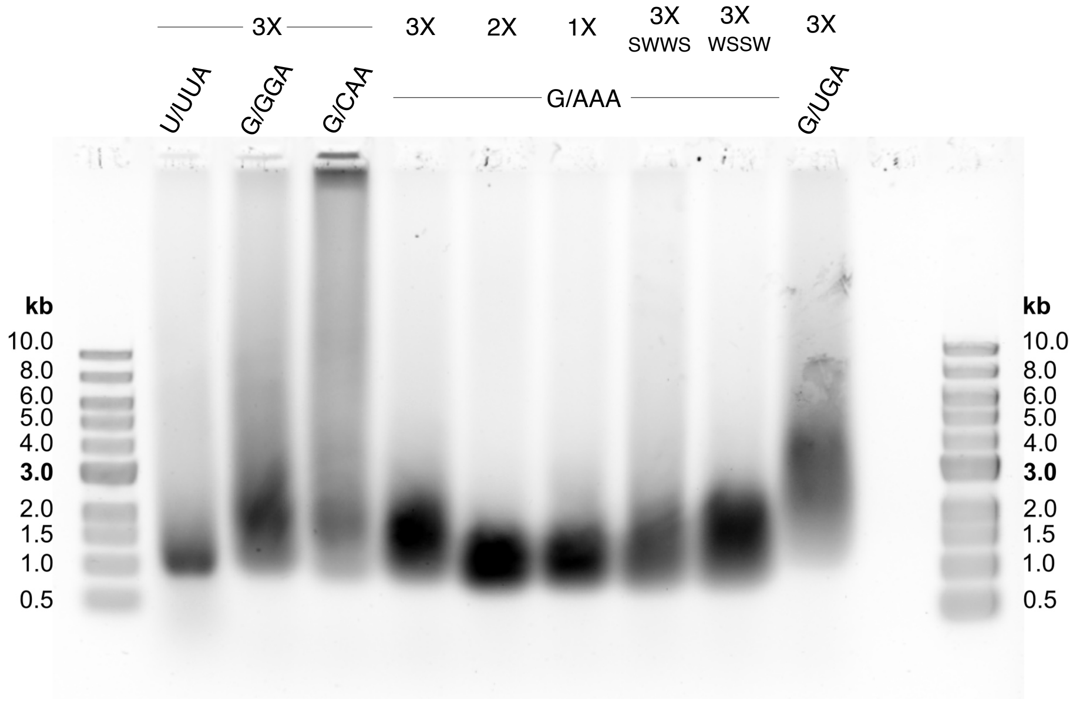


Figure S17: Native gel 0.5% agarose, 0.5X TAE 6 mM $MgCl^{2+}$ run at 60V for 3h on ice. All samples were treated with DNAse I-XT (2% v/v) for 30 min prior to imaging and gel electrophoresis.

**Figure S18. MD simulation of minimal alphaKL motifs**

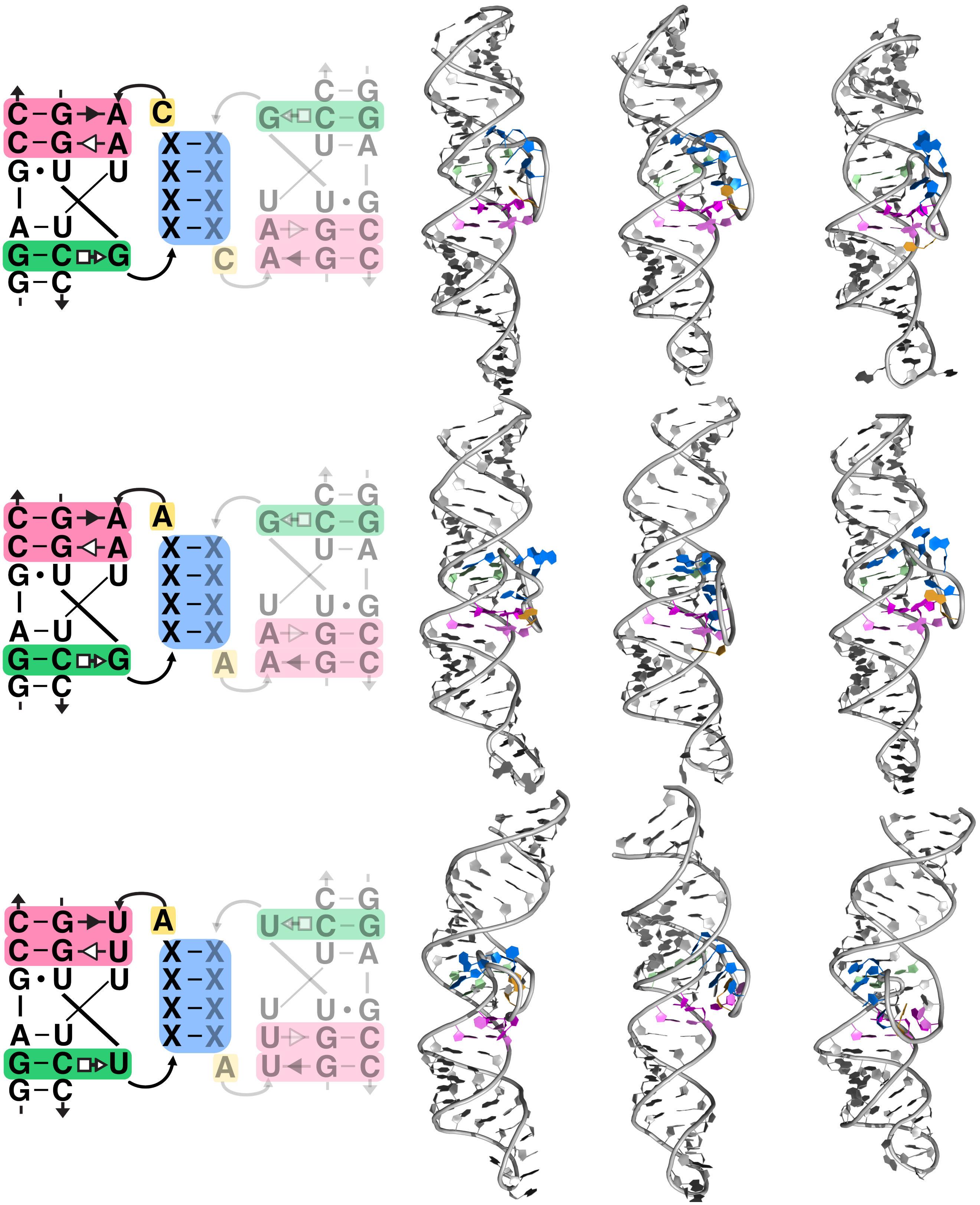

Figure S18: Each row shows the sequence and routing of one variant (Top: G/CAA, Middle: G/AAA, Bottom: U/AUU). The structure renderings show the final configuration after 2 µs of equilibrium MD simulation.

**Figure S19. MD simulation of minimal alphaKL motifs**

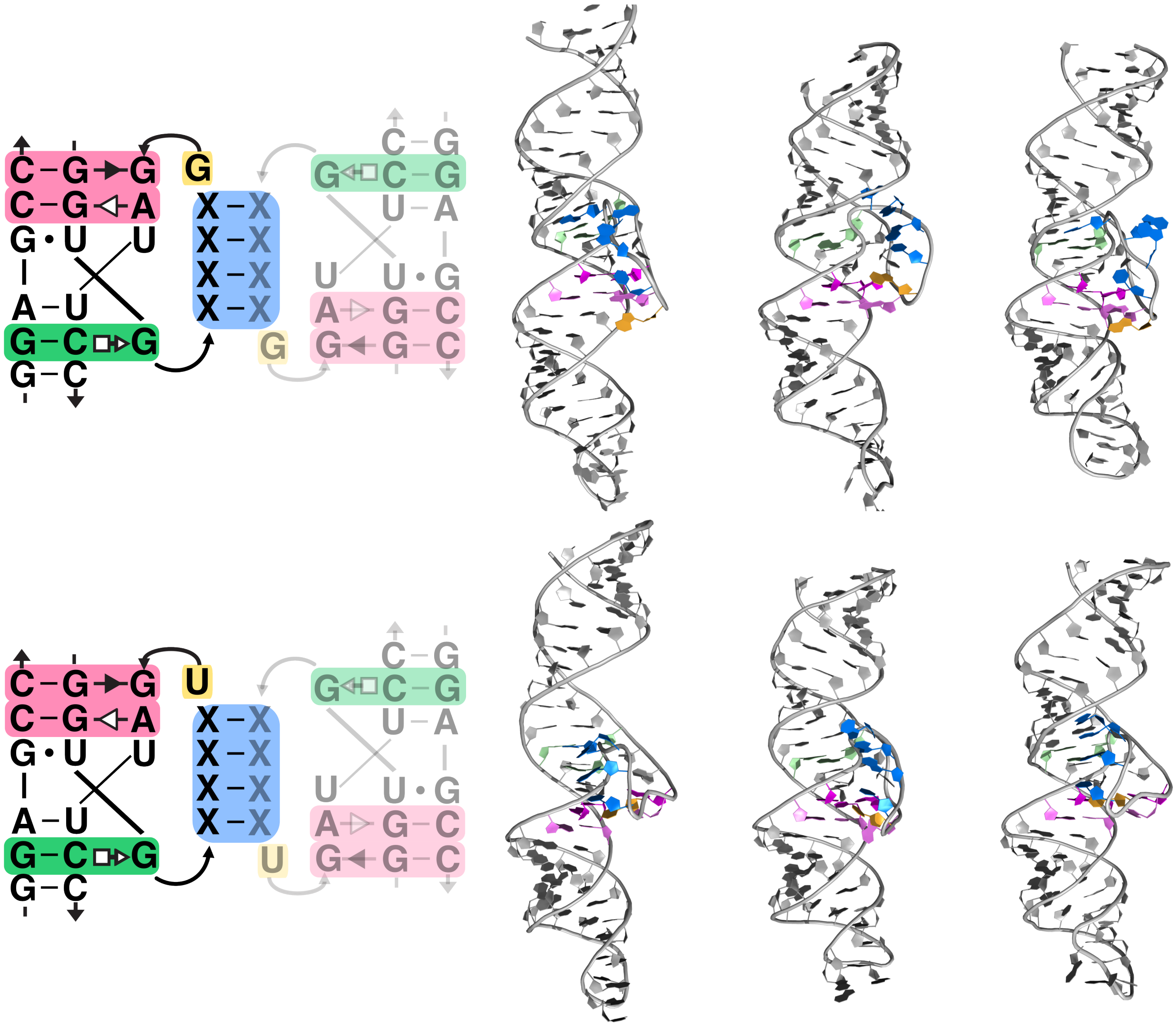

Figure S19: Each row shows the sequence and routing of one variant (Top: G/GGA, Bottom: G/UGA). The structure renderings show the final configuration after 2 µs of equilibrium MD simulation.

**Figure S20. Detailed view of minimal alphaKL motifs**

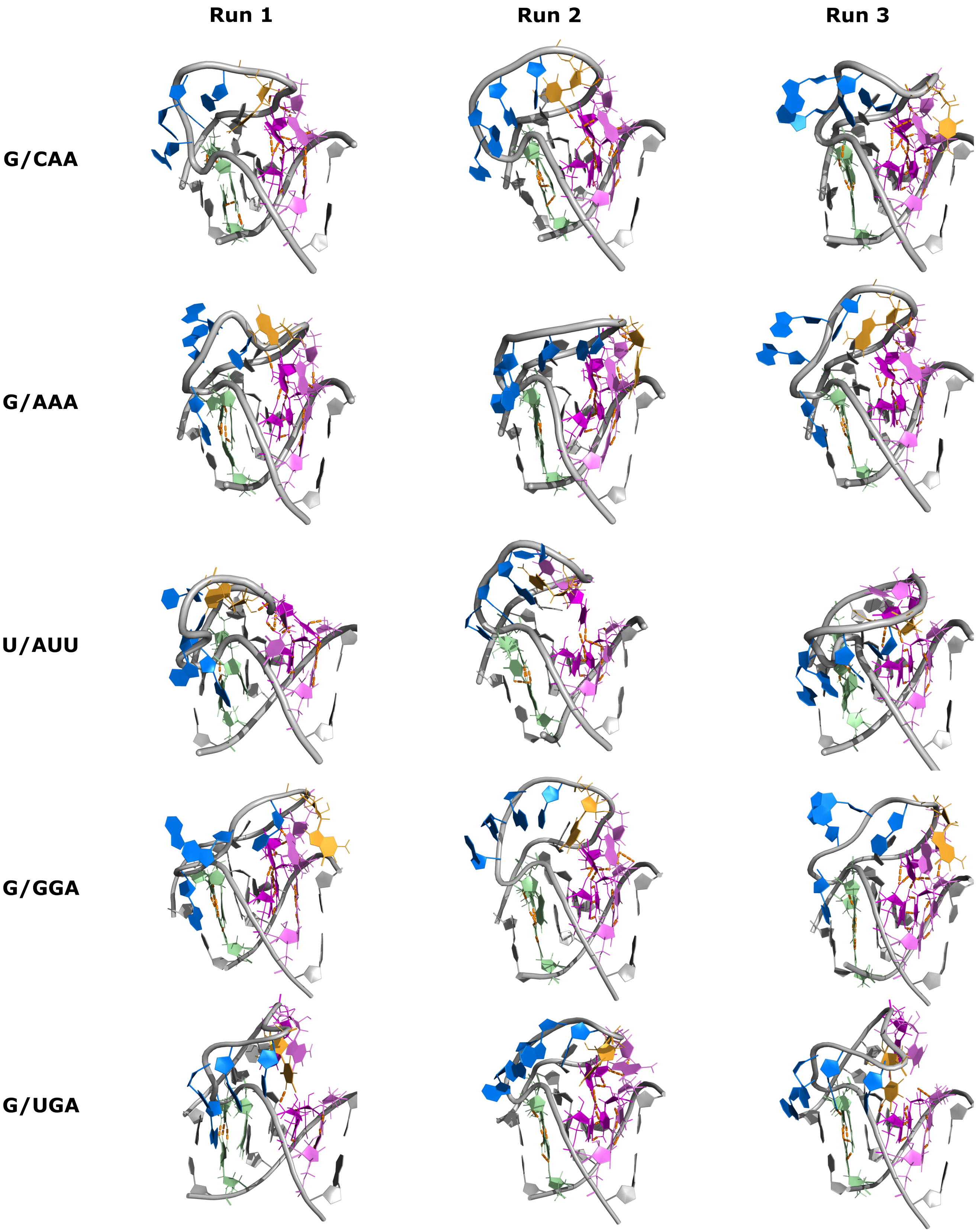


Figure S20: Detailed views of the alphaKL structures from figure S18 & S19 showing 5' triplet (green), binding interface (blue), platform nucleotide (yellow), 3' triplets (pink), and triplet hydrogen bonds (orange).

**Figure S21. MD simulation of minimal alphaKL dimers**

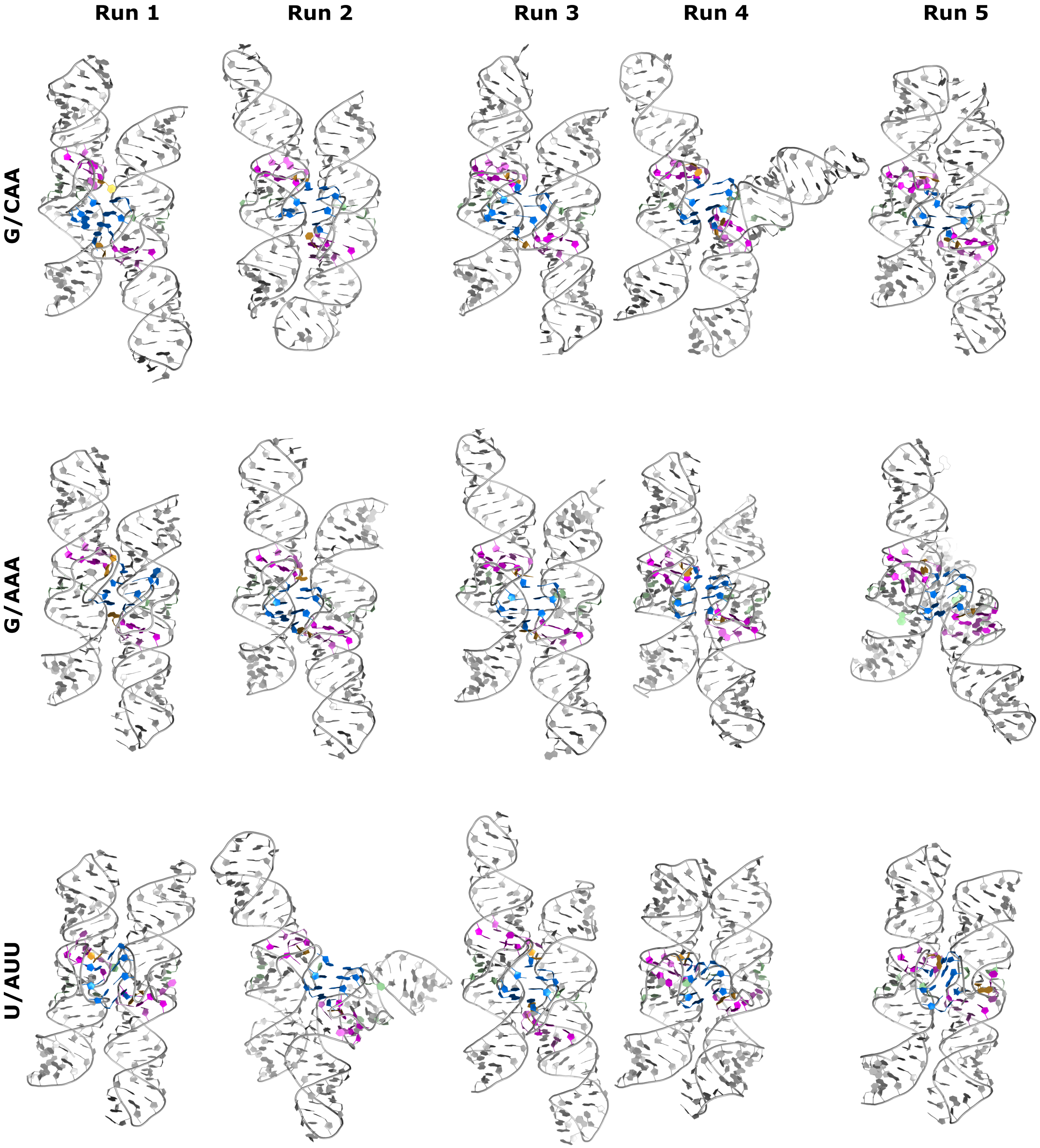


Figure S21: MD simulations of dimers of the minimal alphaKL motif. Each row shows the sequence and routing of one variant (Top: G/AAA, Middle: G/CAA, Bottom: U/AUU). The structure renderings show the final configuration after 2 µs of equilibrium MD simulation.

**Figure S22. MD simulation of minimal alphaKL dimers**

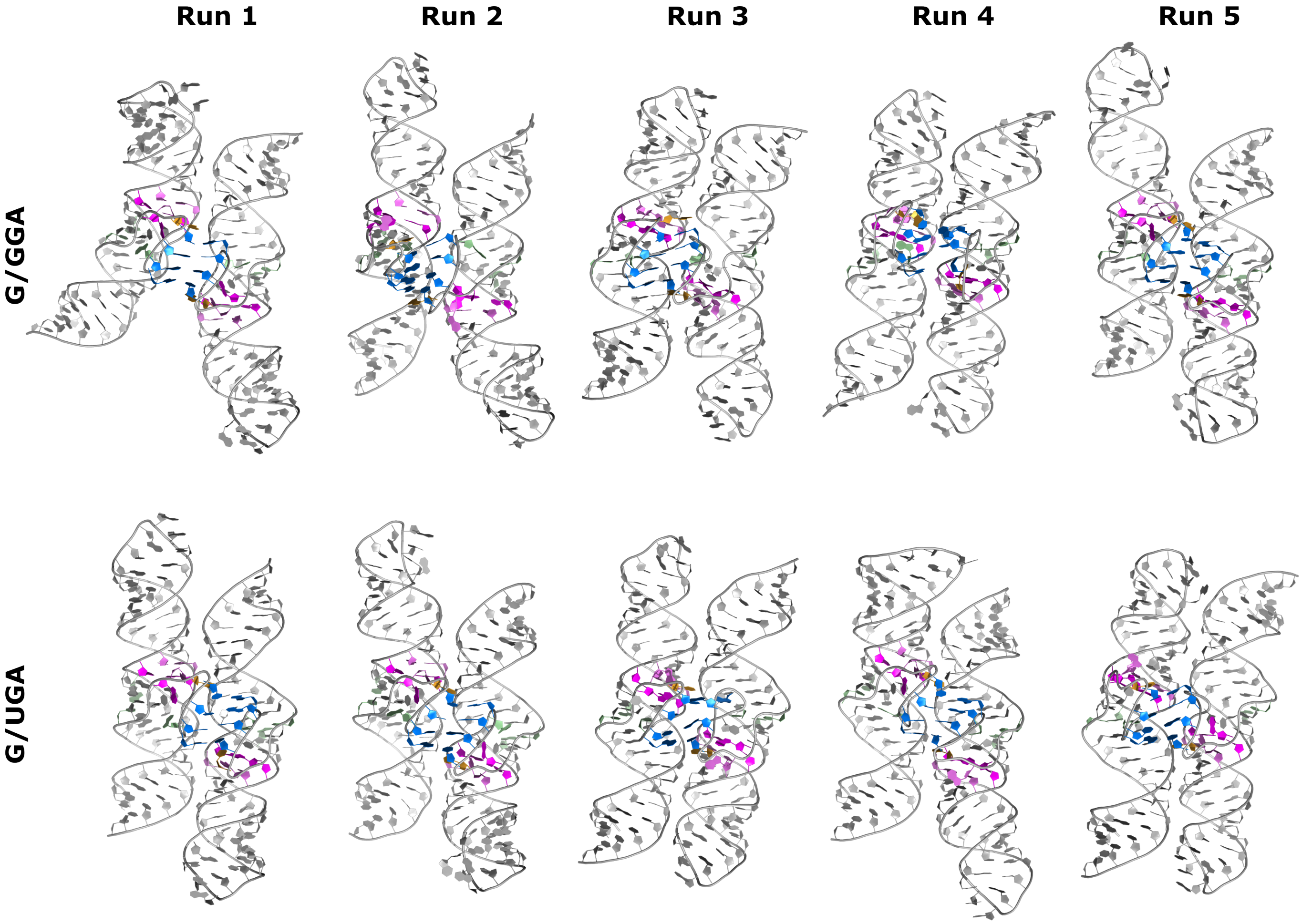


Figure S22: MD simulations of dimers of the minimal alphaKL motif. Each row shows the sequence and routing of one variant (Top: G/GGA, Bottom: G/UGA). The structure renderings show the final configuration after 2 µs of equilibrium MD simulation.

**Figure S23. Detailed view of minimal alphaKL dimers**

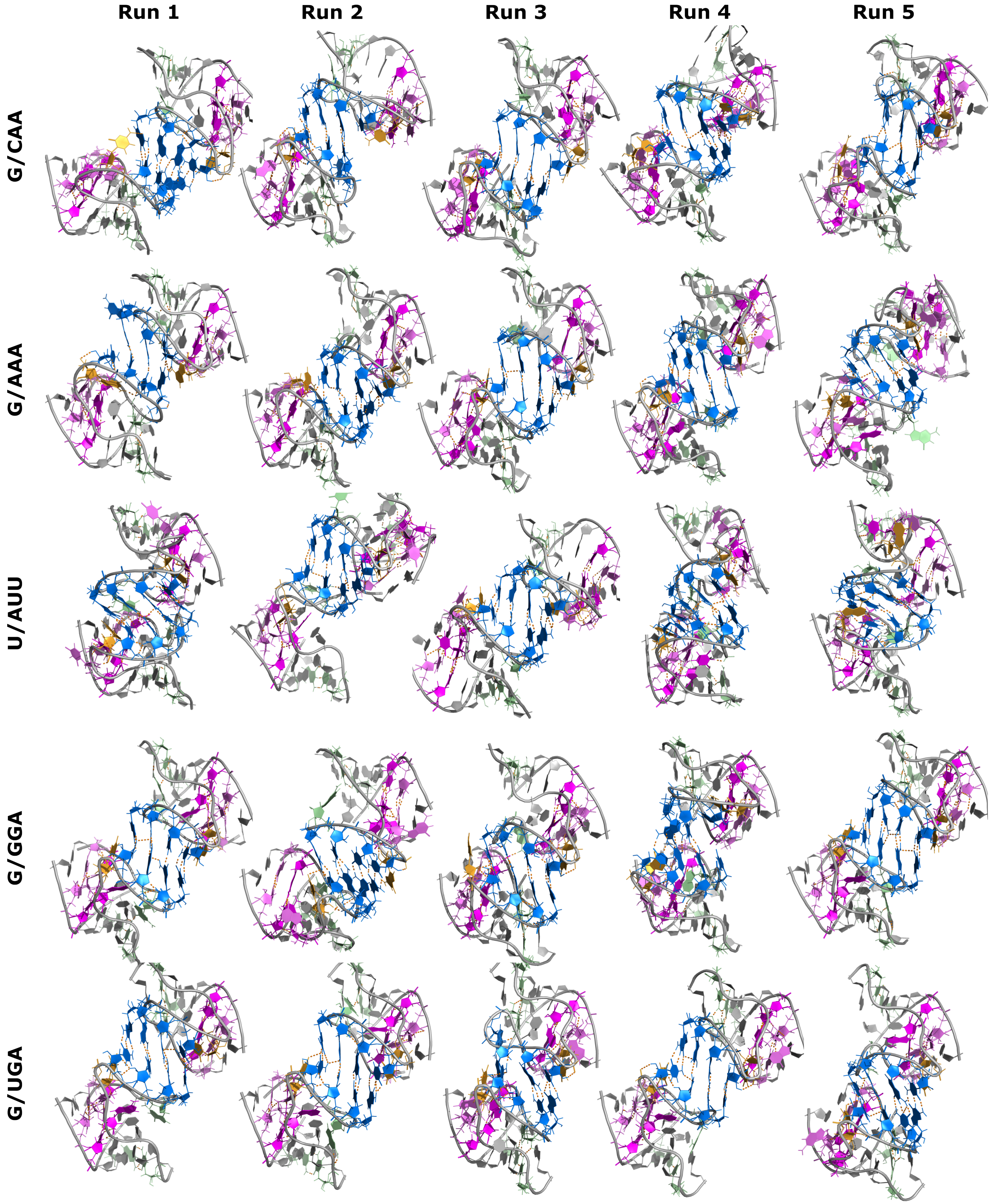


Figure S23: Detailed views of the alphaKL dimers from figure S21 & S22 showing 5' triplet (green), binding interface (blue), platform nucleotide (yellow), 3' triplets (pink), and triplet hydrogen bonds (orange).

**Figure S24. MD simulation of RNA origami assemblies**

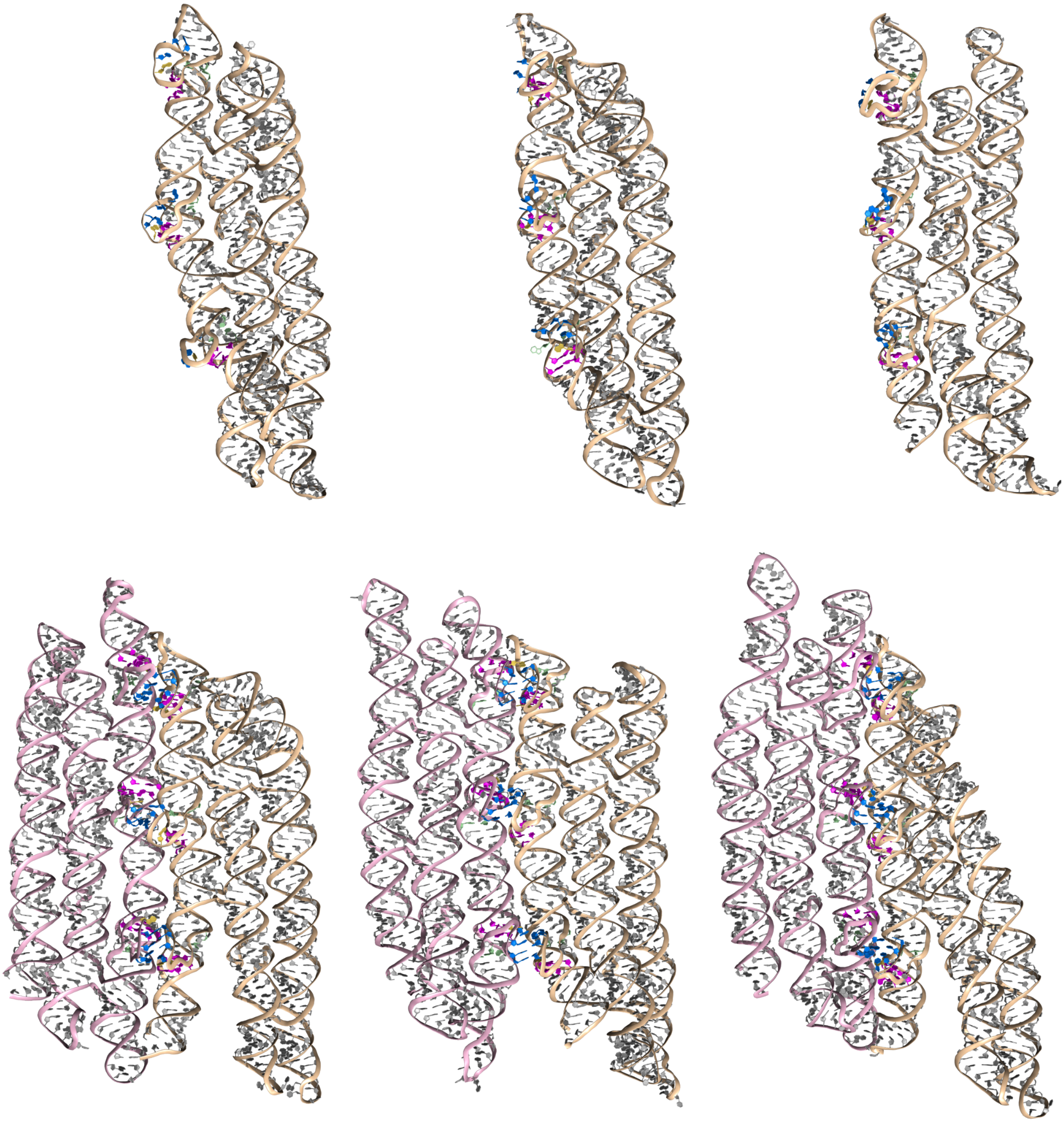

Figure S24: MD simulations of 3-helix tiles which connect via G/AAA alphaKLs as both monomers and dimers. Each structures shows the final configuration after 500 ns of equilibrium MD simulation. The color scheme of the nucleotides matches the minimal motifs.

**Figure S25. Hydrogen bonds in monomers**

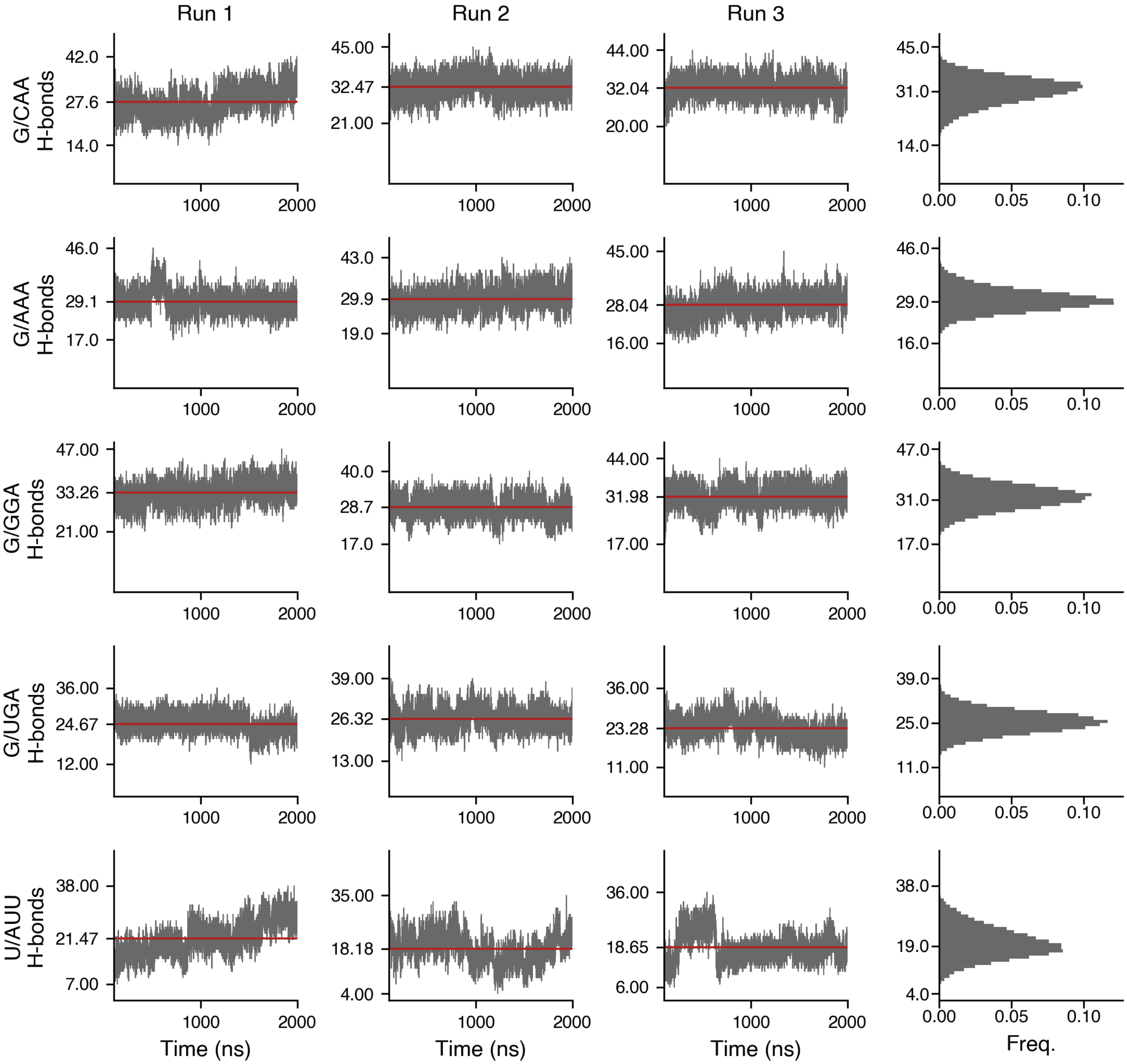


Figure S25: Number of hydrogen bonds in simulations of a monomer of the minimal alphaKL motif. The Y-axis ticks correspond to the minimum, mean and maximum number of hydrogen bonds in each trajectory.

**Figure S26. Percentage of monomer simulation frames with triplets formed**

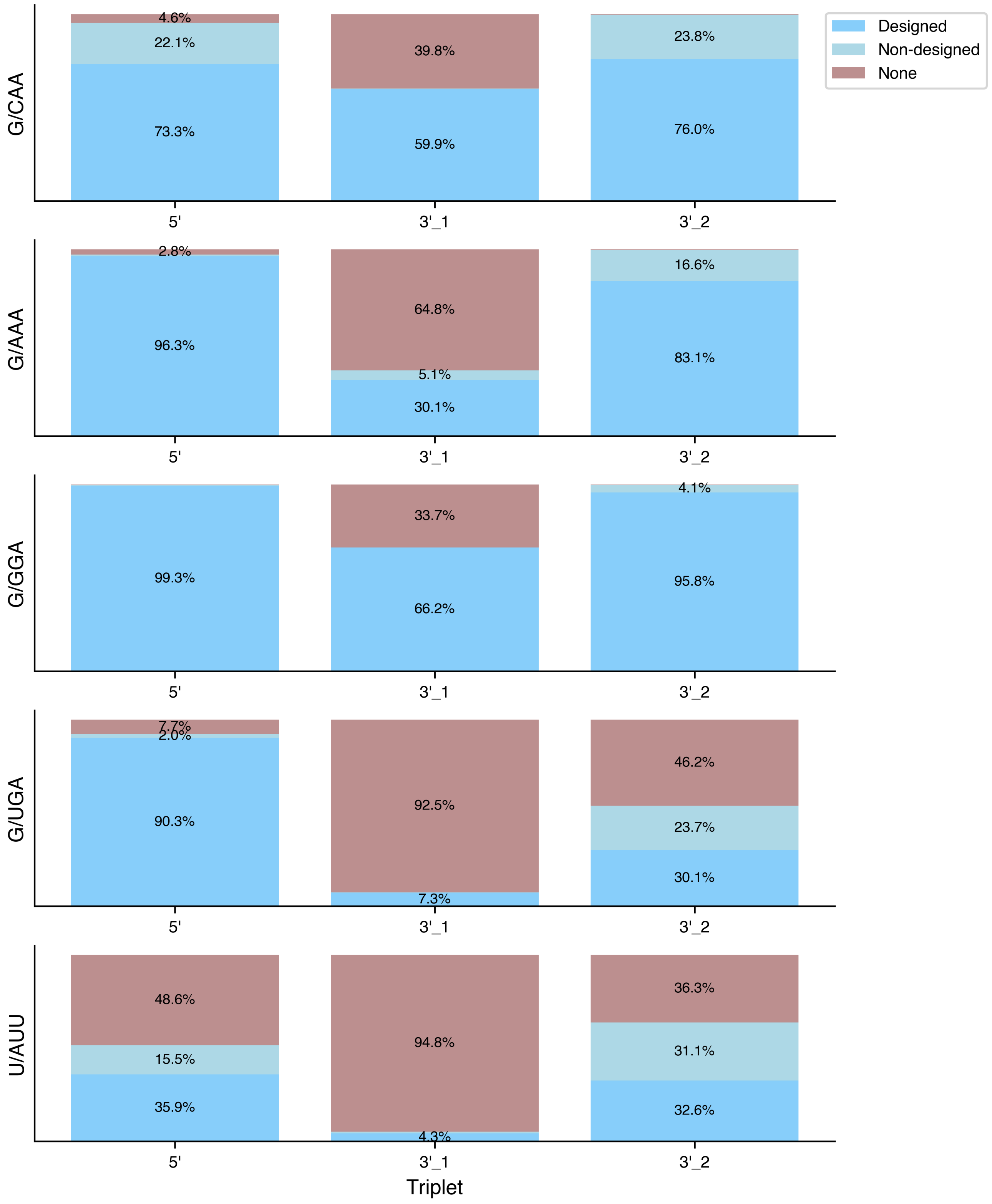


Figure S26: Percentage of frames where the designed triplet nucleotides participate in triplet hydrogen-bonding interaction during monomer simulations.

Table S1: Triplet occupancy for the G/CAA monomer

| | Triplet | Non-triplet |
|---|---|---|
| 5’ | 95.4 | 4.6 |
| 3’_1 | 60.2 | 39.8 |
| 3’_2 | 99.8 | 0.2 |

Table S2: Triplet occupancy for the G/AAA monomer

| | Triplet | Non-triplet |
|---|---|---|
| 5’ | 97.2 | 2.8 |
| 3’_1 | 35.2 | 64.8 |
| 3’_2 | 99.7 | 0.3 |

Table S3: Triplet occupancy for the G/GGA monomer

| | Triplet | Non-triplet |
|---|---|---|
| 5’ | 99.9 | 0.1 |
| 3’_1 | 66.3 | 33.7 |
| 3’_2 | 99.9 | 0.1 |

Table S4: Triplet occupancy for the G/UGA monomer

| | Triplet | Non-triplet |
|---|---|---|
| 5’ | 92.3 | 7.7 |
| 3’_1 | 7.5 | 92.5 |
| 3’_2 | 53.8 | 46.2 |

Table S5: Triplet occupancy for the U/AUU monomer

| | Triplet | Non-triplet |
|---|---|---|
| 5’ | 51.4 | 48.6 |
| 3’_1 | 5.2 | 94.8 |
| 3’_2 | 63.7 | 36.3 |

**Figure S27. Number of homomeric H-bonds in dimers**

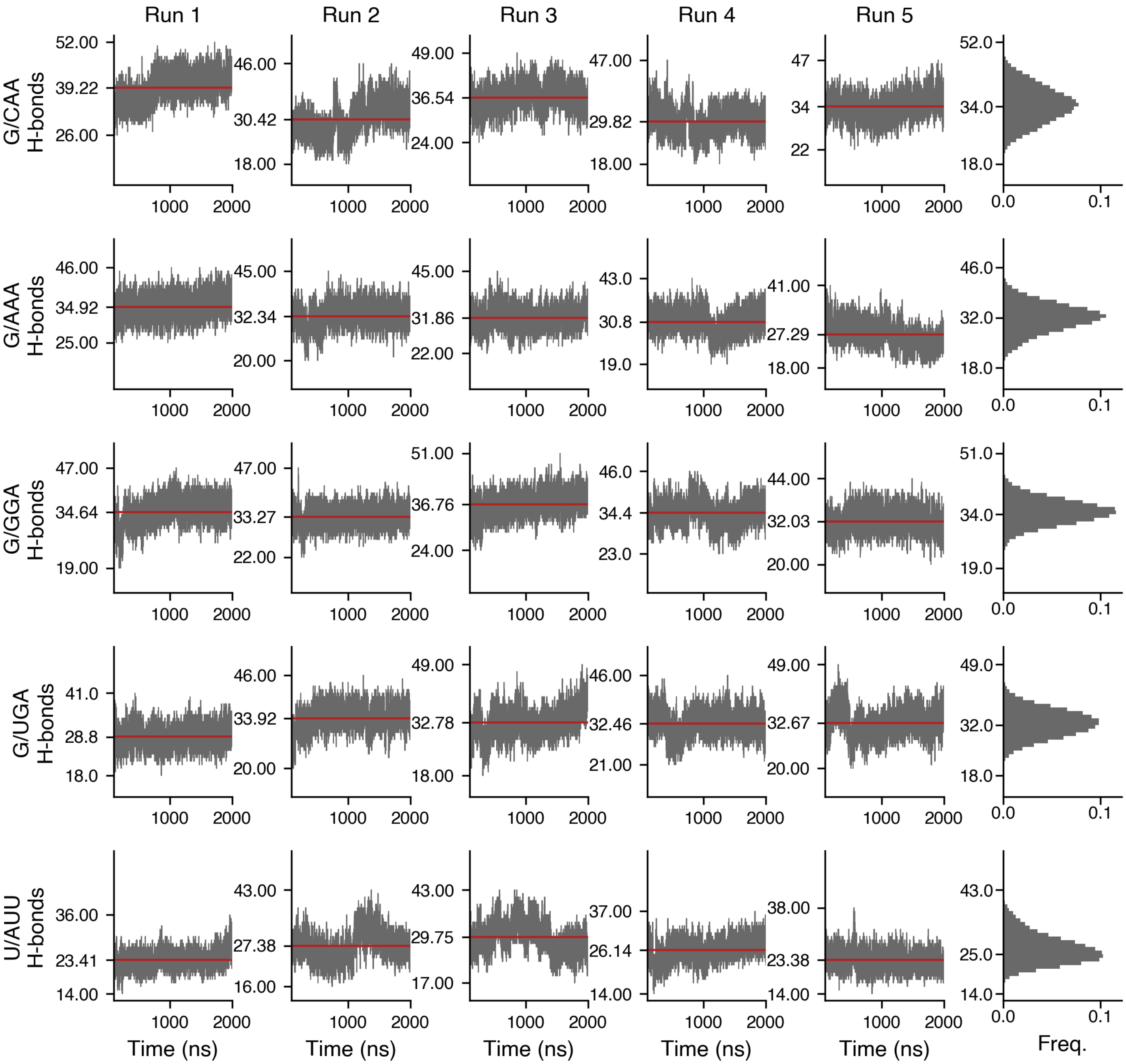


Figure S27: Number of homomeric hydrogen bonds in one monomer during simulations of the minimal alphaKL dimer. The Y-axis ticks correspond to the minimum, mean and maximum number of hydrogen bonds in each simulation trajectory. Note that only homomeric bonds were included to make these trajectories directly comparable with Fig. S25

**Figure S28. Percentage of dimer simulation frames with triplets formed**

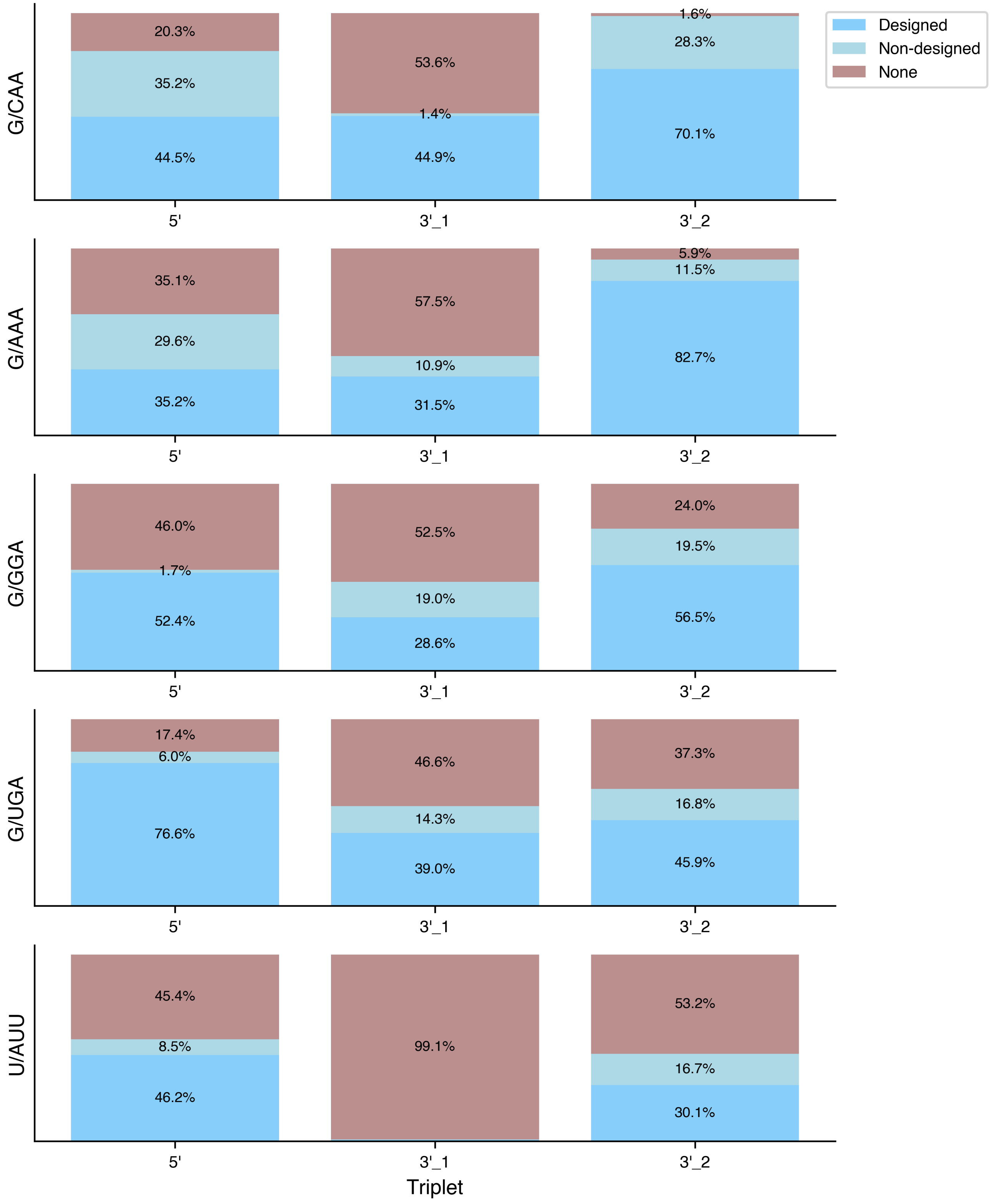


Figure S28: Percentage of frames where the designed triplet nucleotides participate in triplet hydrogen-bonding interaction during dimer simulations.

Table S6: Triplet occupancy for the G/CAA dimer

| | Triplet | Non-triplet |
|---|---|---|
| 5’ | 79.7 | 20.3 |
| 3’_1 | 46.4 | 53.6 |
| 3’_2 | 98.4 | 1.6 |

Table S7: Triplet occupancy for the G/AAA dimer

| | Triplet | Non-triplet |
|---|---|---|
| 5’ | 64.9 | 35.1 |
| 3’_1 | 42.5 | 57.5 |
| 3’_2 | 94.1 | 5.9 |

Table S8: Triplet occupancy for the G/GGA dimer

| | Triplet | Non-triplet |
|---|---|---|
| 5’ | 54.0 | 46.0 |
| 3’_1 | 47.5 | 52.5 |
| 3’_2 | 76.0 | 24.0 |

Table S9: Triplet occupancy for the G/UGA dimer

| | Triplet | Non-triplet |
|---|---|---|
| 5’ | 82.6 | 17.4 |
| 3’_1 | 53.4 | 46.6 |
| 3’_2 | 62.7 | 37.3 |

Table S10: Triplet occupancy for the U/AUU dimer

| | Triplet | Non-triplet |
|---|---|---|
| 5’ | 54.6 | 45.4 |
| 3’_1 | 0.9 | 99.1 |
| 3’_2 | 46.8 | 53.2 |

#### Figure S29. Total number of backbone-backbone H-bonds

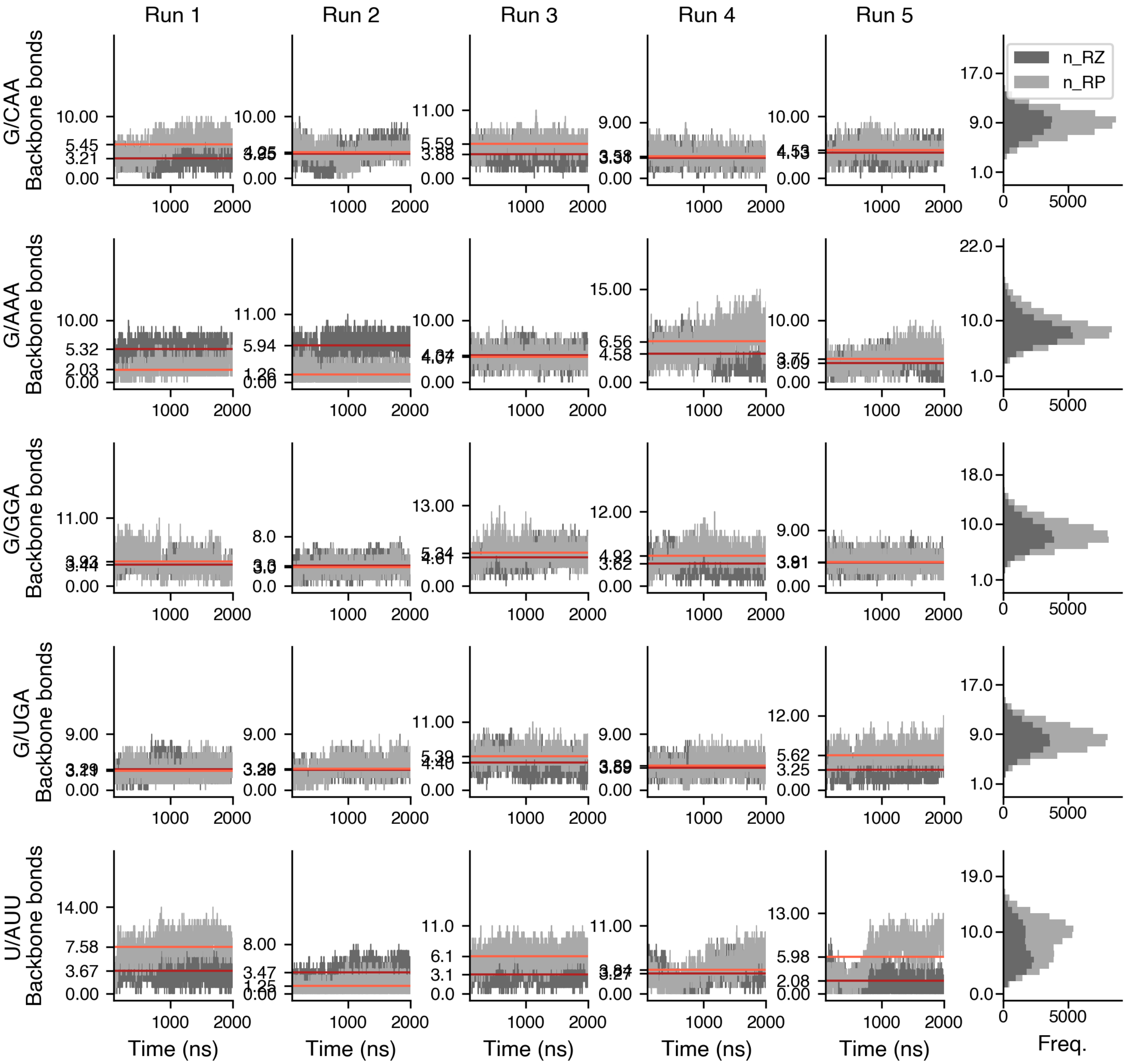


Figure S29: Number of backbone-backbone bonds formed between the two stems during simulations of the minimal alphaKL dimer. The darker color corresponds to ribose-ribose (ribose zipper, RZ) bonds, while the lighter color is ribose-phospate (RP) bonds. The Y-axis ticks correspond to the minimum, mean and maximum number of ribose zippers in each simulation trajectory.

#### Figure S30. Angles between stems

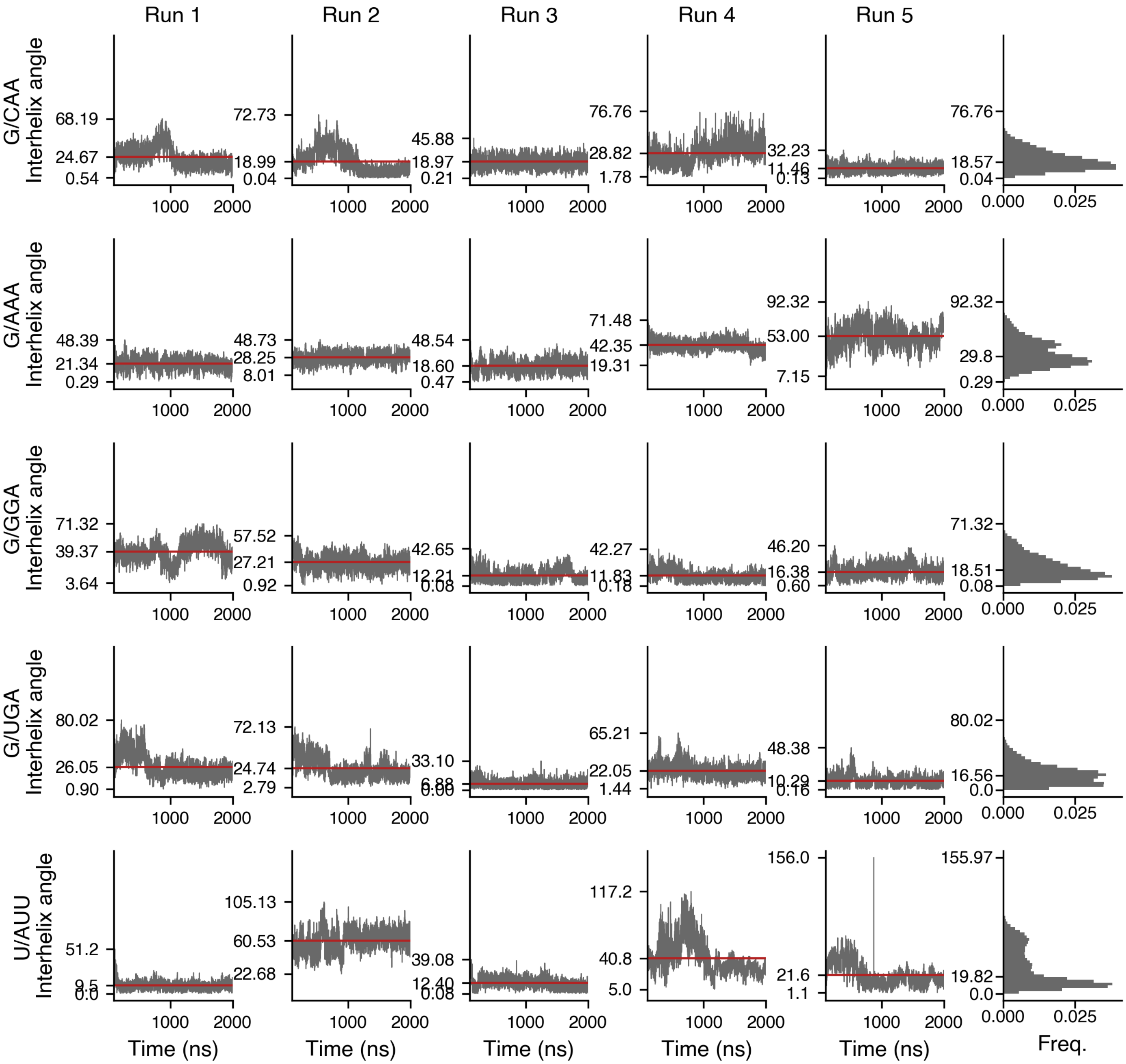


Figure S30: Angles between the two stems during simulations of the minimal alphaKL dimer. The Y-axis ticks correspond to the minimum, mean and maximum angle in each simulation trajectory.

**Figure S31. Formation of designed triplets vs total backbone-backbone H-bonds**

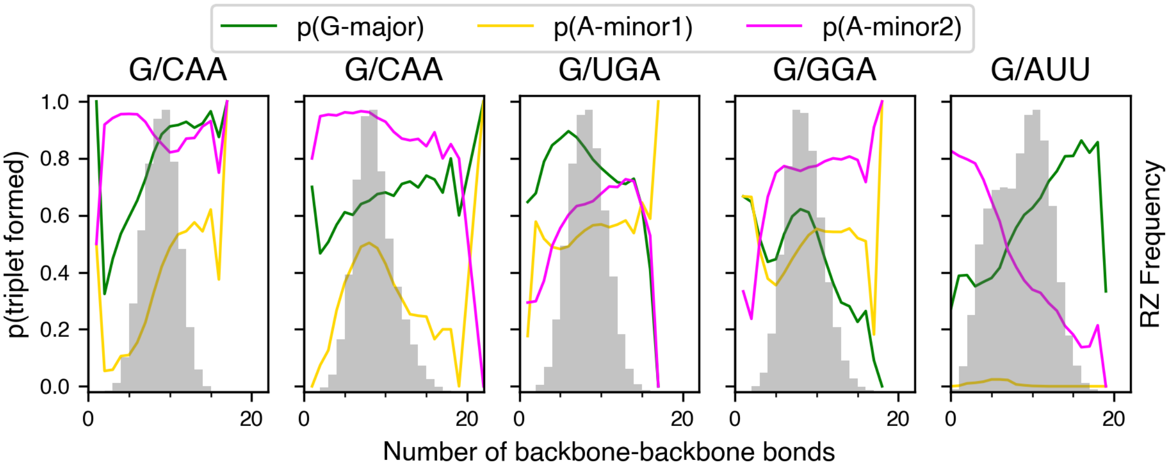


Figure S31: Correlation between the number of backbone-backbone bonds (Fig. S29 and the fraction of frames that each designed triple is formed (Fig. 5b).

**Figure S32. Angle vs number of backbone-backbone H-bonds**

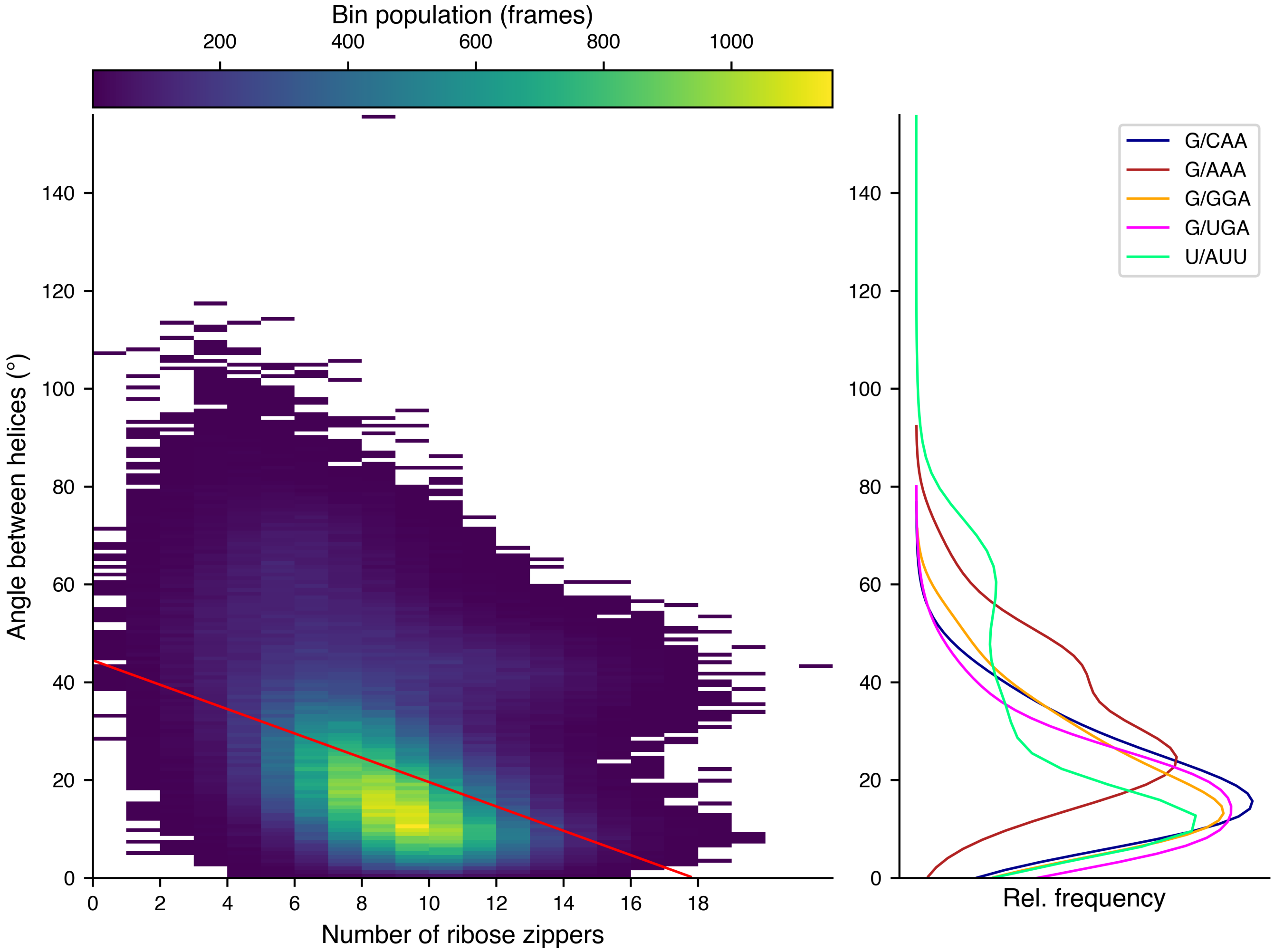


Figure S32: Correlation between the number of backbone-backbone bonds (Fig. S29, smoothed, aggregated data reproduced in the right panel) and the angle between the two stems (Fig. S30). On average, for each additional ribose zipper formed, the angle between the helices decreases by 2.49 degrees (red line).

**Figure S33. Number of homomeric H-bonds in full-tile simulations**

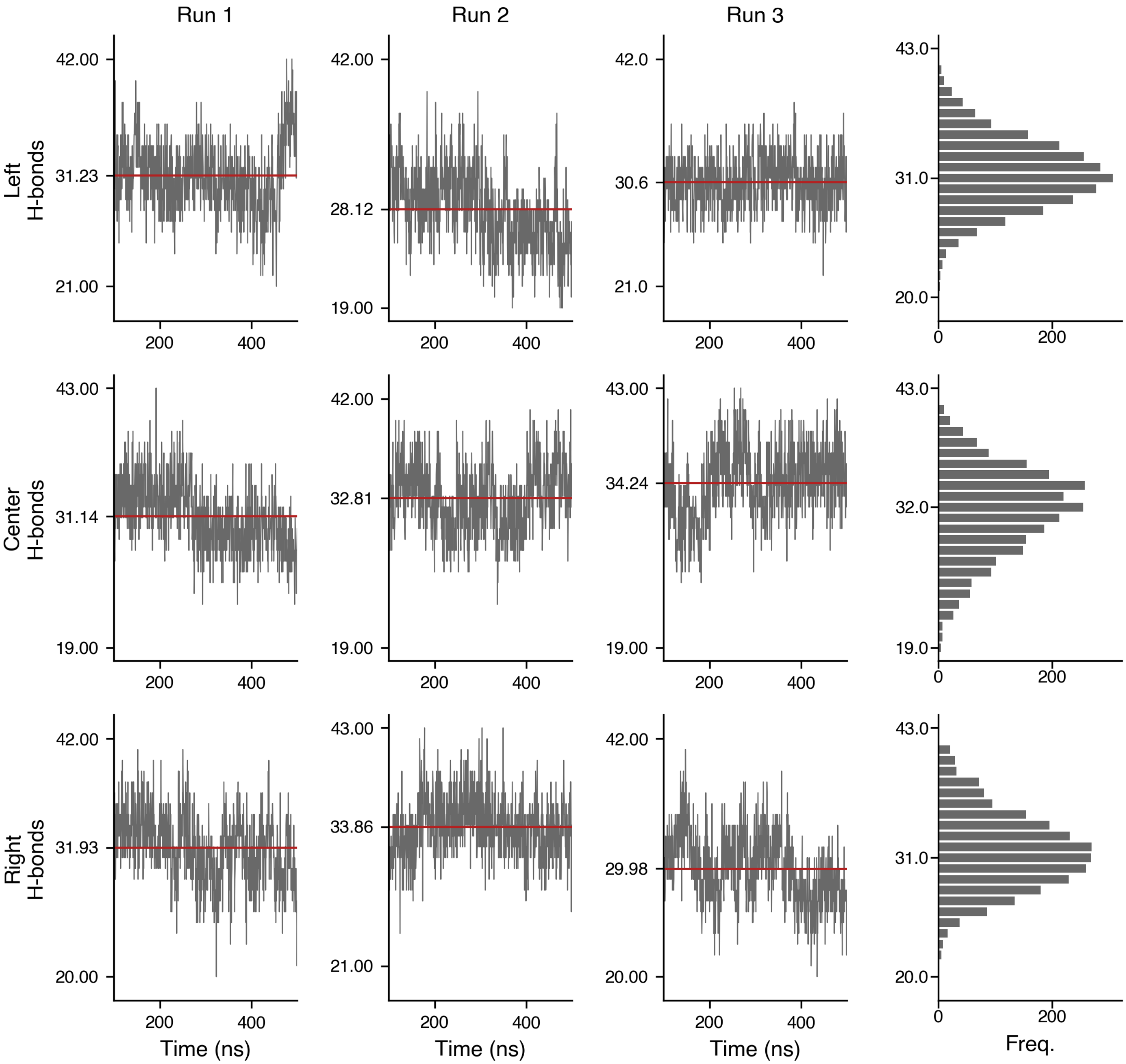


Figure S33: Number of homomeric hydrogen bonds in the three alphaKLs of the "top" tile during simulations of a tile dimer joined by G/AAA-variant alphaKLs. The Y-axis ticks correspond to the minimum, mean and maximum number of hydrogen bonds in each simulation trajectory.

**Figure S34. Backbone-backbone H-bonds in full-tile simulations**

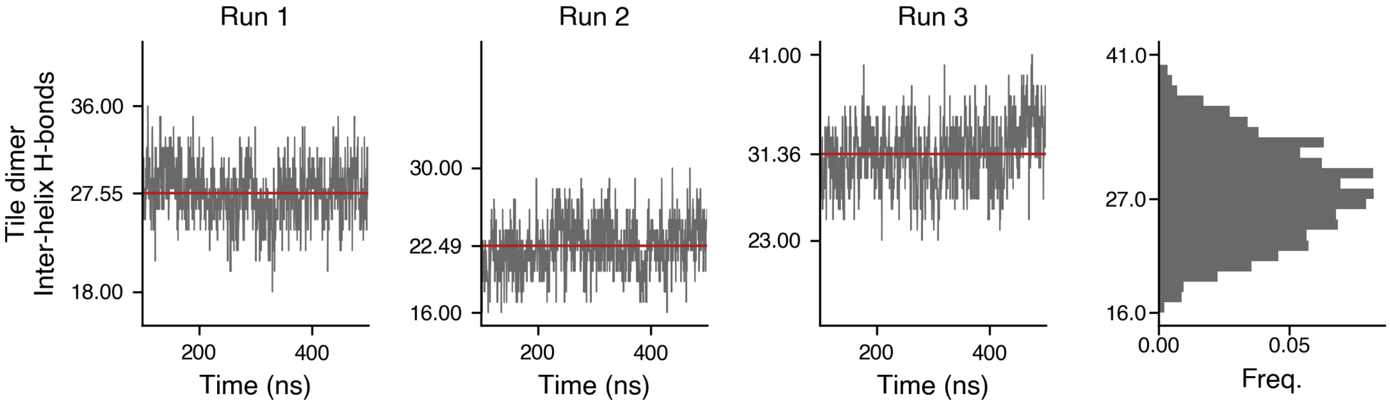


Figure S34: The number of backbone-backbone H-bonds between in the tiles in the two-tile simulations.

**Figure S35. Ribose zipper and ribose phosphate interactions**

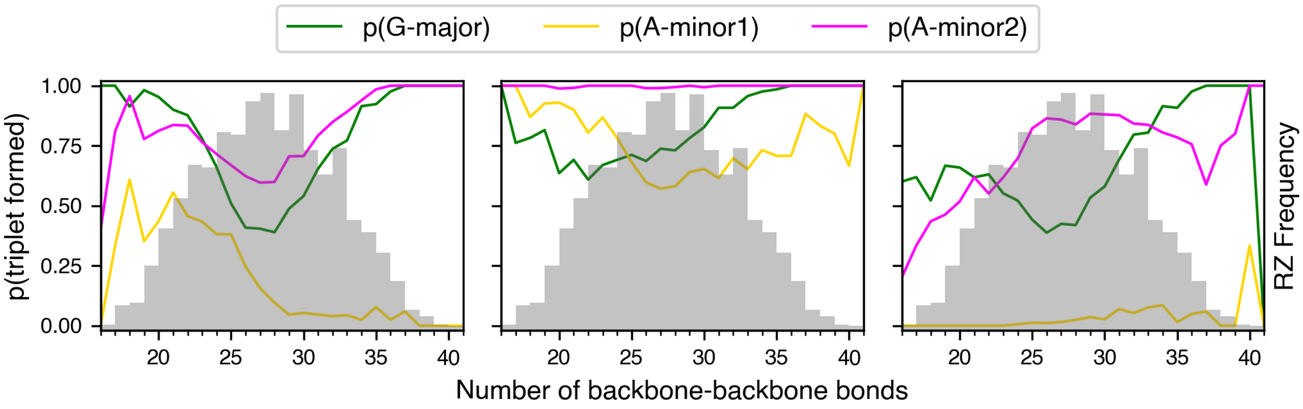


Figure S35: Correlation between the number of backbone-backbone H-bonds (Fig. S34) and the fraction of frames that each designed platform bond is formed (Fig. 5c).

**Figure S36. RMSD initial monomer model vs simulation trajectory**

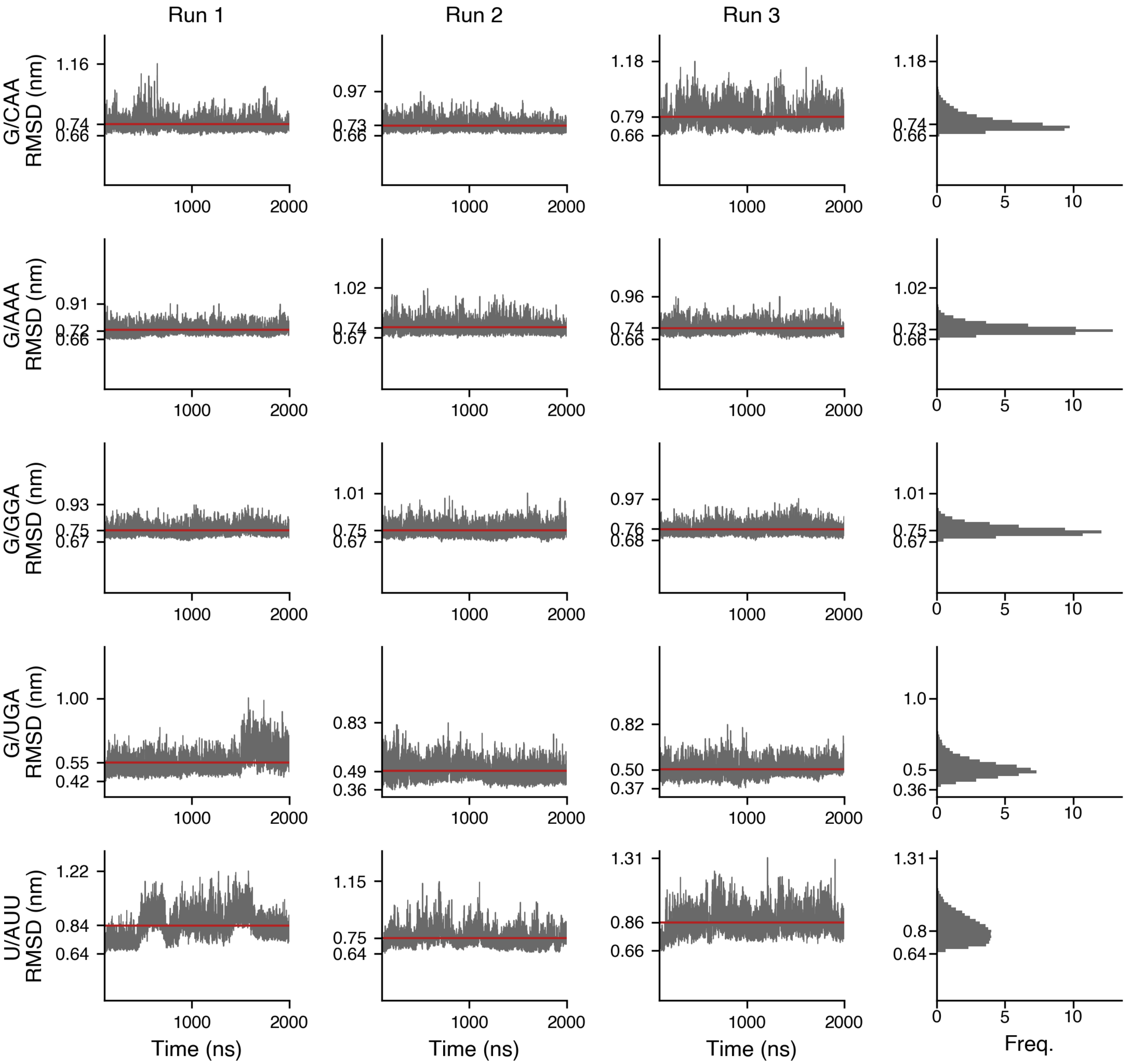


Figure S36: Root mean squared displacement (RMSD) vs the initial model used to start the MD simulations for simulations of a monomer of the minimal alphaKL motif. The Y-axis ticks correspond to the minimum, mean and maximum RMSD value in each simulation trajectory.

**Figure S37. RMSD of initial dimer model vs simulation trajectory**

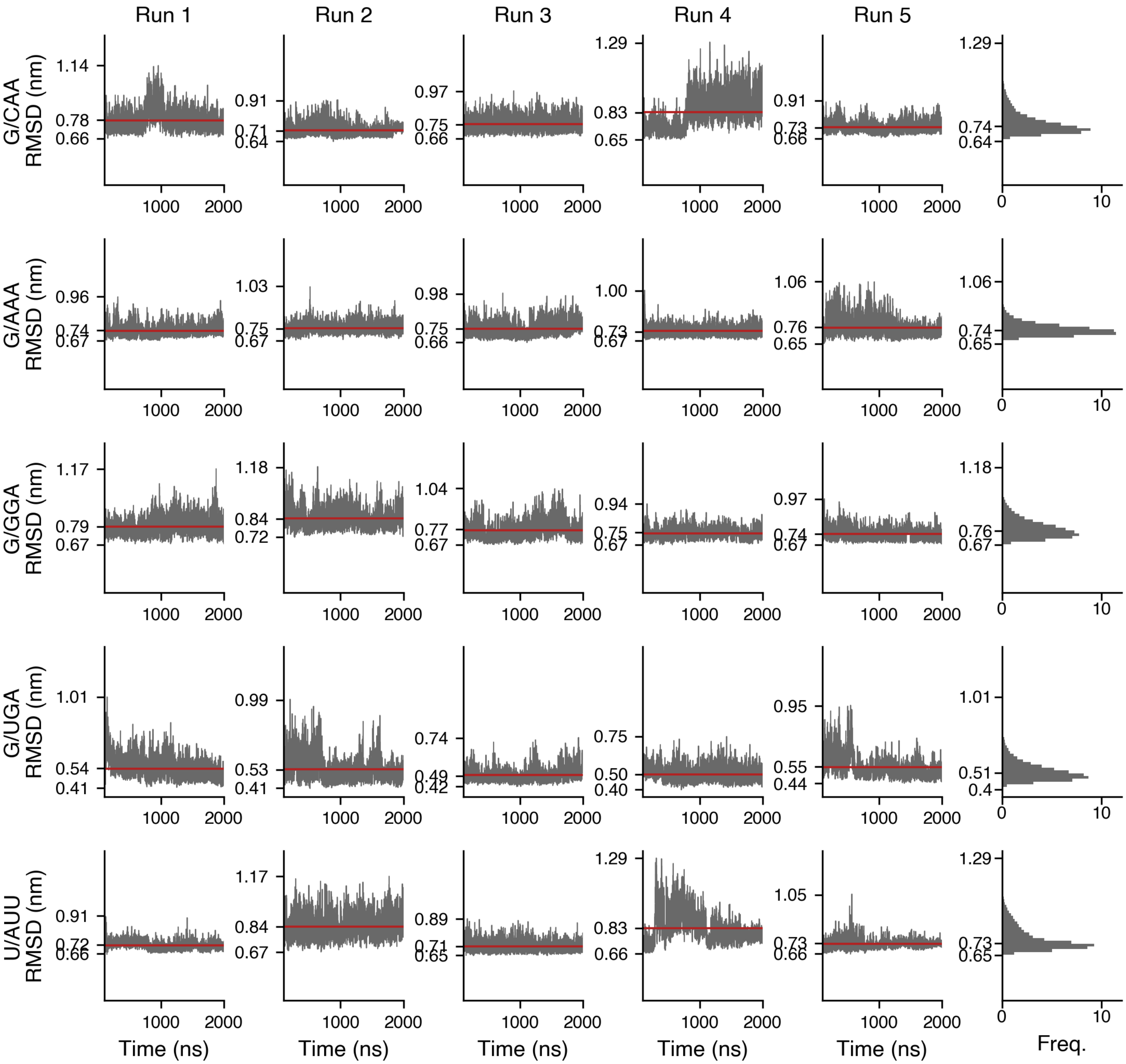


Figure S37: Root mean squared displacement (RMSD) of the first strand vs the initial model used to start the MD simulations for simulations of a dimer of the minimal alphaKL motif. The Y-axis ticks correspond to the minimum, mean and maximum RMSD value in each simulation trajectory. Note that RMSD was only calculated for one strand to make these trajectories directly comparable with the monomeric state (Fig S36)

**Figure S38. RMSD of interface helix vs initial model**

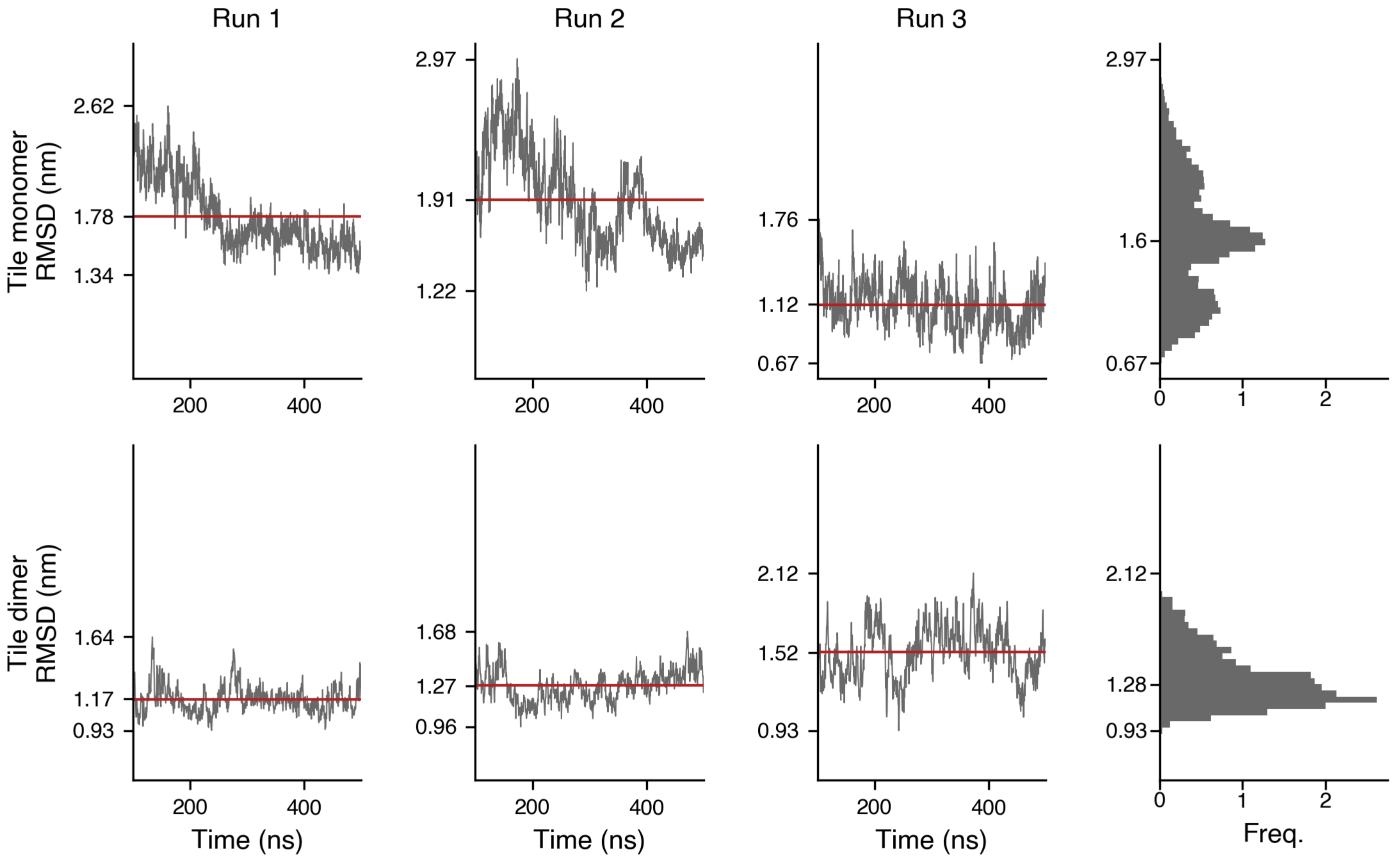


Figure S38: Root mean squared displacement (RMSD) of the interface helix vs the initial model used to start the MD simulations for simulations of a 3-helix tile dimer joined by alphaKLs. Top: Monomeric tile. Bottom: The first tile in the tile dimer. The Y-axis ticks correspond to the minimum, mean and maximum RMSD value in each simulation trajectory.

**Figure S39. RMSD of non-interface helies vs initial model**

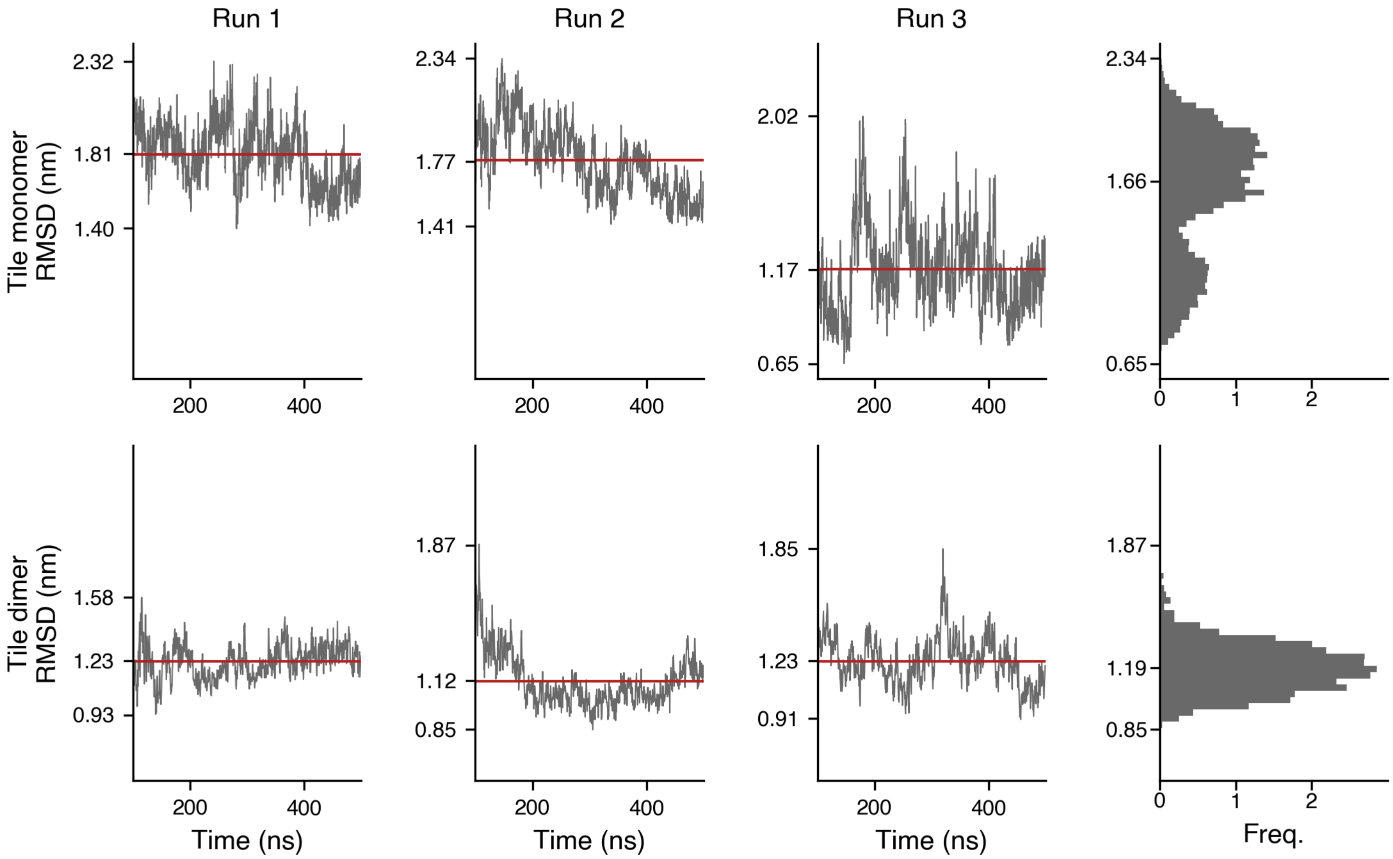


Figure S39: Root mean squared displacement (RMSD) of the non-interface helices vs the initial model used to start the MD simulations for simulations of a 3-helix tile dimer joined by alphaKLs. Top: Monomeric tile. Bottom: The first tile in the tile dimer. The Y-axis ticks correspond to the minimum, mean and maximum RMSD value in each simulation trajectory.

**Figure S40. RMSF of monomer vs dimerized form of tile**

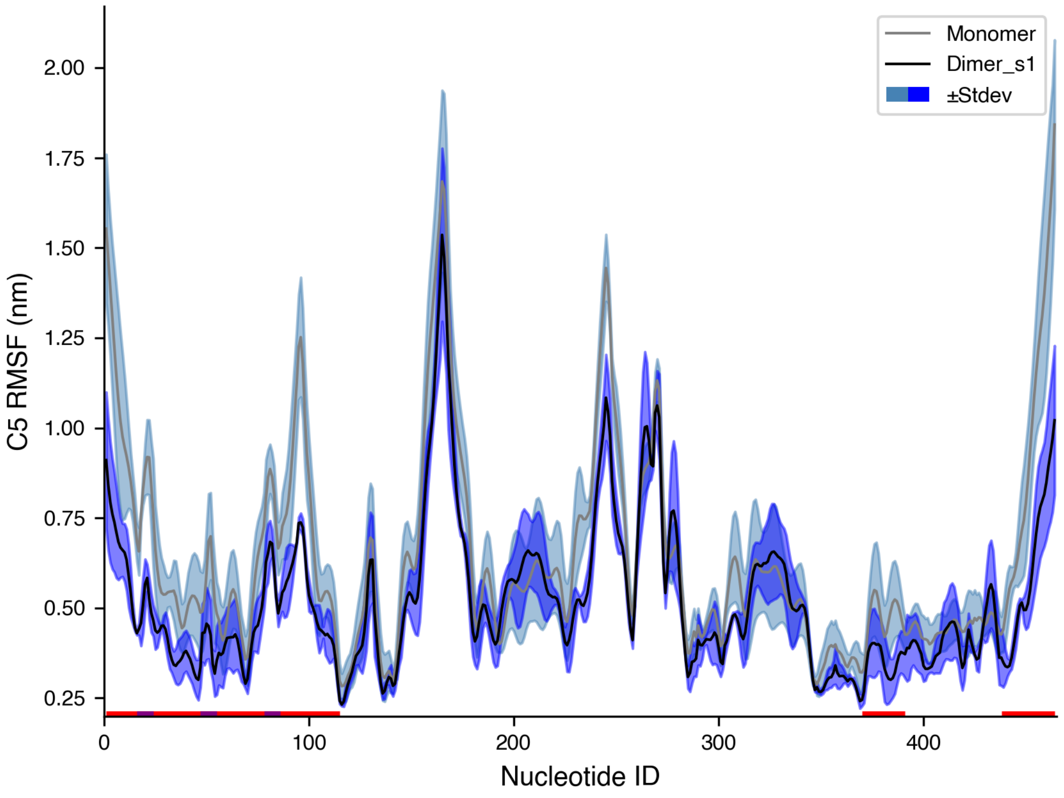


Figure S40: Comparison of the RMSF of C5' atoms between the monomeric tile and the first tile in the tile dimer. The monomer almost always has higher flexibility, particularly at the interface helix (red highlight) and the alphaKLs (purple highlight).

**Figure S41. RMSF difference between monomer and dimerized tile**

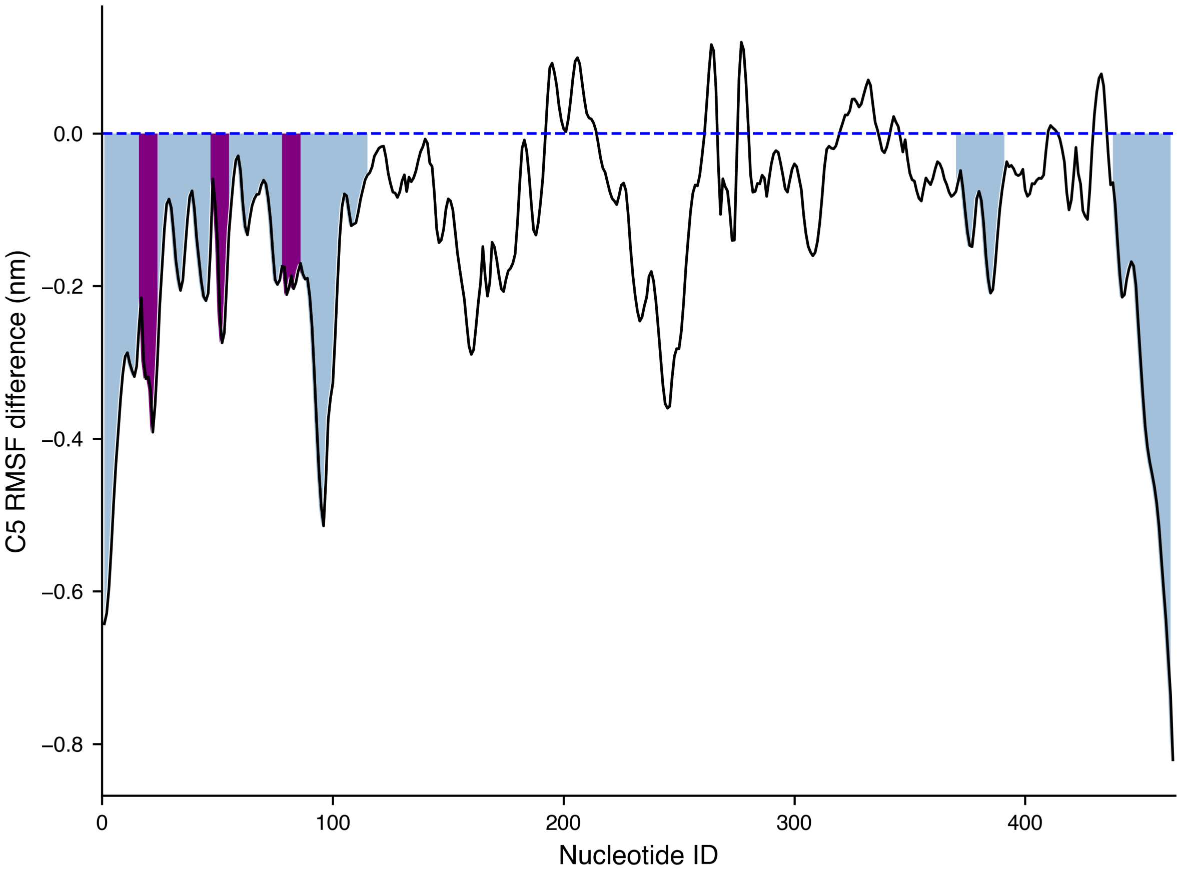


Figure S41: The difference between the monomer and dimerized tile RMSF data shown in Fig. S40. Blue-highlighted regions are the interface helix, while purple-highlighted regions are the alphaKLs.

#### Figure S42. Failed Lattice Design

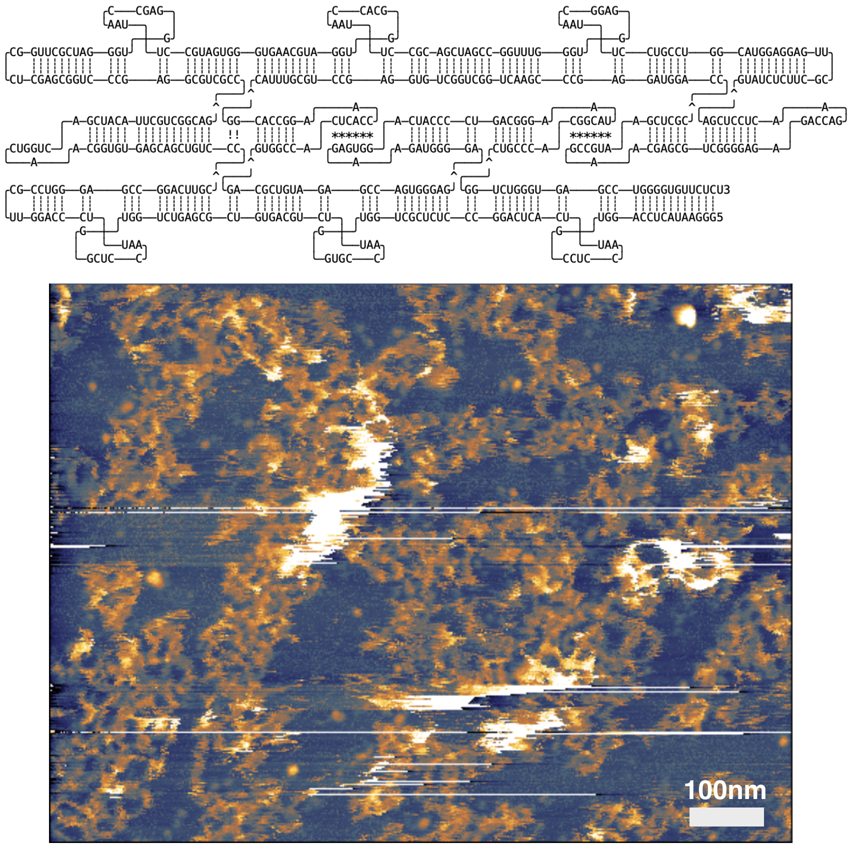


Figure S42: AFM image of failed fibril design with 180KL end-to-end connections linking fibers. After cotranscriptional assembly in solution, RNA products were deposited on 25°C mica.

#### Figure S43. Cotranscriptional nanosquare lattice with G/AAA alphaKL, nucleating on mica

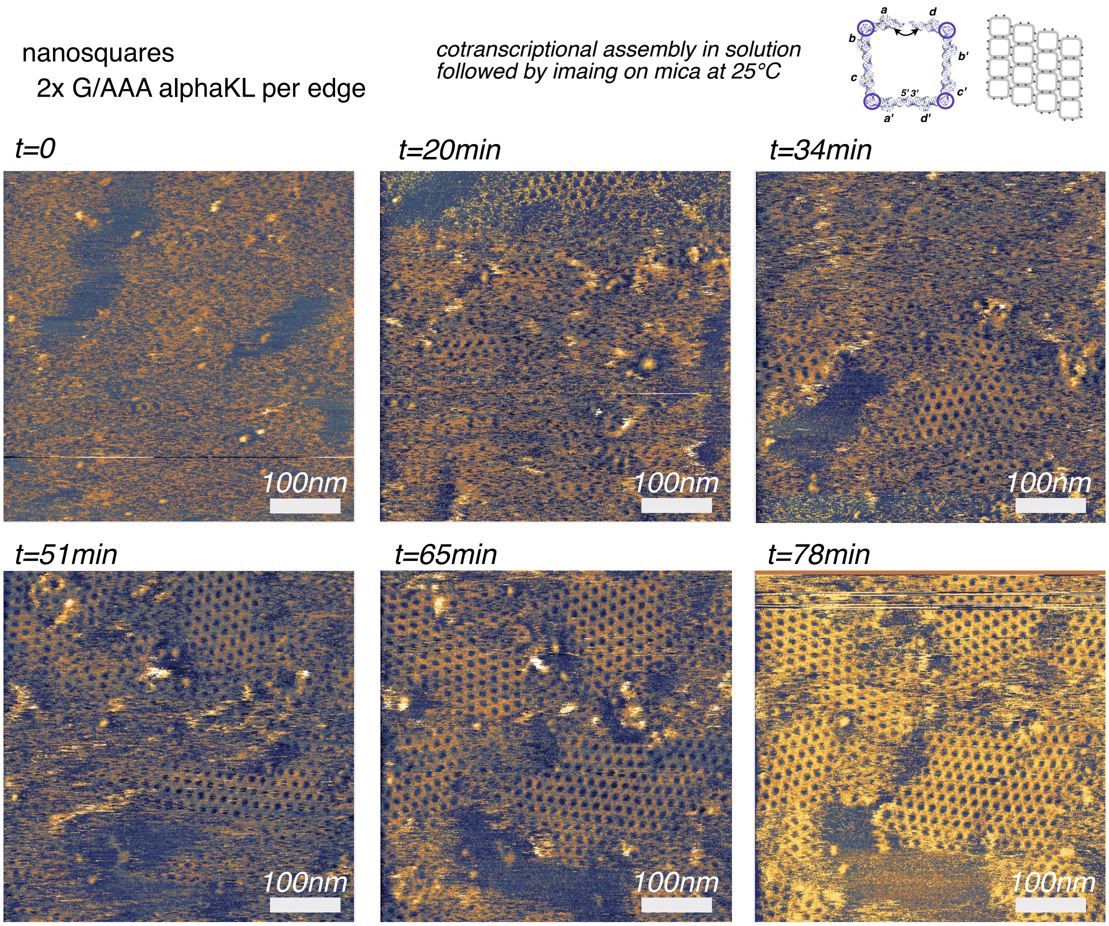


Figure S43: Additional AFM images of nanosquare assemblies. After cotranscriptional assembly in solution, RNA products were deposited on 25°C mica, and imaged continuously in solution. Over the imaging session, the small volume of imaging buffer became more concentrated due to evaporation.

**Figure S44. RNA fiber curvature measurement**

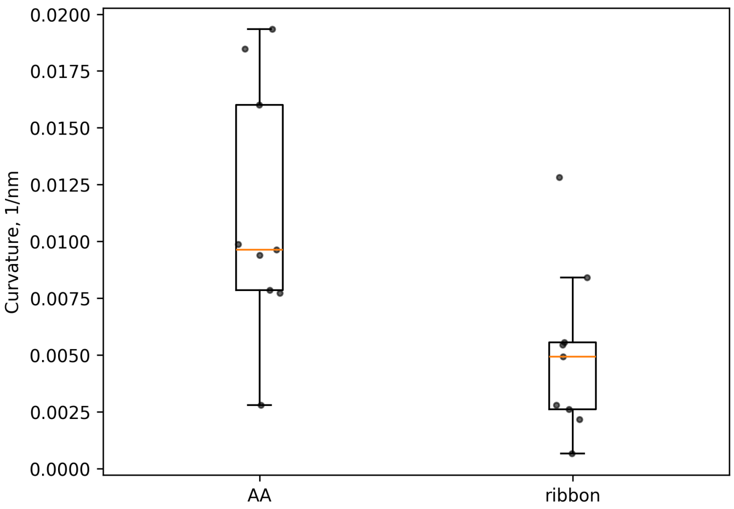


Figure S44: Measurements of G/AAA 3H3x-Fiber and 2H4x-Fiber curvature after cotranscriptional assembly in solution.

#### Figure S45. KL energies

```
Outputting 4bp 1:1 duplex data:

--- Homodimers ---
GGCC <-> GGCC : -5.90 [((((&))))]
GCGC <-> GCGC : -5.10 [((((&))))]
CCGG <-> CCGG : -4.90 [((((&))))]
CGCG <-> CGCG : -4.10 [((((&))))]
AGCU <-> AGCU : -2.50 [((((&))))]
UGCA <-> UGCA : -2.50 [((((&))))]
UCGA <-> UCGA : -2.10 [((((&))))]
GAUC <-> GAUC : -1.80 [((((&))))]
ACGU <-> ACGU : -1.70 [((((&))))]
GUAC <-> GUAC : -1.60 [((((&))))]
CUAG <-> CUAG : -1.40 [((((&))))]
CAUG <-> CAUG : -1.20 [((((&))))]
AAUU <-> AAUU : 0 [((((&))))]
AUAU <-> AUAU : 0 [((((&))))]
UAUA <-> UAUA : 0 [((((&))))]
UUAA <-> UUAA : 0 [.((.&.)).]

--- Heterodimers ---
GGGC <-> GCCC : -5.90 [((((&))))]
GGGG <-> CCCC : -5.80 [((((&))))]
GGCG <-> CGCC : -5.00 [((((&))))]
GCGG <-> CCGC : -5.00 [((((&))))]
GCCG <-> CGGC : -5.00 [((((&))))]
CGGG <-> CCCG : -4.90 [((((&))))]
UCCC <-> GGGA : -4.40 [((((&))))]
AGGC <-> GCCU : -4.20 [((((&))))]
AGCC <-> GGCU : -4.20 [((((&))))]
ACCC <-> GGGU : -4.20 [((((&))))]
UGGC <-> GCCA : -4.20 [((((&))))]
UGCC <-> GGCA : -4.20 [((((&))))]
AGGG <-> CCCU : -4.10 [((((&))))]
UGGG <-> CCCA : -4.10 [((((&))))]
GAGC <-> GCUC : -3.80 [((((&))))]
GACC <-> GGUC : -3.80 [((((&))))]
GUCC <-> GGAC : -3.80 [((((&))))]
GAGG <-> CCUC : -3.70 [((((&))))]
GGAG <-> CUCC : -3.70 [((((&))))]
UCGC <-> GCGA : -3.60 [((((&))))]
GUGC <-> GCAC : -3.60 [((((&))))]
UCGG <-> CCGA : -3.50 [((((&))))]
UCCG <-> CGGA : -3.50 [((((&))))]
GUGG <-> CCAC : -3.50 [((((&))))]
GGUG <-> CACC : -3.50 [((((&))))]
GCAG <-> CUGC : -3.50 [((((&))))]
GCUG <-> CAGC : -3.50 [((((&))))]
ACGC <-> GCGU : -3.40 [((((&))))]
CAGG <-> CCUG : -3.40 [((((&))))]
CUGG <-> CCAG : -3.40 [((((&))))]
AGCG <-> CGCU : -3.30 [((((&))))]
ACGG <-> CCGU : -3.30 [((((&))))]
ACCG <-> CGGU : -3.30 [((((&))))]
UGCG <-> CGCA : -3.30 [((((&))))]
GACG <-> CGUC : -2.90 [((((&))))]
GUCG <-> CGAC : -2.90 [((((&))))]
CUCG <-> CGAG : -2.80 [((((&))))]
AGGA <-> UCCU : -2.70 [((((&))))]
UGGA <-> UCCA : -2.70 [((((&))))]
CACG <-> CGUG : -2.60 [((((&))))]
AGGU <-> ACCU : -2.50 [((((&))))]
AGCA <-> UGCU : -2.50 [((((&))))]
ACCA <-> UGGU : -2.50 [((((&))))]
UCUC <-> GAGA : -2.30 [((((&))))]
AUCC <-> GGAU : -2.20 [((((&))))]
UAGC <-> GCUA : -2.20 [((((&))))]
UACC <-> GGUA : -2.20 [((((&))))]
AGAC <-> GUCU : -2.10 [((((&))))]
AGUC <-> GACU : -2.10 [((((&))))]
ACUC <-> GAGU : -2.10 [((((&))))]
UAGG <-> CCUA : -2.10 [((((&))))]
UUCC <-> GGAA : -2.10 [.(((&))).]
UGAC <-> GUCA : -2.10 [((((&))))]
UGUC <-> GACA : -2.10 [((((&))))]
UCAC <-> GUGA : -2.10 [((((&))))]
AUGC <-> GCAU : -2.00 [((((&))))]
AGAG <-> CUCU : -2.00 [((((&))))]
UGAG <-> CUCA : -2.00 [((((&))))]
UCAG <-> CUGA : -2.00 [((((&))))]
UCUG <-> CAGA : -2.00 [((((&))))]
AUGG <-> CCAU : -1.90 [((((&))))]
ACAC <-> GUGU : -1.90 [((((&))))]
ACGA <-> UCGU : -1.90 [((((&))))]
UUGC <-> GCAA : -1.90 [.(((&))).]
AAGC <-> GCUU : -1.80 [((((&))))]
AACC <-> GGUU : -1.80 [((((&))))]
AGUG <-> CACU : -1.80 [((((&))))]
ACAG <-> CUGU : -1.80 [((((&))))]
ACUG <-> CAGU : -1.80 [((((&))))]
UUGG <-> CCAA : -1.80 [.(((&))).]
UGUG <-> CACA : -1.80 [((((&))))]
AAGG <-> CCUU : -1.70 [((((&))))]
GAUG <-> CAUC : -1.50 [((((&))))]
GUAG <-> CUAC : -1.50 [((((&))))]
GAAC <-> GUUC : -1.40 [((((&))))]
AUCG <-> CGAU : -1.30 [((((&))))]
UACG <-> CGUA : -1.30 [((((&))))]
GAAG <-> CUUC : -1.30 [((((&))))]
UUCG <-> CGAA : -1.20 [.(((&))).]
GUUG <-> CAAC : -1.10 [((((&))))]
CAAG <-> CUUG : -1.00 [((((&))))]
AACG <-> CGUU : -0.90 [((((&))))]
UAGA <-> UCUA : -0.70 [((((&))))]
AUGA <-> UCAU : -0.50 [((((&))))]
AUCA <-> UGAU : -0.50 [((((&))))]
AUCU <-> AGAU : -0.50 [((((&))))]
AGUA <-> UACU : -0.50 [((((&))))]
ACUA <-> UAGU : -0.50 [((((&))))]
UACA <-> UGUA : -0.50 [((((&))))]
AGAA <-> UUCU : -0.40 [(((.&.)))]
UUGA <-> UCAA : -0.40 [.(((&))).]
UUCA <-> UGAA : -0.40 [.(((&))).]
AAGA <-> UCUU : -0.30 [((((&))))]
AUGU <-> ACAU : -0.30 [((((&))))]
ACAA <-> UUGU : -0.20 [(((.&.)))]
UAUC <-> GAUA : -0.20 [((((&))))]
AAGU <-> ACUU : -0.10 [((((&))))]
AACA <-> UGUU : -0.10 [((((&))))]
AACU <-> AGUU : -0.10 [((((&))))]
AAAA <-> UUUU : 0 [(((.&.)))]
AAAU <-> AUUU : 0 [((((&))))]
AAAG <-> CUUU : 0 [((((&))))]
AAAC <-> GUUU : 0 [((((&))))]
AAUA <-> UAUU : 0 [((((&))))]
AAUG <-> CAUU : 0 [((((&))))]
AAUC <-> GAUU : 0 [((((&))))]
AUAA <-> UUAU : 0 [(((.&.)))]
AUAG <-> CUAU : 0 [((((&))))]
AUAC <-> GUAU : 0 [((((&))))]
AUUA <-> UAAU : 0 [((((&))))]
AUUG <-> CAAU : 0 [((((&))))]
AUUC <-> GAAU : 0 [((((&))))]
UAAA <-> UUUA : 0 [(((.&.)))]
UAAG <-> CUUA : 0 [((((&))))]
UAAC <-> GUUA : 0 [((((&))))]
UAUG <-> CAUA : 0 [((((&))))]
UUAG <-> CUAA : 0 [.(((&))).]
UUAC <-> GUAA : 0 [.(((&))).]
UUUG <-> CAAA : 0 [.(((&))).]
UUUC <-> GAAA : 0 [.(((&))).]
```

Figure S45: Analysis of all possible 4 nt kissing loop sequences, sorted by predicted energy.

#### Figure S46. KL energies with angles

```
Computed with flanking dangling nts :

--- Homodimers ---
G-GGCC-A <-> G-GGCC-A : -8.70 [.((((.&.)))).]
G-CGCG-A <-> G-CGCG-A : -8.50 [((((.&)))).]
G-GCGC-A <-> G-GCGC-A : -7.90 [.((((.&.)))).]
G-CCGG-A <-> G-CCGG-A : -7.50 [.((((.&.)))).]
G-AGCU-A <-> G-AGCU-A : -4.70 [.((((.&.)))).]
G-GAUC-A <-> G-GAUC-A : -4.60 [.((((.&.)))).]
G-GUAC-A <-> G-GUAC-A : -4.40 [.((((.&.)))).]
G-UGCA-A <-> G-UGCA-A : -4.10 [.((((.&.)))).]
G-CUAG-A <-> G-CUAG-A : -4.00 [.((((.&.)))).]
G-ACGU-A <-> G-ACGU-A : -3.90 [.((((.&.)))).]
G-CAUG-A <-> G-CAUG-A : -3.80 [.((((.&.)))).]
G-UCGA-A <-> G-UCGA-A : -3.70 [.((((.&.)))).]
G-AUAU-A <-> G-AUAU-A : -0.60 [.((((.&.)))).]
G-UAUA-A <-> G-UAUA-A : -0.40 [((((.&)))).]
G-AAUU-A <-> G-AAUU-A : 0 [.((((.&.)))).]
G-UUAA-A <-> G-UUAA-A : 0 [.((((.&.)))).]

--- Heterodimers ---
G-GGGG-A <-> G-CCCC-A : -8.90 [((((.&.)))).]
G-GGGC-A <-> G-GCCC-A : -8.70 [.((((.&.)))).]
G-GGCG-A <-> G-CGCC-A : -7.70 [.((((.&.)))).]
G-GCGG-A <-> G-CCGC-A : -7.70 [.((((.&.)))).]
G-GCCG-A <-> G-CGGC-A : -7.70 [.((((.&.)))).]
G-CGGG-A <-> G-CCCG-A : -7.50 [.((((.&.)))).]
G-AGGC-A <-> G-GCCU-A : -6.70 [.((((.&.)))).]
G-AGCC-A <-> G-GGCU-A : -6.70 [.((((.&.)))).]
G-ACCC-A <-> G-GGGU-A : -6.70 [.((((.&.)))).]
G-UGCG-A <-> G-CGCA-A : -6.60 [((((.&)))).]
G-UCCC-A <-> G-GGGA-A : -6.60 [.((((.&.)))).]
G-GAGC-A <-> G-GCUC-A : -6.60 [.((((.&.)))).]
G-GACC-A <-> G-GGUC-A : -6.60 [.((((.&.)))).]
G-GUCC-A <-> G-GGAC-A : -6.60 [.((((.&.)))).]
G-AGGG-A <-> G-CCCU-A : -6.50 [.((((.&.)))).]
G-UGGC-A <-> G-GCCA-A : -6.40 [.((((.&.)))).]
G-UGCC-A <-> G-GGCA-A : -6.40 [.((((.&.)))).]
G-GAGG-A <-> G-CCUC-A : -6.40 [.((((.&.)))).]
G-GUGC-A <-> G-GCAC-A : -6.40 [.((((.&.)))).]
G-GGAG-A <-> G-CUCC-A : -6.40 [.((((.&.)))).]
G-UGGG-A <-> G-CCCA-A : -6.20 [.((((.&.)))).]
G-GUGG-A <-> G-CCAC-A : -6.20 [.((((.&.)))).]
G-GGUG-A <-> G-CACC-A : -6.20 [.((((.&.)))).]
G-GCAG-A <-> G-CUGC-A : -6.20 [.((((.&.)))).]
G-GCUG-A <-> G-CAGC-A : -6.20 [.((((.&.)))).]
G-CAGG-A <-> G-CCUG-A : -6.00 [.((((.&.)))).]
G-CUGG-A <-> G-CCAG-A : -6.00 [.((((.&.)))).]
G-ACGC-A <-> G-GCGU-A : -5.90 [.((((.&.)))).]
G-UCGC-A <-> G-GCGA-A : -5.80 [.((((.&.)))).]
G-AGCG-A <-> G-CGCU-A : -5.70 [.((((.&.)))).]
G-ACGG-A <-> G-CCGU-A : -5.70 [.((((.&.)))).]
G-ACCG-A <-> G-CGGU-A : -5.70 [.((((.&.)))).]
G-UCGG-A <-> G-CCGA-A : -5.60 [.((((.&.)))).]
G-UCCG-A <-> G-CGGA-A : -5.60 [.((((.&.)))).]
G-GACG-A <-> G-CGUC-A : -5.60 [.((((.&.)))).]
G-GUCG-A <-> G-CGAC-A : -5.60 [.((((.&.)))).]
G-CUCG-A <-> G-CGAG-A : -5.40 [.((((.&.)))).]
G-CACG-A <-> G-CGUG-A : -5.20 [.((((.&.)))).]
G-AUCC-A <-> G-GGAU-A : -4.70 [.((((.&.)))).]
G-AGGU-A <-> G-ACCU-A : -4.70 [.((((.&.)))).]
G-AGAC-A <-> G-GUCU-A : -4.60 [.((((.&.)))).]
G-AGUC-A <-> G-GACU-A : -4.60 [.((((.&.)))).]
G-AGGA-A <-> G-UCCU-A : -4.60 [.((((.&.)))).]
G-ACUC-A <-> G-GAGU-A : -4.60 [.((((.&.)))).]
G-UGUG-A <-> G-CACA-A : -4.60 [((((.&)))).]
G-AUGC-A <-> G-GCAU-A : -4.50 [.((((.&.)))).]
G-UCUC-A <-> G-GAGA-A : -4.50 [.((((.&.)))).]
G-AGAG-A <-> G-CUCU-A : -4.40 [.((((.&.)))).]
G-AGCA-A <-> G-UGCU-A : -4.40 [.((((.&.)))).]
G-ACAC-A <-> G-GUGU-A : -4.40 [.((((.&.)))).]
G-ACCA-A <-> G-UGGU-A : -4.40 [.((((.&.)))).]
G-UAGC-A <-> G-GCUA-A : -4.40 [.((((.&.)))).]
G-UACC-A <-> G-GGUA-A : -4.40 [.((((.&.)))).]
G-AAGC-A <-> G-GCUU-A : -4.30 [.((((.&.)))).]
G-AACC-A <-> G-GGUU-A : -4.30 [.((((.&.)))).]
G-AUGG-A <-> G-CCAU-A : -4.30 [.((((.&.)))).]
G-UGAC-A <-> G-GUCA-A : -4.30 [.((((.&.)))).]
G-UGUC-A <-> G-GACA-A : -4.30 [.((((.&.)))).]
G-UGGA-A <-> G-UCCA-A : -4.30 [.((((.&.)))).]
G-UCAC-A <-> G-GUGA-A : -4.30 [.((((.&.)))).]
G-AGUG-A <-> G-CACU-A : -4.20 [.((((.&.)))).]
G-ACAG-A <-> G-CUGU-A : -4.20 [.((((.&.)))).]
G-ACUG-A <-> G-CAGU-A : -4.20 [.((((.&.)))).]
G-UAGG-A <-> G-CCUA-A : -4.20 [.((((.&.)))).]
G-UUCC-A <-> G-GGAA-A : -4.20 [.((((.&.)))).]
G-GAAC-A <-> G-GUUC-A : -4.20 [.((((.&.)))).]
G-GAUG-A <-> G-CAUC-A : -4.20 [.((((.&.)))).]
G-GUAG-A <-> G-CUAC-A : -4.20 [.((((.&.)))).]
G-AAGG-A <-> G-CCUU-A : -4.10 [.((((.&.)))).]
G-UGAG-A <-> G-CUCA-A : -4.10 [.((((.&.)))).]
G-UCAG-A <-> G-CUGA-A : -4.10 [.((((.&.)))).]
G-UCUG-A <-> G-CAGA-A : -4.10 [.((((.&.)))).]
G-UUGC-A <-> G-GCAA-A : -4.00 [.((((.&.)))).]
G-GAAG-A <-> G-CUUC-A : -4.00 [.((((.&.)))).]
G-ACGA-A <-> G-UCGU-A : -3.90 [(((.&))).]
G-UUGG-A <-> G-CCAA-A : -3.80 [.((((.&.)))).]
G-GUUG-A <-> G-CAAC-A : -3.80 [.((((.&.)))).]
G-AUCG-A <-> G-CGAU-A : -3.70 [.((((.&.)))).]
G-CAAG-A <-> G-CUUG-A : -3.60 [.((((.&.)))).]
G-UACG-A <-> G-CGUA-A : -3.40 [.((((.&.)))).]
G-AACG-A <-> G-CGUU-A : -3.30 [.((((.&.)))).]
G-UUCG-A <-> G-CGAA-A : -3.20 [.((((.&.)))).]
G-AUCU-A <-> G-AGAU-A : -2.70 [.((((.&.)))).]
G-AUAC-A <-> G-GUAU-A : -2.50 [.((((.&.)))).]
G-AUGU-A <-> G-ACAU-A : -2.50 [.((((.&.)))).]
G-AUGA-A <-> G-UCAU-A : -2.40 [.((((.&.)))).]
G-AUCA-A <-> G-UGAU-A : -2.40 [.((((.&.)))).]
G-AGUA-A <-> G-UACU-A : -2.40 [.((((.&.)))).]
G-ACUA-A <-> G-UAGU-A : -2.40 [.((((.&.)))).]
G-UAUC-A <-> G-GAUA-A : -2.40 [.((((.&.)))).]
G-AAUC-A <-> G-GAUU-A : -2.30 [.((((.&.)))).]
G-AAGU-A <-> G-ACUU-A : -2.30 [.((((.&.)))).]
G-AACU-A <-> G-AGUU-A : -2.30 [.((((.&.)))).]
G-AUAG-A <-> G-CUAU-A : -2.30 [.((((.&.)))).]
G-AUUC-A <-> G-GAAU-A : -2.30 [.((((.&.)))).]
G-UAGA-A <-> G-UCUA-A : -2.30 [.((((.&.)))).]
G-AAGA-A <-> G-UCUU-A : -2.20 [.((((.&.)))).]
G-AGAA-A <-> G-UUCU-A : -2.20 [.((((.&.)))).]
G-UACA-A <-> G-UGUA-A : -2.10 [.((((.&.)))).]
G-AACA-A <-> G-UGUU-A : -2.00 [.((((.&.)))).]
G-ACAA-A <-> G-UUGU-A : -2.00 [.((((.&.)))).]
G-UAAC-A <-> G-GUUA-A : -2.00 [.((((.&.)))).]
G-UAUG-A <-> G-CAUA-A : -2.00 [.((((.&.)))).]
G-UUAC-A <-> G-GUAA-A : -2.00 [.((((.&.)))).]
G-AAAC-A <-> G-GUUU-A : -1.90 [.((((.&.)))).]
G-AAUG-A <-> G-CAUU-A : -1.90 [.((((.&.)))).]
G-AUUG-A <-> G-CAAU-A : -1.90 [.((((.&.)))).]
G-UUGA-A <-> G-UCAA-A : -1.90 [.((((.&.)))).]
G-UUCA-A <-> G-UGAA-A : -1.90 [.((((.&.)))).]
G-UAAG-A <-> G-CUUA-A : -1.80 [.((((.&.)))).]
G-UUAG-A <-> G-CUAA-A : -1.80 [.((((.&.)))).]
G-UUUC-A <-> G-GAAA-A : -1.80 [.((((.&.)))).]
G-AAAG-A <-> G-CUUU-A : -1.70 [.((((.&.)))).]
G-UUUG-A <-> G-CAAA-A : -1.40 [.((((.&.)))).]
G-AAUA-A <-> G-UAUU-A : -0.10 [.((((.&.)))).]
G-AUAA-A <-> G-UUAU-A : -0.10 [.((((.&.)))).]
G-AUUA-A <-> G-UAAU-A : -0.10 [.((((.&.)))).]
G-AAAA-A <-> G-UUUU-A : 0 [((((.&.)))).]
G-AAAU-A <-> G-AUUU-A : 0 [.((((.&.)))).]
G-UAAA-A <-> G-UUUA-A : 0 [.((((.&.)))).]
```

Figure S46: Analysis of all possible 4 nt kissing loop sequences including dangle energies, sorted by predicted energy.

#### Figure S47. Other Potential KL Connector Contexts for AlphaKL

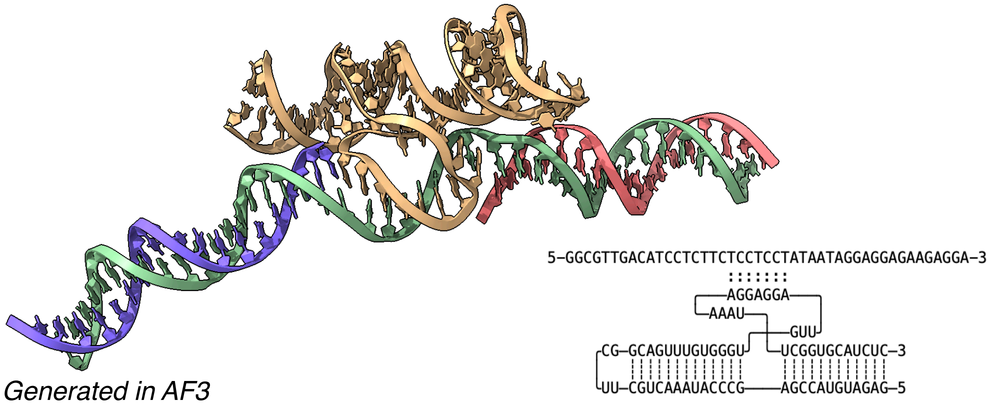


Proposed elongated-alphaKL:DNA Complex

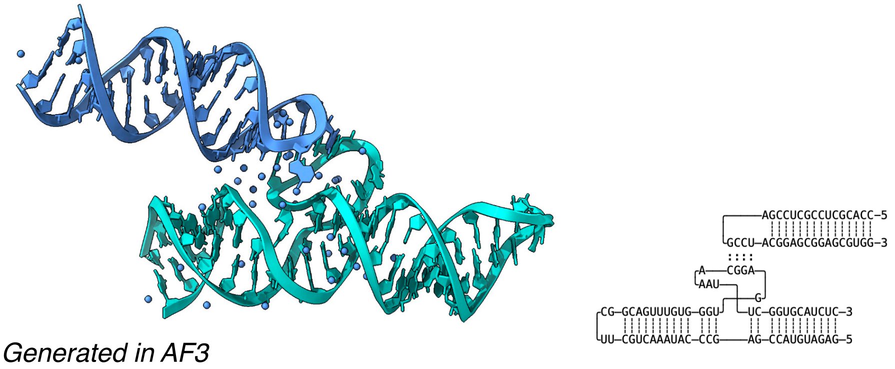


Proposed alphaKL:stem loop complex

Figure S47: AlphaKL connectors can potentially be used in other contexts than the describes loop-loop interaction. As the programmable interface is simply a stretch of 4bps, we can imagine increasing the length of the interface to a longer one by expanding the major and minor groove triples further. In such a context a DNA strand could possibly be addressed sequence-specifically (top). Alternatively, the alphaKL might also be used to tether hairpin terminal loops to a larger RNA origami if the precise parallel alignment is not required.

#### Figure S48. Sequence Design

Design: Nanoring-3x-alphaKL (G/CAA)

DNA Template Sequence:

```
AGAGAACCCACCTAGCGAGCGTACGAAACCATCTCGCACACTCTCTACAGTTCCCTCTTCTG
CAGAGCTCAGTTACCGCTTCCGAGCGCCACCGGAAGGCTCTGCGGACAGTCGACGTGCGGTA
TTCATCGAGGGCACCCGTGAGGGATTCGGATGCGAAGCGATGGTTGCCTCCCTCGCTGCGCG
AACACAGCAAGGCCGTACCTAGGGCTCCCACGCGAACATGGGAATTGGAGCCACCCCAGGCA
CCTATCCACAGAATTGCCGGCACCCTCCAGTGCTAACCACGGAATTGGCTCCACCCCTTGCG
AACAAAGGGCTCCGTGATCAGTAGGCGGATTCCCTATAGTGAGTCGTATTAGAATGCCC
```

Forward Primer: GGGCATTCTAATACGACTCACTATA

Reverse Primer: AGAGAACCCACCTAGCGAG

RNA Sequence:

```
GGGAAUCCGCCUACUGAUCACGGAGCCCUUUGUUCGCAAGGGGUGGAGCCAAUUCCGUGGUU
AGCACUGGAGGGUGCCGGCAAUUCUGUGGAUAGGUGCCUGGGGUGGCUCCAAUUCCCAUGUU
CGCGUGGGAGCCCUAGGUACGGCCUUGCUGUGUUCGCGCAGCGAGGGAGGCAACCAUCGCUU
CGCAUCCGAAUCCCUCACGGGUGCCCUCGAUGAAUACCGCACGUCGACUGUCCGCAGAGCCU
UCCGGUGGCGCUCGGAAGCGGUAACUGAGCUCUGCAGAAGAGGGAACUGUAGAGAGUGUGCG
AGAUGGUUUCGUACGCUCGCUAGGUGGGUUCUCU
```

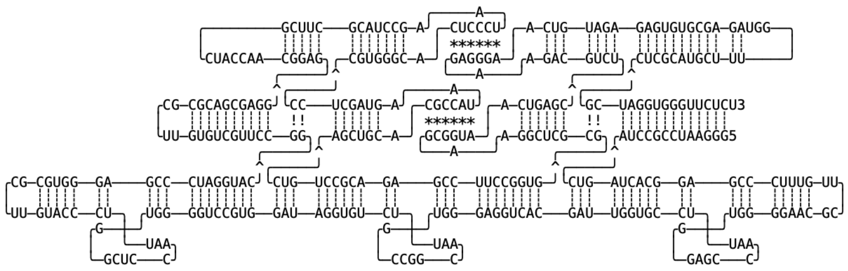

Figure S48: DNA template and primer sequences, RNA sequences and designed structure.

#### Figure S49. Sequence Design

##### Design: Nanoring-control (no motif)

DNA Template Sequence:

AGAGAACCCACCTAGCGAGCGTACGAAACCATCTCGCACACTCTCTACAGTTCCCTCTTCTG
CAGAGCTCAGTTACCGCTTCCGAGCGCCACCGGAAGGCTCTGCGGACAGTCGACGTGCGGTA
TTCATCGAGGGCACCCGTGAGGGATTCGGATGCGAAGCGATGGTTGCCTCCCTCGCTGCGCG
AACACAGCAAGGCCGTACCTAGGGCTCCCACGCGAACATGGGAACCCCAGGCACCTATCCAC
AGAACCCTCCAGTGCTAACCACGGAACCCCTTGCGAACAAAGGGCTCCGTGATCAGTAGGCG
GATTCCCTATAGTGAGTCGTATTAGAATGCCC

Forward Primer: GGGCATTCTAATACGACTCACTATA

Reverse Primer: AGAGAACCCACCTAGCGAG

RNA Sequence:

GGGAAUCCGCCUACUGAUCACGGAGCCCUUUGUUCGCAAGGGGUUCCGUGGUUAGCACUGGA
GGGUUCUGUGGAUAGGUGCCUGGGGUUCCCAUGUUCGCGUGGGAGCCCUAGGUACGGCCUUG
CUGUGUUCGCGCAGCGAGGGAGGCAACCAUCGCUUCGCAUCCGAAUCCCUCACGGGUGCCCU
CGAUGAAUACCGCACGUCGACUGUCCGCAGAGCCUUCCGGUGGCGCUCGGAAGCGGUAACUG
AGCUCUGCAGAAGAGGGAACUGUAGAGAGUGUGCGAGAUGGUUUCGUACGCUCGCUAGGUGG
GUUCUCU

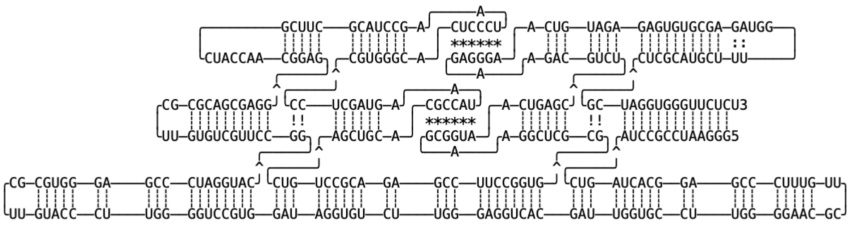

Figure S49: DNA template and primer sequences, RNA sequences and designed structure.

#### Figure S50. Sequence Design

##### Design: Strip-3x-alphaKL (G/CAA)

DNA Template Sequence:

```
AGAGAACACCCCAGGCTCACCCAGACCGACGGGTGCCGTATTCCCGTCAGGGGTAGTCCACT
CTTCTACCCCTCTCCCACTGGCTCTACAGCGTCCACCGGTGAGTGGTTCCGGTGCCGTAAAC
GCAGGCTCCACAGCCAGCCAGTTCGGGCTCCTACCTGGGCGAGCTTACGGCTTGCTCGCAGC
CCCACGGCGAACCATGGAGCTCATAGAGAAGCGAACTCCTCCATGCCAGGCAGGAATTGCCT
CCACCCAAACCGGCTAGCTGCGGAATTGGTGCCACCTACGTTCACCCACTACGGAATTGGCT
CCACCCTAGCGAACCGGAGCTCGCCAGGGCTCCGCAGCGGCTGCCGACAGCGTACCGTCGAC
AGGGGCAAGTCCGGCTCCCAGGCGAACCTGGGAATTGGAGCCACCAGACTCGCGACACTGCA
GAATTGGCACCACCAGCGAGAGGGCCTGAGTGAATTGGAGGCACCTGGAGTATTCCCTATAG
TGAGTCGTATTAGAATGCCC
```

Forward Primer: GGGCATTCTAATACGACTCACTATA

Reverse Primer: AGAGAACACCCCAGGCTC

RNA Sequence:

```
GGGAAUACUCCAGGUGCCUCCAAUUCACUCAGGCCCUCUCGCUGGUGGUGCCAAUUCUGCAG
UGUCGCGAGUCUGGUGGCUCCAAUUCCCAGGUUCGCCUGGGAGCCGGACUUGCCCCUGUCGA
CGGUACGCUGUCGGCAGCCGCUGCGGAGCCCUGGCGAGCUCCGGUUCGCUAGGGUGGAGCCA
AUUCCGUAGUGGGUGAACGUAGGUGGCACCAAUUCCGCAGCUAGCCGGUUUGGGUGGAGGCA
AUUCCUGCCUGGCAUGGAGGAGUUCGCUUCUCUAUGAGCUCCAUGGUUCGCCGUGGGGCUGC
GAGCAAGCCGUAAGCUCGCCCAGGUAGGAGCCCGAACUGGCUGGCUGUGGAGCCUGCGUUUA
CGGCACCGGAACCACUCACCGGUGGACGCUGUAGAGCCAGUGGGAGAGGGGUAGAAGAGUGG
ACUACCCCUGACGGGAAUACGGCACCCGUCGGUCUGGGUGAGCCUGGGGUGUUCUCU
```

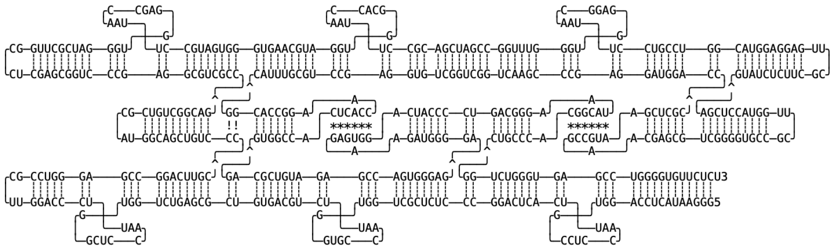

Figure S50: DNA template and primer sequences, RNA sequences and designed structure.

#### Figure S51. Sequence Design

Design: Strip-3x-alphaKL (G/AAA)

DNA Template Sequence:

AGAGAACACCCCAGGCTCACCCAGACCGACGGGTGCCGTATTCCCGTCAGGGGTAGTCCACT
CTTCTACCCCTCTCCCACTGGCTCTACAGCGTCCACCGGTGAGTGGTTCCGGTGCCGTAAAC
GCAGGCTCCACAGCCAGCCAGTTCGGGCTCCTACCTGGGCGAGCTTACGGCTTGCTCGCAGC
CCCACGGCGAACCATGGAGCTCATAGAGAAGCGAACTCCTCCATGCCAGGCAGGAATTTCCT
CCACCCAAACCGGCTAGCTGCGGAATTTGTGCCACCTACGTTCACCCACTACGGAATTTGCT
CCACCCTAGCGAACCGGAGCTCGCCAGGGCTCCGCAGCGGCTGCCGACAGCGTACCGTCGAC
AGGGGCAAGTCCGGCTCCCAGGCGAACCTGGGAATTTGAGCCACCAGACTCGCGACACTGCA
GAATTTGCACCACCAGCGAGAGGGCCTGAGTGAATTTGAGGCACCTGGAGTATTCCCTATAG
TGAGTCGTATTAGAATGCCC

Forward Primer: GGGCATTCTAATACGACTCACTATA

Reverse Primer: AGAGAACACCCCAGGCTC

RNA Sequence:
GGGAAUACUCCAGGUGCCUCAAAUUCACUCAGGCCCUCUCGCUGGUGGUGCAAAUUCUGCAG
UGUCGCGAGUCUGGUGGCUCAAAUUCCCAGGUUCGCCUGGGAGCCGGACUUGCCCCUGUCGA
CGGUACGCUGUCGGCAGCCGCUGCGGAGCCCUGGCGAGCUCCGGUUCGCUAGGGUGGAGCAA
AUUCCGUAGUGGGUGAACGUAGGUGGCACAAAUUCCGCAGCUAGCCGGUUUGGGUGGAGGAA
AUUCCUGCCUGGCAUGGAGGAGUUCGCUUCUCUAUGAGCUCCAUGGUUCGCCGUGGGGCUGC
GAGCAAGCCGUAAGCUCGCCCAGGUAGGAGCCCGAACUGGCUGGCUGUGGAGCCUGCGUUUA
CGGCACCGGAACCACUCACCGGUGGACGCUGUAGAGCCAGUGGGAGAGGGGUAGAAGAGUGG
ACUACCCCUGACGGGAAUACGGCACCCGUCGGUCUGGGUGAGCCUGGGGUGUUCUCU

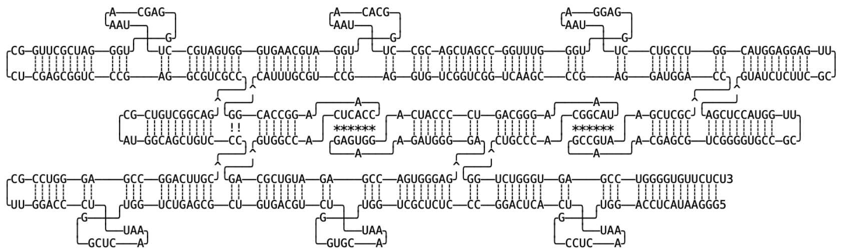

Figure S51: DNA template and primer sequences, RNA sequences and designed structure.

#### Figure S52. Sequence Design

Design: Strip-3x-alphaKL (G/AAA)

DNA Template Sequence:

```
AGAGAACACCCCAGGCTCACCCAGACCGACGGGTGCCGTATTCCCGTCAGGGGTAGTCCACT
CTTCTACCCCTCTCCCACTGGCTCTACAGCGTCCACCGGTGAGTGGTTCCGGTGCCGTAAAC
GCAGGCTCCACAGCCAGCCAGTTCGGGCTCCTACCTGGGCGAGCTTACGGCTTGCTCGCAGC
CCCACGGCGAACCATGGAGCTCATAGAGAAGCGAACTCCTCCATGCCAGGCAGGAATTTCCT
CCACCCAAACCGGCTAGCTGCGGAATTTGTGCCACCTACGTTCACCCACTACGGAATTTGCT
CCACCCTAGCGAACCGGAGCTCGCCAGGGCTCCGCAGCGGCTGCCGACAGCGTACCGTCGAC
AGGGGCAAGTCCGGCTCCCAGGCGAACCTGGGAATTTGAGCCACCAGACTCGCGACACTGCA
GAATTTGCACCACCAGCGAGAGGGCCTGAGTGAATTTGAGGCACCTGGAGTATTCCCTATAG
TGAGTCGTATTAGAATGCCC
```

Forward Primer: GGGCATTCTAATACGACTCACTATA

Reverse Primer: AGAGAACACCCCAGGCTC

RNA Sequence:

```
GGGAAUACUCCAGGUGCCUCAAAUUCACUCAGGCCCUCUCGCUGGUGGUGCAAAUUCUGCAG
UGUCGCGAGUCUGGUGGCUCAAAUUCCCAGGUUCGCCUGGGAGCCGGACUUGCCCCUGUCGA
CGGUACGCUGUCGGCAGCCGCUGCGGAGCCCUGGCGAGCUCCGGUUCGCUAGGGUGGAGCAA
AUUCCGUAGUGGGUGAACGUAGGUGGCACAAAUUCCGCAGCUAGCCGGUUUGGGUGGAGGAA
AUUCCUGCCUGGCAUGGAGGAGUUCGCUUCUCUAUGAGCUCCAUGGUUCGCCGUGGGGCUGC
GAGCAAGCCGUAAGCUCGCCCAGGUAGGAGCCCGAACUGGCUGGCUGUGGAGCCUGCGUUUA
CGGCACCGGAACCACUCACCGGUGGACGCUGUAGAGCCAGUGGGAGAGGGGUAGAAGAGUGG
ACUACCCCUGACGGGAAUACGGCACCCGUCGGUCUGGGUGAGCCUGGGGUGUUCUCU
```

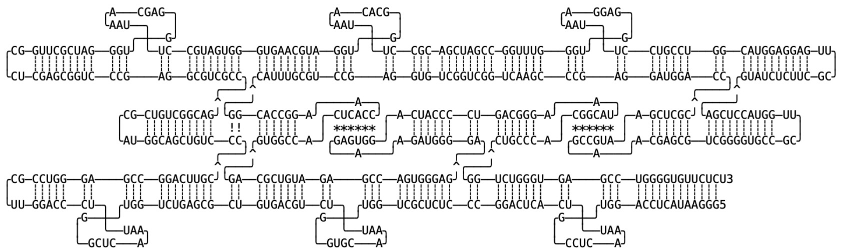

Figure S52: DNA template and primer sequences, RNA sequences and designed structure.

**Figure S53. Sequence Design**

#### Design: Strip-3x-alphaKL (G/GGA)

DNA Template Sequence:

```
AGAGAACACCCCAGGCTCACCCAGACCGACGCGTGCCGTATTCGCGTCAGGGGTAGTCCACT
CTTCTACCCCTCTCCCACTGGCTCTACAGCGTCCGTCACTGAGTGGTTGTGACGCCGTAAAC
GCAGGCTCCACAGCCAGCCAGTTCGGGCTCCTACCTGGGCGAGCTTACGGCTTGCTCGCAGC
CCCACGGCGAACCATGGAGCTCATAGAGAAGCGAACTCCTCCATGCCAGGCAGGAATCCCCT
CCACCCAAACCGGCTAGCTGCGGAATCCGTGCCACCTACGTTCACCCACTAAGGAATCCGCT
CCACCCTAGCGAACCGGAGCTCGCCAGGGCTCCTCAGCGGCTGCCGACAGCGTACCGTCGAC
AGGGGCAAGTGAGGCCCATAGGCGAACCTATGGATCCGAGCCACCCCACTCGCGACACTGCA
GAATCCGCACCACCAGCGAGAGGGCCTGAGTGAATCCGAGGCACCTGGAGTATTCCCTATAG
TGAGTCGTATTAGAATGCCC
```

Forward Primer: GGGCATTCTAATACGACTCACTATA

Reverse Primer: AGAGAACACCCCAGGCTC

RNA Sequence:

```
GGGAAUACUCCAGGUGCCUCGGAUUCACUCAGGCCCUCUCGCUGGUGGUGCGGAUUCUGCAG
UGUCGCGAGUGGGGUGGCUCGGAUCCAUAGGUUCGCCUAUGGGCCUCACUUGCCCCUGUCGA
CGGUACGCUGUCGGCAGCCGCUGAGGAGCCCUGGCGAGCUCCGGUUCGCUAGGGUGGAGCGG
AUUCCUUAGUGGGUGAACGUAGGUGGCACGGAUUCCGCAGCUAGCCGGUUUGGGUGGAGGGG
AUUCCUGCCUGGCAUGGAGGAGUUCGCUUCUCUAUGAGCUCCAUGGUUCGCCGUGGGGCUGC
GAGCAAGCCGUAAGCUCGCCCAGGUAGGAGCCCGAACUGGCUGGCUGUGGAGCCUGCGUUUA
CGGCGUCACAACCACUCAGUGACGGACGCUGUAGAGCCAGUGGGAGAGGGGUAGAAGAGUGG
ACUACCCCUGACGCGAAUACGGCACGCGUCGGUCUGGGUGAGCCUGGGGUGUUCUCU
```

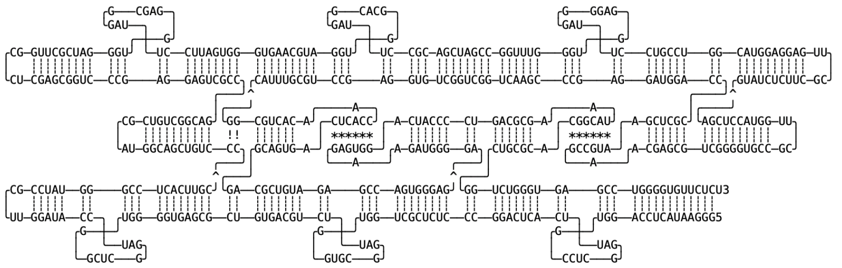

Figure S53: DNA template and primer sequences, RNA sequences and designed structure.

#### Figure S54. Sequence Design

##### Design: Strip-3x-alphaKL (G/UGA)

DNA Template Sequence:

```
AGAGAACACCCCAGGCTCACCCAGACCGACGCGTGCCGTATTCGCGTCAGGGGTAGTCCACT
CTTCTACCCCTCTCCCACTGGCTCTACAGCGTCCGTCACTGAGTGGTTGTGACGCCGTAAAC
GCAGGCTCCACAGCCAGCCAGTTCGGGCTCCTACCTGGGCGAGCTTACGGCTTGCTCGCAGC
CCCACGGCGAACCATGGAGCTCATAGAGAAGCGAACTCCTCCATGCCAGGCAGGAATCACCT
CCACCCAAACCGGCTAGCTGCGGAATCAGTGCCACCTACGTTCACCCACTAAGGAATCAGCT
CCACCCTAGCGAACCGGAGCTCGCCAGGGCTCCTCAGCGGCTGCCGACAGCGTACCGTCGAC
AGGGGCAAGTGAGGCCCATAGGCGAACCTATGGATCAGAGCCACCCCACTCGCGACACTGCA
GAATCAGCACCACCAGCGAGAGGGCCTGAGTGAATCAGAGGCACCTGGAGTATTCCCTATAG
TGAGTCGTATTAGAATGCCC
```

Forward Primer: GGGCATTCTAATACGACTCACTATA

Reverse Primer: AGAGAACACCCCAGGCTC

RNA Sequence:

```
GGGAAUACUCCAGGUGCCUCUGAUUCACUCAGGCCCUCUCGCUGGUGGUGCUGAUUCUGCAG
UGUCGCGAGUGGGGUGGCUCUGAUCCAUAGGUUCGCCUAUGGGCCUCACUUGCCCCUGUCGA
CGGUACGCUGUCGGCAGCCGCUGAGGAGCCCUGGCGAGCUCCGGUUCGCUAGGGUGGAGCUG
AUUCCUUAGUGGGUGAACGUAGGUGGCACUGAUUCCGCAGCUAGCCGGUUUGGGUGGAGGUG
AUUCCUGCCUGGCAUGGAGGAGUUCGCUUCUCUAUGAGCUCCAUGGUUCGCCGUGGGGCUGC
GAGCAAGCCGUAAGCUCGCCCAGGUAGGAGCCCGAACUGGCUGGCUGUGGAGCCUGCGUUUA
CGGCGUCACAACCACUCAGUGACGGACGCUGUAGAGCCAGUGGGAGAGGGGUAGAAGAGUGG
ACUACCCCUGACGCGAAUACGGCACGCGUCGGUCUGGGUGAGCCUGGGGUGUUCUCU
```

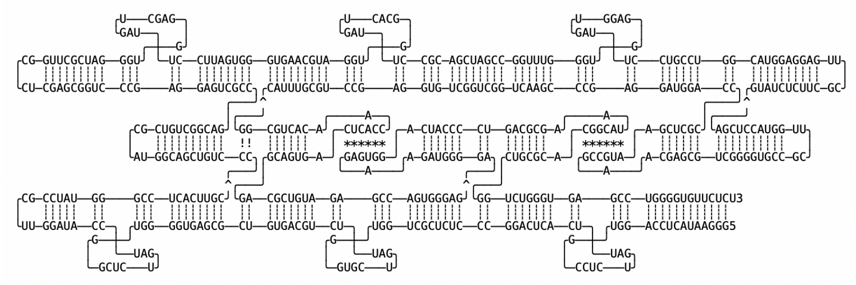

Figure S54: DNA template and primer sequences, RNA sequences and designed structure.

#### Figure S55. Sequence Design

##### Design: Strip-3x-alphaKL (U/AUU)

dsDNA Template Sequence:

AGAGAACACCCCAGGCTCACCCAGACCGACGGGTGCCGTATTCCCGTCAGGGGTAGTCCACT
CTTCTACCCCTCTCCCACTGGCTCTACAGCGTCCACCGGTGAGTGGTTCCGGTGCCGTAAAC
GCAGGCTCCACAGCCAGCCAGTTCGGGCTCCTACCTGGGCGAGCTTACGGCTTGCTCGCAGC
CCCACGGCGAACCATGGAGCTCATAGAGAAGCGAACTCCTCCATGCCAGGCAGGAAAATCCT
CAACCCAAACCGGCTAGCTGCGGAAAATGTGCAACCTACGTTCACCCACTACGGAAAATGCT
CAACCCTAGCGAACCGGAGCTCGCCAGGGCTCCGCAGCGGCTGCCGACAGCGTACCGTCGAC
AGGGGCAAGTCCGGCTCCCAGGCGAACCTGGGAAAATGAGCAACCAGACTCGCGACACTGCA
GAAAATGCACAACCAGCGAGAGGGCCTGAGTGAAAATGAGGAACCTGGAGTATTCCCTATAG
TGAGTCGTATTAGAATGCCC

Forward Primer: GGGCATTCTAATACGACTCACTATA

Reverse Primer: AGAGAACACCCCAGGCT

RNA Sequence:

GGGAAUACUCCAGGUUCCUCAUUUUCACUCAGGCCCUCUCGCUGGUUGUGCAUUUUCUGCAGUGUCGC
GAGUCUGGUUGCUCAUUUUCCCAGGUUCGCCUGGGAGCCGGACUUGCCCCUGUCGACGGUACGCUGUC
GGCAGCCGCUGCGGAGCCCUGGCGAGCUCCGGUUCGCUAGGGUUGAGCAUUUUCCGUAGUGGGUGAAC
GUAGGUUGCACAUUUUCCGCAGCUAGCCGGUUUGGGUUGAGGAUUUUCCUGCCUGGCAUGGAGGAGUU
CGCUUCUCUAUGAGCUCCAUGGUUCGCCGUGGGGCUGCGAGCAAGCCGUAAGCUCGCCCAGGUAGGAG
CCCGAACUGGCUGGCUGUGGAGCCUGCGUUUACGGCACCGGAACCACUCACCGGUGGACGCUGUAGAG
CCAGUGGGAGAGGGGUAGAAGAGUGGACUACCCCUGACGGGAAUACGGCACCCGUCGGUCUGGGUGAG
CCUGGGGUGUUCUCU

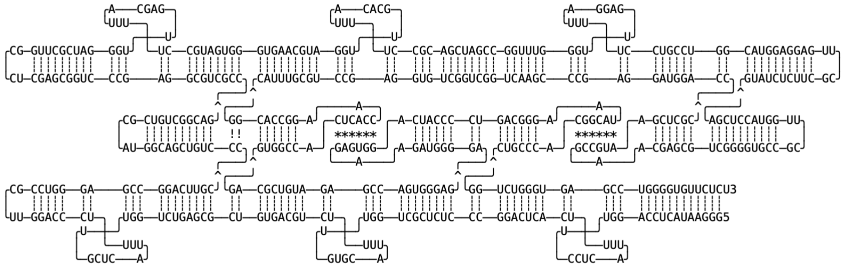

Figure S55: DNA template and primer sequences, RNA sequences and designed structure.

#### Figure S56. Sequence Design

##### Design: Strip-2x-alphaKL (G/AAA)

DNA Template Sequence:

AGAGAACACCCCAGGCTCACCCAGACCGACGGGTGCCGTATTCCCGTCAGGGGTAGTCCACT
CTTCTACCCCTCTCCCACTGGCTCTACAGCGTCCACCGGTGAGTGGTTCCGGTGCCGTAAAC
GCAGGCTCCACAGCCAGCCAGTTCGGGCTCCTACCTGGGCGAGCTTACGGCTTGCTCGCAGC
CCCACGGCGAACCATGGAGCTCATAGAGAAGCGAACTCCTCCATGCCAGGCAGGAATTTCCT
CCACCCAAACCGGCTAGCTGCGGAACCTACGTTCACCCACTACGGAATTTGCTCCACCCTAG
CGAACCGGAGCTCGCCAGGGCTCCGCAGCGGCTGCCGACAGCGTACCGTCGACAGGGGCAAG
TCCGGCTCCCAGGCGAACCTGGGAATTTGAGCCACCAGACTCGCGACACTGCAGAACCAGCG
AGAGGGCCTGAGTGAATTTGAGGCACCTGGAGTATTCCCTATAGTGAGTCGTATTAGAATGC
CC

Forward Primer: GGGCATTCTAATACGACTCACTATA

Reverse Primer: AGAGAACACCCCAGGCTCAC

RNA Sequence:
GGGAAUACUCCAGGUGCCUCAAAUUCACUCAGGCCCUCUCGCUGGUUCUGCAGUGUCGCGAG
UCUGGUGGCUCAAAUUCCCAGGUUCGCCUGGGAGCCGGACUUGCCCCUGUCGACGGUACGCU
GUCGGCAGCCGCUGCGGAGCCCUGGCGAGCUCCGGUUCGCUAGGGUGGAGCAAAUUCCGUAG
UGGGUGAACGUAGGUUCCGCAGCUAGCCGGUUUGGGUGGAGGAAAUUCCUGCCUGGCAUGGA
GGAGUUCGCUUCUCUAUGAGCUCCAUGGUUCGCCGUGGGGCUGCGAGCAAGCCGUAAGCUCG
CCCAGGUAGGAGCCCGAACUGGCUGGCUGUGGAGCCUGCGUUUACGGCACCGGAACCACUCA
CCGGUGGACGCUGUAGAGCCAGUGGGAGAGGGGUAGAAGAGUGGACUACCCCUGACGGGAAU
ACGGCACCCGUCGGUCUGGGUGAGCCUGGGGUGUUCUCU

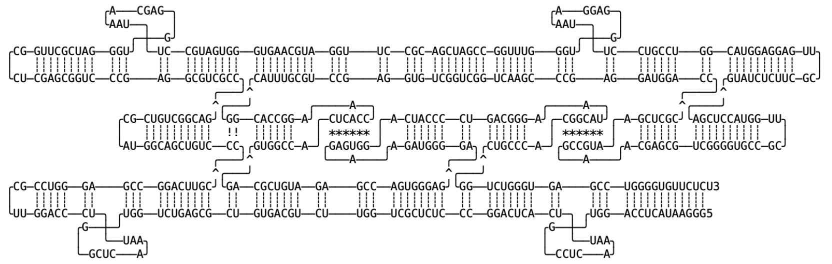

Figure S56: DNA template and primer sequences, RNA sequences and designed structure.

#### Figure S57. Sequence Design

##### Design: Strip-1x-alphaKL (G/AAA)

DNA Template Sequence:

AGAGAACACCCCAGGCTCACCCAGACCGACGGGTGCCGTATTCCCGTCAGGGGTAGTCCACT
CTTCTACCCCTCTCCCACTGGCTCTACAGCGTCCACCGGTGAGTGGTTCCGGTGCCGTAAAC
GCAGGCTCCACAGCCAGCCAGTTCGGGCTCCTACCTGGGCGAGCTTACGGCTTGCTCGCAGC
CCCACGGCGAACCATGGAGCTCATAGAGAAGCGAACTCCTCCATGCCAGGCAGGAACCCAAA
CCGGCTAGCTGCGGAATTTGTGCCACCTACGTTCACCCACTACGGAACCCTAGCGAACCGGA
GCTCGCCAGGGCTCCGCAGCGGCTGCCGACAGCGTACCGTCGACAGGGGCAAGTCCGGCTCC
CAGGCGAACCTGGGAACCAGACTCGCGACACTGCAGAATTTGCACCACCAGCGAGAGGGCCT
GAGTGAACCTGGAGTATTCCCTATAGTGAGTCGTATTAGAATGCCC

Forward Primer: GGGCATTCTAATACGACTCACTATA

Reverse Primer: AGAGAACACCCCAGGCTCAC

RNA Sequence:
GGGAAUACUCCAGGUUCACUCAGGCCCUCUCGCUGGUGGUGCAAAUUCUGCAGUGUCGCGAG
UCUGGUUCCCAGGUUCGCCUGGGAGCCGGACUUGCCCCUGUCGACGGUACGCUGUCGGCAGC
CGCUGCGGAGCCCUGGCGAGCUCCGGUUCGCUAGGGUUCCGUAGUGGGUGAACGUAGGUGGC
ACAAAUUCCGCAGCUAGCCGGUUUGGGUUCCUGCCUGGCAUGGAGGAGUUCGCUUCUCUAUG
AGCUCCAUGGUUCGCCGUGGGGCUGCGAGCAAGCCGUAAGCUCGCCCAGGUAGGAGCCCGAA
CUGGCUGGCUGUGGAGCCUGCGUUUACGGCACCGGAACCACUCACCGGUGGACGCUGUAGAG
CCAGUGGGAGAGGGGUAGAAGAGUGGACUACCCCUGACGGGAAUACGGCACCCGUCGGUCUG
GGUGAGCCUGGGGUGUUCUCU

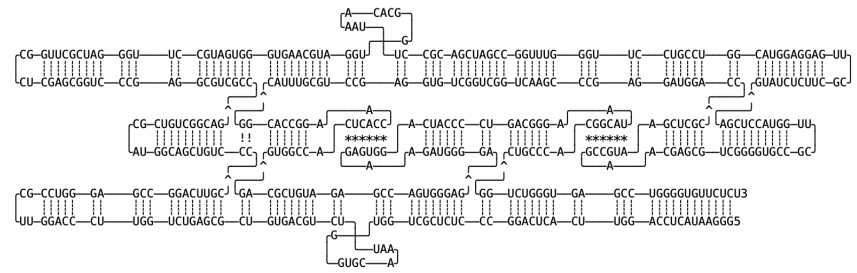

Figure S57: DNA template and primer sequences, RNA sequences and designed structure.

##### Figure S58. Sequence Design

Design: Strip-3x-alphaKL-SWWS (G/AAA)

DNA Template Sequence:

AGAGAACACCCCAGGCTCACCCAGACCGACGCGTGCCGTATTCGCGTCAGGGGTAGTCCACT
CTTCTACCCCTCTCCCACTGGCTCTACAGCGTCCGTCACTGAGTGGTTGTGACGCCGTAAAC
GCAGGCTCCACAGCCAGCCAGTTCGGGCTCCTACCTGGGCGAGCTTACGGCTTGCTCGCAGC
CCCACGGCGAACCATGGAGCTCATAGAGAAGCGAACTCCTCCATGCCAGGCAGGAATTTGAT
GCACCCAAACCGGCTAGCTGCGGAATTTGTAGCACCTACGTTCACCCACTAAGGAATTTGAA
CCACCCTAGCGAACCGGAGCTCGCCAGGGCTCCTCAGCGGCTGCCGACAGCGTACCGTCGAC
AGGGGCAAGTGAGGCCCATAGGCGAACCTATGGATTTGTTCCACCCCACTCGCGACACTGCA
GAATTTCTACCACCAGCGAGAGGGCCTGAGTGAATTTCATCCACCTGGAGTATTCCCTATAG
TGAGTCGTATTAGAATGCCC

Forward Primer: GGGCATTCTAATACGACTCACTATA

Reverse Primer: AGAGAACACCCCAGGCTCAC

RNA Sequence:

GGGAAUACUCCAGGUGGAUGAAAUUCACUCAGGCCCUCUCGCUGGUGGUAGAAAUUCUGCAG
UGUCGCGAGUGGGGUGGAACAAAUCCAUAGGUUCGCCUAUGGGCCUCACUUGCCCCUGUCGA
CGGUACGCUGUCGGCAGCCGCUGAGGAGCCCUGGCGAGCUCCGGUUCGCUAGGGUGGUUCAA
AUUCCUUAGUGGGUGAACGUAGGUGCUACAAAUUCCGCAGCUAGCCGGUUUGGGUGCAUCAA
AUUCCUGCCUGGCAUGGAGGAGUUCGCUUCUCUAUGAGCUCCAUGGUUCGCCGUGGGGCUGC
GAGCAAGCCGUAAGCUCGCCCAGGUAGGAGCCCGAACUGGCUGGCUGUGGAGCCUGCGUUUA
CGGCGUCACAACCACUCAGUGACGGACGCUGUAGAGCCAGUGGGAGAGGGGUAGAAGAGUGG
ACUACCCCUGACGCGAAUACGGCACGCGUCGGUCUGGGUGAGCCUGGGGUGUUCUCU

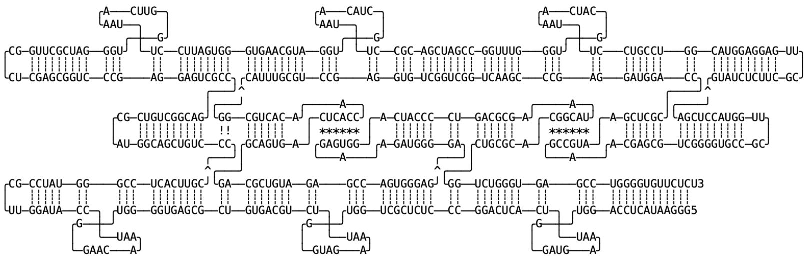

Weakest KL Energy (2GCs, not-stacked):
GAUG <-> CAUC : -1.50 [((((&)))))]
GUAG <-> CUAC : -1.50 [((((&)))))]
GAAC <-> GUUC : -1.40 [((((&)))))]

Figure S58: DNA template and primer sequences, RNA sequences and designed structure.

#### Figure S59. Sequence Design

##### Design: Strip-3x-alphaKL-WSSW (G/AAA)

DNA Template Sequence:

AGAGAACACCCCAGGCTCACCCAGACCGACGCGTGCCGTATTCGCGTCAGGGGTAGTCCACT
CTTCTACCCCTCTCCCACTGGCTCTACAGCGTCCGTCACTGAGTGGTTGTGACGCCGTAAAC
GCAGGCTCCACAGCCAGCCAGTTCGGGCTCCTACCTGGGCGAGCTTACGGCTTGCTCGCAGC
CCCACGGCGAACCATGGAGCTCATAGAGAAGCGAACTCCTCCATGCCAGGCAGGAATTTAGG
ACACCCAAACCGGCTAGCTGCGGAATTTTCCACACCTACGTTCACCCACTAAGGAATTTACC
TCACCCTAGCGAACCGGAGCTCGCCAGGGCTCCTCAGCGGCTGCCGACAGCGTACCGTCGAC
AGGGGCAAGTGAGGCCCATAGGCGAACCTATGGATTTAGGTCACCCCACTCGCGACACTGCA
GAATTTTGGACACCAGCGAGAGGGCCTGAGTGAATTTTCCTCACCTGGAGTATTCCCTATAG
TGAGTCGTATTAGAATGCCC

Forward Primer: GGGCATTCTAATACGACTCACTATA

Reverse Primer: AGAGAACACCCCAGGCTCAC

RNA Sequence:

GGGAAUACUCCAGGUGAGGAAAAUUCACUCAGGCCCUCUCGCUGGUGUCCAAAAUUCUGCAG
UGUCGCGAGUGGGGUGACCUAAAUCCAUAGGUUCGCCUAUGGGCCUCACUUGCCCCUGUCGA
CGGUACGCUGUCGGCAGCCGCUGAGGAGCCCUGGCGAGCUCCGGUUCGCUAGGGUGAGGUAA
AUUCCUUAGUGGGUGAACGUAGGUGUGGAAAAUUCCGCAGCUAGCCGGUUUGGGUGUCCUAA
AUUCCUGCCUGGCAUGGAGGAGUUCGCUUCUCUAUGAGCUCCAUGGUUCGCCGUGGGGCUGC
GAGCAAGCCGUAAGCUCGCCCAGGUAGGAGCCCGAACUGGCUGGCUGUGGAGCCUGCGUUUA
CGGCGUCACAACCACUCAGUGACGGACGCUGUAGAGCCAGUGGGAGAGGGGUAGAAGAGUGG
ACUACCCCUGACGCGAAUACGGCACGCGUCGGUCUGGGUGAGCCUGGGGUGUUCUCU

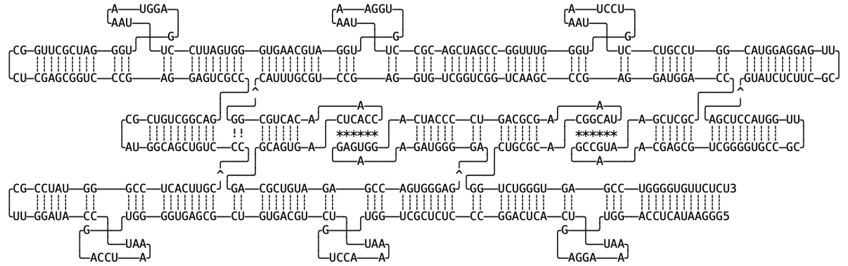

Weaker KL Energy (2GC IN MIDDLE):
AGGA <-> UCCU : -2.70 [((((&))))]
UGGA <-> UCCA : -2.70 [((((&))))]
AGGU <-> ACCU : -2.50 [((((&))))]

Figure S59: DNA template and primer sequences, RNA sequences and designed structure.

#### Figure S60. Sequence Design

##### Design: Square-Array-8x-alphaKL (G/AAA)

DNA Template Sequence:

GACCATAAGGCTCGACGAGGCGGAGTTTGGCTCCGTCGACCCAGCATAAGGGCTCGGAGCTG
CCGCAAGAGGCTCGGTGTGAGGGATTCACCGAATTGGCTGCACCTCTCGCGGTAGTTCAGCT
CCGAATTTGAGCCACCCTTATACCGGATCGACGGAATTTGCACCACCAAACCCCGTAGTTCC
TCGTCGAATTTCAGCCACCTTATAGTCGGAGATTAAGAATTTCCTCCACCGCTTGATGTAGT
TCGTCCCTGAATTTGTGCCACCGATTTCGCACCGAGCTTGAATTTGCTCCACCGGTAGTGGT
AGTTCGAATAGGAATTTGAGGCACCACGCCTTCCCTCTTGGCGTGGCTCCTATTCGCCACCA
CCGGCTCAAACTCGACGCGAAACCGGCTCAGGGACGCACCAAGCGGCTCTCAACCTCCTATA
GTGAGTCGTATTAGAATGCCC

Forward Primer: GGGCATTCTAATACGACTCACTATA

Reverse Primer: GACCATAAGGCTCGACGAGG

RNA Sequence:
GGAGGUUGAGAGCCGCUUGGUGCGUCCCUGAGCCGGUUUCGCGUCGAGUUUGAGCCGGUGGU
GGCGAAUAGGAGCCACGCCAAGAGGGAAGGCGUGGUGCCUCAAAUUCCUAUUCGAACUACCA
CUACCGGUGGAGCAAAUUCAAGCUCGGUGCGAAAUCGGUGGCACAAAUUCAGGGACGAACUA
CAUCAAGCGGUGGAGGAAAUUCUUAAUCUCCGACUAUAAGGUGGCUGAAAUUCGACGAGGAA
CUACGGGGUUUGGUGGUGCAAAUUCCGUCGAUCCGGUAUAAGGGUGGCUCAAAUUCGGAGCU
GAACUACCGCGAGAGGUGCAGCCAAUUCGGUGAAUCCCUCACACCGAGCCUCUUGCGGCAGC
UCCGAGCCCUUAUGCUGGGUCGACGGAGCCAAACUCCGCCUCGUCGAGCCUUAUGGUC

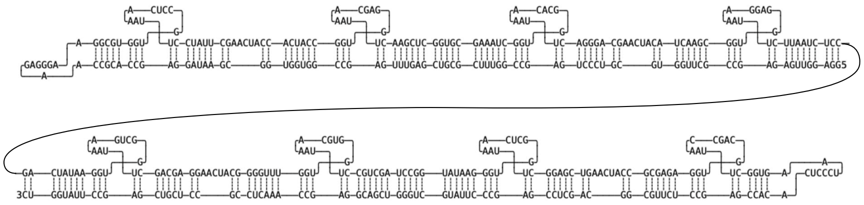

Figure S60: DNA template and primer sequences, RNA sequences and designed structure.

#### Figure S61. Sequence Design

##### Design: 2H-8x-alphaKL-wide (G/AAA)

dsDNA Template Sequence:

GACCAGACAGAATTTGCGACACCGAAGTTGGGGTCCCGAAGGACCGCCTGCGAACAGGCCCT
GAATTGGAATTTTCGCCACCCCACTGAACCCTTGTGGGGCTCCAATCCAGGCCAACCTCGGC
TCCGTCTAGTCGGCTGCGAGGCTCGTCACTGTGCCACACTGGGCTCACCCCTGGGTTCTTGG
GGTGAATTTCCTCCACCCAGTGCGGGTCCAATGTGAATTTGCTCCACCCTCGTTGTGCCTTC
GAGGGCTCACATTAGACGAGAGCAGGGCTCTACCAGCCGTGAGCATCGGCTCCATCGTATGC
CATAATGGGCTCCTCCCTGGCACATTGGGAGGAATTTGTGCCACCCATTACGGGCTGCCGAA
GCAGCGACGGCGAACCGTCCACACGATGGAATTTGCACCACCGATGCCCACGGCTAATAGAA
TTTGAGCCACCCTGCCCTCCACAGCGACGAATTTGAGGCACCTCGCAACCTATAGTGAGTCG
TATTAGAATGCCC

Forward Primer: GGGCATTCTAATACGACTCACTATAGG

Reverse Primer: GACCAGACAGAATTTGCGACAC

RNA Sequence:

GGUUGCGAGGUGCCUCAAAUUCGUCGCUGUGGAGGGCAGGGUGGCUCAAAUUCUAUUAGCCG
UGGGCAUCGGUGGUGCAAAUUCCAUCGUGUGGACGGUUCGCCGUCGCUGCUUCGGCAGCCCG
UAAUGGGUGGCACAAAUUCCUCCCAAUGUGCCAGGGAGGAGCCCAUUAUGGCAUACGAUGGA
GCCGAUGCUCACGGCUGGUAGAGCCCUGCUCUCGUCUAAUGUGAGCCCUCGAAGGCACAACG
AGGGUGGAGCAAAUUCACAUUGGACCCGCACUGGGUGGAGGAAAUUCACCCCAAGAACCCAG
GGGUGAGCCCAGUGUGGCACAGUGACGAGCCUCGCAGCCGACUAGACGGAGCCGAGGUUGGC
CUGGAUUGGAGCCCCACAAGGGUUCAGUGGGGUGGCGAAAAUUCCAAUUCAGGGCCUGUUCG
CAGGCGGUCCUUCGGGACCCCAACUUCGGUGUCGCAAAUUCUGUCUGGUC

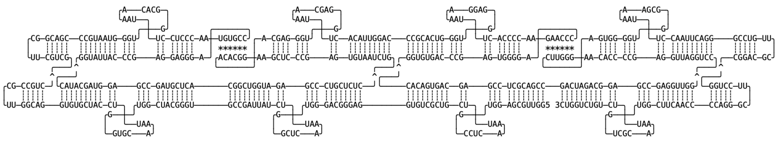

Figure S61: DNA template and primer sequences, RNA sequences and designed structure.

#### Figure S62. Sequence Design

##### Design: 2H-8x-alphaKL-narrow (G/AAA)

dsDNA Template Sequence:

GGGACCCAGCATTCGAAAATACGCACCGGGTGAATTTTGCCCACCGGCGCAGAATTTCCTCC
ACCAACGTCCATGATTCGTTGGCTCTGCACCGGCTCACCCAGTGCTACACCAAGGCTCTACC
TAGGCTCTCTACAATCCCATTTGCCCATCACATTTTCATGGTTAATATGACCCTCCTTGCAC
TTTTAGAAAGGCCACGACTACATGTTAACTGGCTCCTCTCCGGCTCAAATCCAGCGCATAGG
TGGCTCGTGCCTGGCTCCGAATTAAGTGCTTACTCGGAATTTGTGCCACCAGACACGAATTT
TCGCCACCACCCATGCCAGCGCGAACGCCGTAGCCCGAAGGCCAGCTAGATTTGAATTTGCG
ACACCGGAAAGGAATTTGCACCACCAGCTAACACGTAGTCGCGGTGGACAAATGGAATTGTA
AAGAATTTGAGGCACCTAGATAGAATTTGGCACACCTTGGCGTATGGATCCCTATAGTGAGT
CGTATTAGAATGCCC

Forward Primer: GGGCATTCTAATACGACTCACTATAGG

Reverse Primer: GGGACCCAGCATTCGAAAATAC

RNA Sequence:

GGGAUCCAUACGCCAAGGUGUGCCAAAUUCUAUCUAGGUGCCUCAAAUUCUUUACAAUUCCA
UUUGUCCACCGCGACUACGUGUUAGCUGGUGGUGCAAAUUCCUUUCCGGUGUCGCAAAUUCA
AAUCUAGCUGGCCUUCGGGCUACGGCGUUCGCGCUGGCAUGGGUGGUGGCGAAAAUUCGUGU
CUGGUGGCACAAAUUCCGAGUAAGCACUUAAUUCGGAGCCAGGCACGAGCCACCUAUGCGCU
GGAUUUGAGCCGGAGAGGAGCCAGUUAACAUGUAGUCGUGGCCUUUCUAAAAGUGCAAGGAG
GGUCAUAUUAACCAUGAAAAUGUGAUGGGCAAAUGGGAUUGUAGAGAGCCUAGGUAGAGCCU
UGGUGUAGCACUGGGUGAGCCGGUGCAGAGCCAACGAAUCAUGGACGUUGGUGGAGGAAAUU
CUGCGCCGGUGGGCAAAAUUCACCCGGUGCGUAUUUUCGAAUGCUGGGUCCC

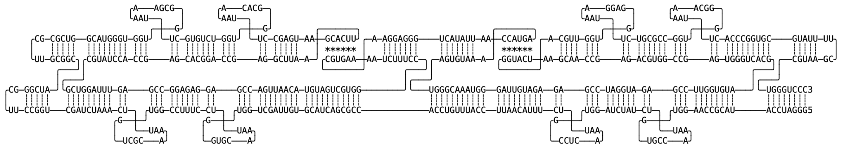

Figure S62: DNA template and primer sequences, RNA sequences and designed structure.
